# Evaluation of the anti-inflammatory effects of synthesised tanshinone I and isotanshinone I analogues in zebrafish

**DOI:** 10.1101/773309

**Authors:** Matthew J. Foulkes, Katherine M. Henry, Stephen A. Renshaw, Simon Jones

**Affiliations:** Department of Chemistry, The University of Sheffield, Dainton Building, Brook Hill, Sheffield, S3 7HF, UK; The Bateson Centre, The University of Sheffield, Sheffield, S10 2TN, UK; Department of Infection, Immunity & Cardiovascular Disease, The University of Sheffield, Sheffield, S10 2RX, UK

## Abstract

During inflammation, dysregulated neutrophil behaviour can play a major role in chronic inflammatory diseases such as chronic obstructive pulmonary disease, for which current treatments are generally ineffective. Recently, tanshinones have shown promising antiinflammatory effects by targeting neutrophils *in vivo*, yet are still an underexplored general group of compounds. Here, an existing six step synthetic route was optimised and used to prepare a small family of substituted tanshinone and isomeric isotanshinone analogues, together with the synthesis of other structurally similar molecules. Evaluation of these using a transgenic zebrafish model of inflammation revealed that many of these compounds exhibit promising anti-inflammatory effects *in vivo*. Several compounds affect neutrophil recruitment and/or resolution of neutrophilic inflammation, and broad structure-activity relationships were constructed. In particular, the methoxy-substituted tanshinone **39** specifically accelerates resolution of inflammation without affecting organism host defence, making this a particularly attractive candidate for potential pro-resolution therapeutics. On the other hand, β-lapachones exhibit effects on neutrophil recruitment yet not on resolution. Notable differences in toxicity profiles between compound classes were also observed.

## Introduction

Tanshinones are a group of aromatic diterpenoid compounds comprising four fused rings; two of these form a naphthalene or tetrahydronaphthalene moiety, the third ring usually bears an *ortho*- (or *para*-) quinone, and the fourth ring is a furan or dihydrofuran. Tanshinone I (TI) **1** is one of the main known tanshinones found naturally in the plant *Salvia miltiorrhiza* (also known as Chinese red sage or danshen), whilst others include tanshinone IIA (TIIA) **2**, and the dihydro counterparts, dihydrotanshinone I **3** and cryptotanshinone **4** (Figure 1). The plant has been traditionally used as a remedy in Chinese herbal medicine since as early as the first century AD, mainly for treatment of cardiovascular disease.^1^ Isolation of tanshinones from the plant usually requires a series of extractions starting from the dried roots immersed in alcoholic solution, followed by chromatographic separation,^2–5^ although this procedure is usually only suitable for obtaining milligram quantities for biological testing.

**Figure 1.**
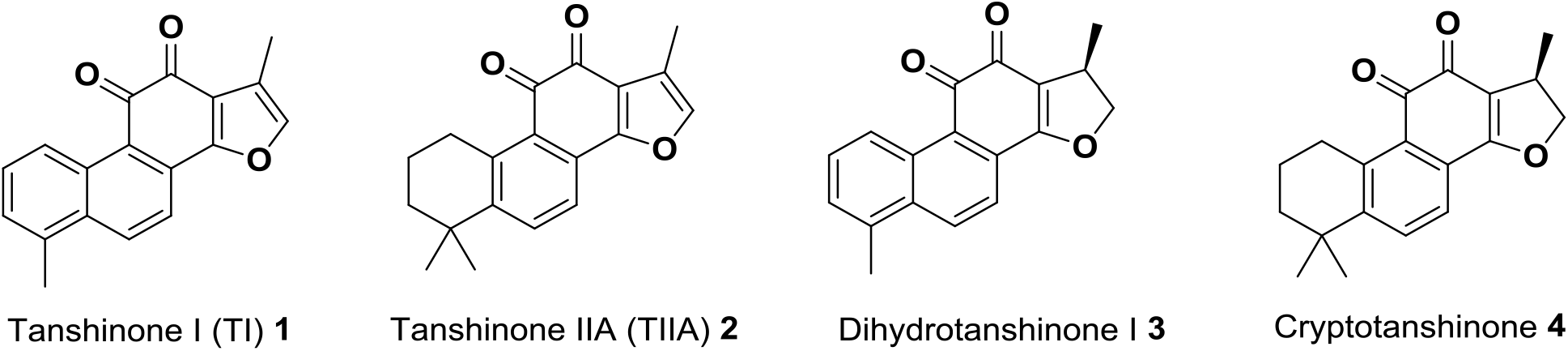
Structures of some naturally occurring tanshinones **1-4**.

Isolated tanshinones exhibit various biological activities both *in vitro* and *in vivo*. In particular, TI **1** has been shown to display numerous effects including anti-inflammatory effects,^6–9^ antimicrobial activity,^10,11^ cytotoxicity towards cancer cells,^12–15^ neuroprotective effects,^16^ benefits for learning and memory enhancement,^17^ antioxidant activity,^18,19^ and the ability to induce cell apoptosis.^12,20,21^ In most cases however, the mechanism of action is poorly understood, and there is often limited or no knowledge of structure-activity relationships.

β-Lapachone **5** and nor-β-lapachone **6** are molecules with a high degree of structural similarity to tanshinones (Figure 2). β-Lapachone **5** is a naturally-occurring *ortho*-quinone compound obtained from the bark of the lapacho tree *Tabebuia avellanedae* in Brazil, a plant used for centuries as a traditional medicine for various treatments, including as an analgesic, an antiinflammatory agent, an antineoplastic agent, and a diuretic.^22,23^ In more recent studies, β-lapachone **5** has exhibited anti-inflammatory effects,^24–29^ as well as various anti-cancer activities,^30–33^ anti-angiogenesis activity,^34,35^ antimicrobial effects,^36–38^ and wound healing properties.^39^ In addition, both β-lapachone **5** and TIIA **2** have been found to act as inhibitors of NAD(P)H quinone oxidoreductase 1 (NQO1), leading to an increase in cellular NAD(+).^40–42^ This recent finding connecting the two structures, alongside the clear structural similarities, may suggest that these two classes of compounds could exhibit common biological activities *in vivo*. In contrast, reports of previous studies involving nor-β-lapachone **6** are scarce. Although nor-β-lapachone **6** is considered an anti-cancer drug candidate,^43,44^ at this time there appear to be no studies involving nor-β-lapachone **6** in relation to inflammation, neutrophils or zebrafish, perhaps because β-lapachone **5** is a naturally occurring compound, whereas nor-β-lapachone **6** is not.

**Figure 2.**
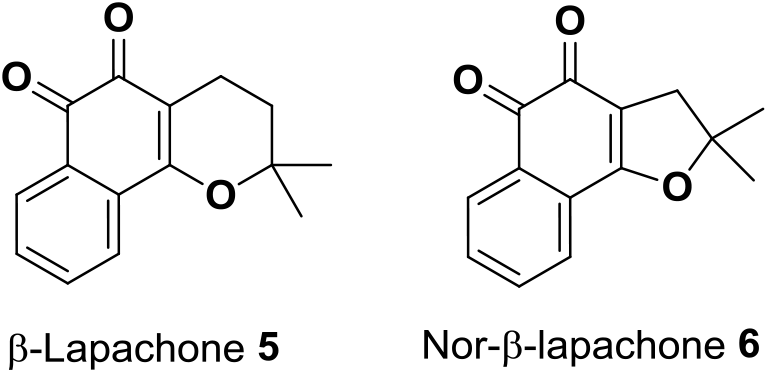
Structures of β-lapachone **5** and nor-β-lapachone **6**

Neutrophils, the most abundant white blood cell in humans, play a vital role in the inflammatory response. Their recruitment to the site of pathogen invasion or injury is essential for elimination of invading pathogens by various mechanisms.^45–49^ Strict regulation of neutrophil function is required: whilst these processes are required for effective host defence and response, they also need to be limited to avoid persistent inflammation.^50^ Subsequent timely neutrophil clearance from this site, as part of the resolution of inflammation, is also necessary for healthy recovery.^51,52^ Failure of this process can result in retention of persistent inflammatory neutrophils, which can have damaging effects on the body, playing a key role in diseases such as chronic obstructive pulmonary disease (COPD),^53,54^ asthma,^55,56^ and rheumatoid arthritis.^57,58^ Such conditions are increasingly prevalent diseases; according to the World Health Organisation, COPD is the fourth highest cause of global mortality, claiming over three million lives in 2015 (5% of all global deaths), a figure projected to increase further in future years.^59^ Current treatments for unresolved neutrophilic inflammation are non-specific, poorly effective, and can exhibit many undesired side-effects;^60,61^ thus there is an unmet need for new neutrophil-specific clinical treatments.

The zebrafish (*Danio rerio*) is an excellent *in vivo* model for the study of inflammation and neutrophil biology,^62^ and has been utilised effectively in a number of recent studies in the discovery of previously unidentified compounds with anti-inflammatory properties *in vivo*.^63–66^ Screening in a transgenic zebrafish inflammation model identified TIIA **2** as a compound which significantly accelerated resolution of neutrophilic inflammation *in vivo*, without affecting initial neutrophil recruitment or total neutrophil number.^64^ The related molecule cryptotanshinone **4** displayed broadly similar anti-inflammatory effects, although this compound also decreased initial neutrophil recruitment. However, no other tanshinones have been investigated in this manner, meaning that structure-activity relationships both within and between different tanshinone classes (TI and TIIA) in this model are largely unexplored. This work investigates construction of structure-activity relationships for such molecules in the zebrafish inflammation model, through analogue synthesis and testing.

## Results and Discussion

### Chemical synthesis of TI analogues and isomers

Synthesis of TI analogues utilised a route similar to that of Jiao and co-workers,^67^ but with important modifications (Scheme 1). Commercially available 5-bromovanillin **7** was first converted to the hydroquinone **8** *via* Baeyer-Villiger oxidation and subsequent ester hydrolysis, followed by Fe(III)-mediated oxidation to the corresponding benzoquinone **9**.^68^ Both transformations were performed on a multi-gram scale (up to 25 grams) in high yield, with a lack of chromatographic purification representing a more time- and cost-effective large-scale isolation of the desired benzoquinone **9**. This material was treated with various substituted carboxylic acids in a radical decarboxylative coupling. Using 3-(2-methylphenyl)propionic acid **10**, employing optimal literature conditions provided the desired alkylation product **11**, but in a significantly lower yield of around 20%, in comparison to yields of 65-79% previously reported.^67^ By carrying the reaction out in the dark, slightly modifying the work-up by adding ice and reducing the volume of aqueous sodium hydroxide used, and adding the oxidant as a more dilute solution, each separately led to an increased yield. Interestingly, reducing the number of equivalents of carboxylic acid **10** from 2 to 1.2 had no detrimental effect on the yield. Combining these changes resulted in a 42% yield of alkylation product **11** after chromatographic purification. In each case, formation of a small quantity (< 10%) of doubly-alkylated product was observed, yet none of the alternative monoalkylated product (alkylated adjacent to the methoxide group) was seen, which was consistent with previous reports, and likely a directing effect due to the bromide substituent.^67,69^ Additional optimisation studies were also carried out. Using the unsubstituted 3-phenylpropionic acid **12** as a model substrate (to eliminate the possibility of an additional substituent having any electronic or steric impact on the reaction), a number of reaction parameters were investigated using a Design of Experiments (DoE) approach (further information provided in the SI). Using the Custom Design function in JMP software (JMP®, Version 12. SAS Institute Inc., Cary, NC, 1989-2007), the single most important factor was determined to be the method of aqueous persulfate addition. Adding the persulfate solution *via* glass dropping funnel gave a consistently elevated yield compared to addition *via* metal syringe, an observation noted elsewhere with aqueous persulfate solutions,^70^ thought to be a result of the metal syringe accelerating the rate of decomposition of the oxidant solution.^71^ Using silver(I) phosphate gave higher yields than other silver(I) salts investigated, whilst heating at 85 °C and increasing the number of equivalents of silver(I) salt both gave mild increases in product yield, with all other changes identified as having minimal or no effect.

**Scheme 1.**
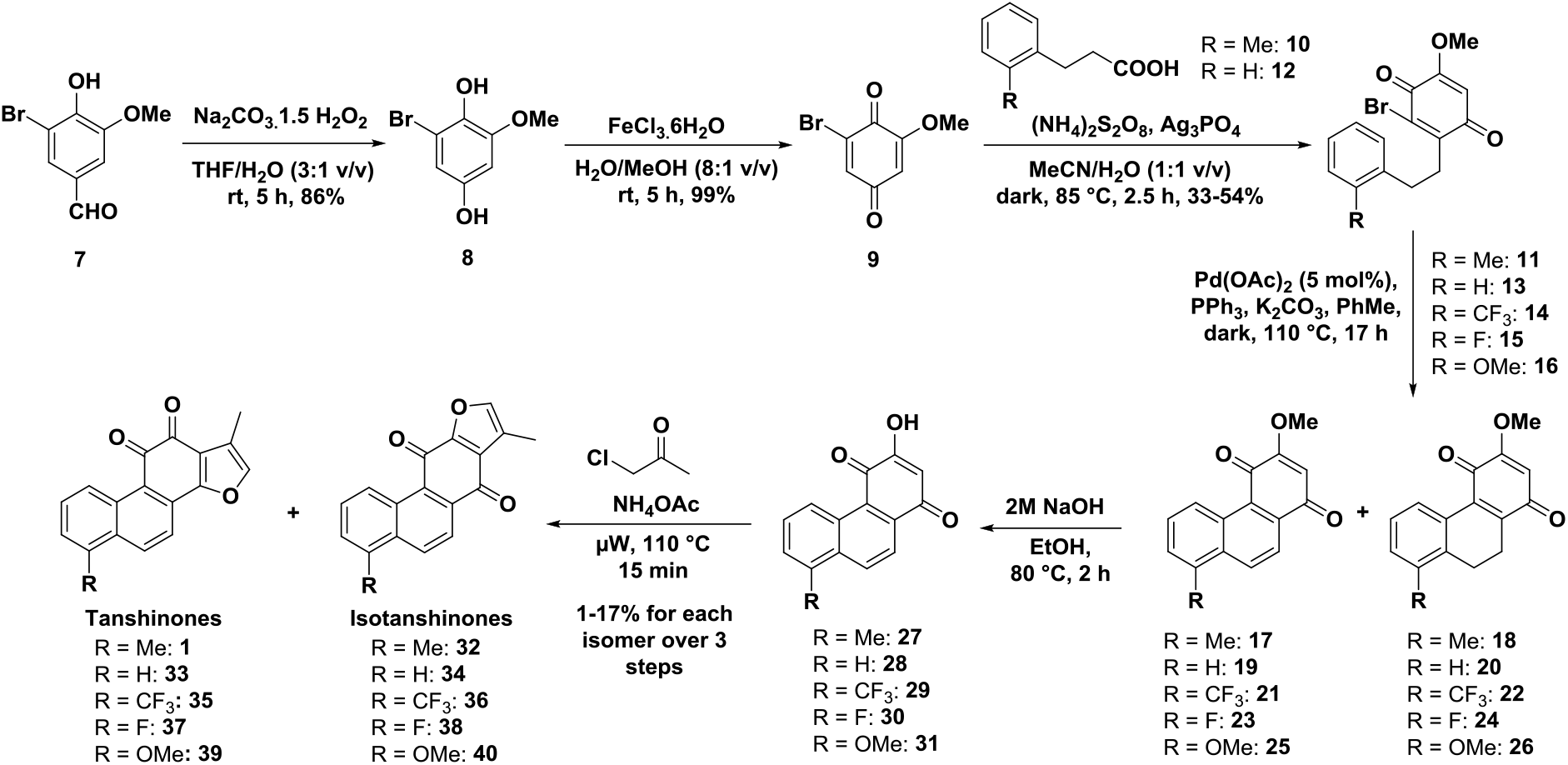
Synthesis of tanshinones and isotanshinones.

The optimised conditions were applied to the reaction for other substituted carboxylic acids. Products bearing methyl **11**, hydrogen **13**, trifluoromethyl **14** and fluoro **15** substituents were all formed in consistently moderate yields of 38-54%. For the methoxy substituent, some benzoquinone starting material **9** remained, which could not be separated from the desired alkylation product **16** by either chromatographic purification on silica, or recrystallisation. Analysis of the ^1^H NMR spectrum of the mixture indicated a 60:40 ratio of the alkylation product **16** and the starting material **9**, corresponding to an approximate 33% overall yield of desired product **16**.

Employing the conditions for the intramolecular Heck reaction used in the literature for the methyl-substituted bromide **11**,^67^ only gave the desired cyclisation product in a maximum of 40% yield after purification, as a mixture of aromatised **17** and non-aromatised **18** compounds. This was unacceptably low given the high quantities of palladium catalyst used (45 mol%).

Attempted optimisation using the unsubstituted variant **13** as a model system, by reducing the amount of palladium catalyst from 45 mol% to 10 mol%, gave a correspondingly low yield. Changing the ligand from triphenylphosphine to tri(*o*-tolyl)phosphine had no obvious effect, neither did the addition of either 1,4-benzoquinone or palladium on carbon as a co-oxidant. However, when the reaction was performed at a three-fold lower concentration (again using 10 mol% catalyst), the yield increased to 68% after purification. At a five-fold lower concentration (approximately 0.01 M), the yield further increased to 74%, and this was conserved when the catalyst loading was reduced to 5 mol%, with a 73% yield obtained. These results thus suggested that this particular intramolecular reaction was especially sensitive to concentration effects. It should be noted that throughout these studies, the ratio of aromatised **19** and non-aromatised products **20** varied considerably, possibly due to variations in the quality of the nitrogen atmosphere or the palladium source used. However, this was not considered an issue, as any non-aromatised product **20** was converted to the fully aromatised product in subsequent steps.

The improved Heck reaction conditions were used to prepare the corresponding analogues **17-26**. Whilst mass returns were good for the methyl- and hydrogen-substituted variants **17-18** and **19-20**, heteroatom-substituted compounds were lower. In the specific case of the methoxy-substituted compounds, purified starting material **16** was predominately isolated initially, suggesting that the palladium catalyst underwent faster oxidative addition with the simple benzoquinone **9** than with the alkylated compound **16**. After subjecting this recovered material to the Heck conditions for a second time, the desired products **25-26** were successfully produced although also with a low mass return.

Attempted demethylation of the mixture of inseparable methyl ethers **17-18** to form the alcohol **27** using boron trichloride (1.2 or 3 equivalents) or tetra-*n*-butylammonium iodide (TBAI) returned only starting material. However, heating at reflux in a 2 M sodium hydroxide solution of ethanol led to complete demethylation with concomitant aromatisation, successfully yielding the desired alcohol **27**. This avoided the need for additional oxygen gas to be supplied to the reaction, as previously utilised in the literature,^67^ and worked well for all analogues **27-30**, and for the methoxy compound **31**, where only the single desired demethylation product was observed.

In a change of conditions from the literature procedure, reaction of the alcohol **27** with chloroacetone as the solvent with 1 equivalent of ammonium acetate resulted in isolation of the desired TI **1**, together with the isomeric isotanshinone I (iso-TI) **32**.^67,72–75^ The two isomers were each isolated in 12% yield over three steps, presumably formed *via* tautomeric intermediates. This reaction worked consistently for each of the different analogues to yield tanshinones **1, 33, 35, 37, 39** and isotanshinones **32, 34, 36, 38, 40**. Yields were higher for the electron-neutral methyl- and hydrogen-substituted compounds, and were somewhat lower for the heteroatom-substituted variants. Only a few examples of isotanshinones are known, including iso-TI **32** and isotanshinone IIA **41**,^72,73,75,76^ and related isotanshinones based on the parent compounds dihydrotanshinone I **3** and cryptotanshinone I **4**, which have recently been investigated as anti-cancer agents.^77,78^ These all occur naturally in *Salvia miltiorrhiza*, although isotanshinone I **32** and isotanshinone IIA **41** have been produced synthetically.^72,73^ However as a class of compounds, isotanshinones are largely underexplored, both in terms of synthesis and biological studies. Thus, this synthesis provided efficient access to a range of both substituted tanshinone I analogues and isotanshinones of interest.

In order to investigate the importance of the *ortho*-quinone moiety of TI **1** for its biological activity (as a functional group common to all major tanshinones), a ‘masked’ analogue lacking this functionality was desired. Synthesis of a dimethyl acetal analogue of TI **1** was readily accomplished in a two-step procedure (Scheme 2) by reduction of the *ortho*-quinone **1** with sodium borohydride, followed by quickly treating the diol **42** with 2,2-dimethoxypropane and acid giving acetal **43** in 34% yield over two steps. This was stable in solution, in air, and to column chromatography, in contrast to the diol **42** that spontaneously decomposed back to TI **1** in solution and on exposure to air, matching similar observations made previously for TIIA **2**.79

**Scheme 2.**
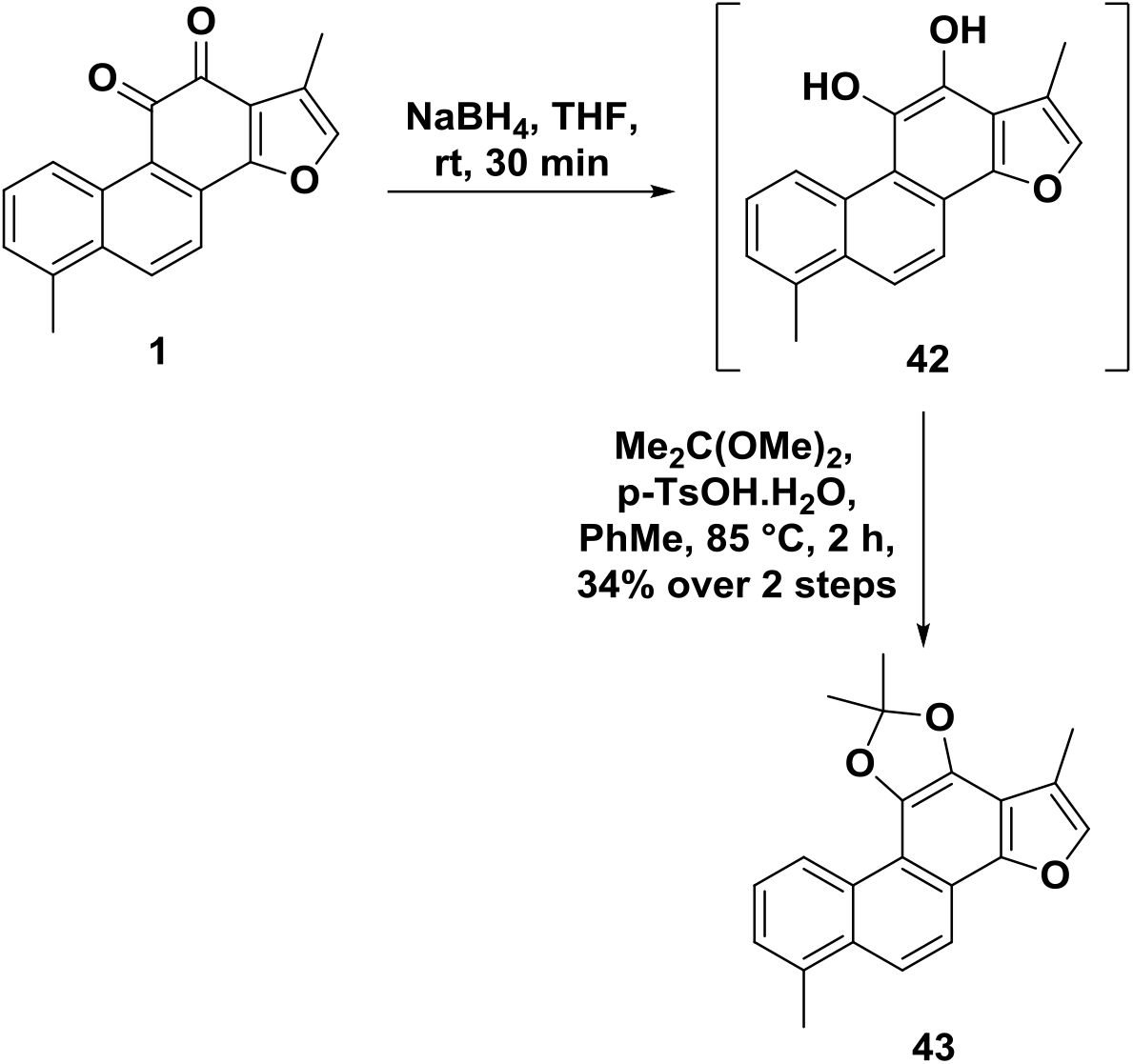
Synthesis of the TI-acetal **43**.

### Chemical synthesis of lapachols and β-lapachones

Structurally related β-lapachones were synthesised readily from commercially sourced lapachol **44** (Scheme 3). β-Lapachone **5** was synthesised in a single step and in high yield from treatment of lapachol **44** with concentrated sulfuric acid.

**Scheme 3.**
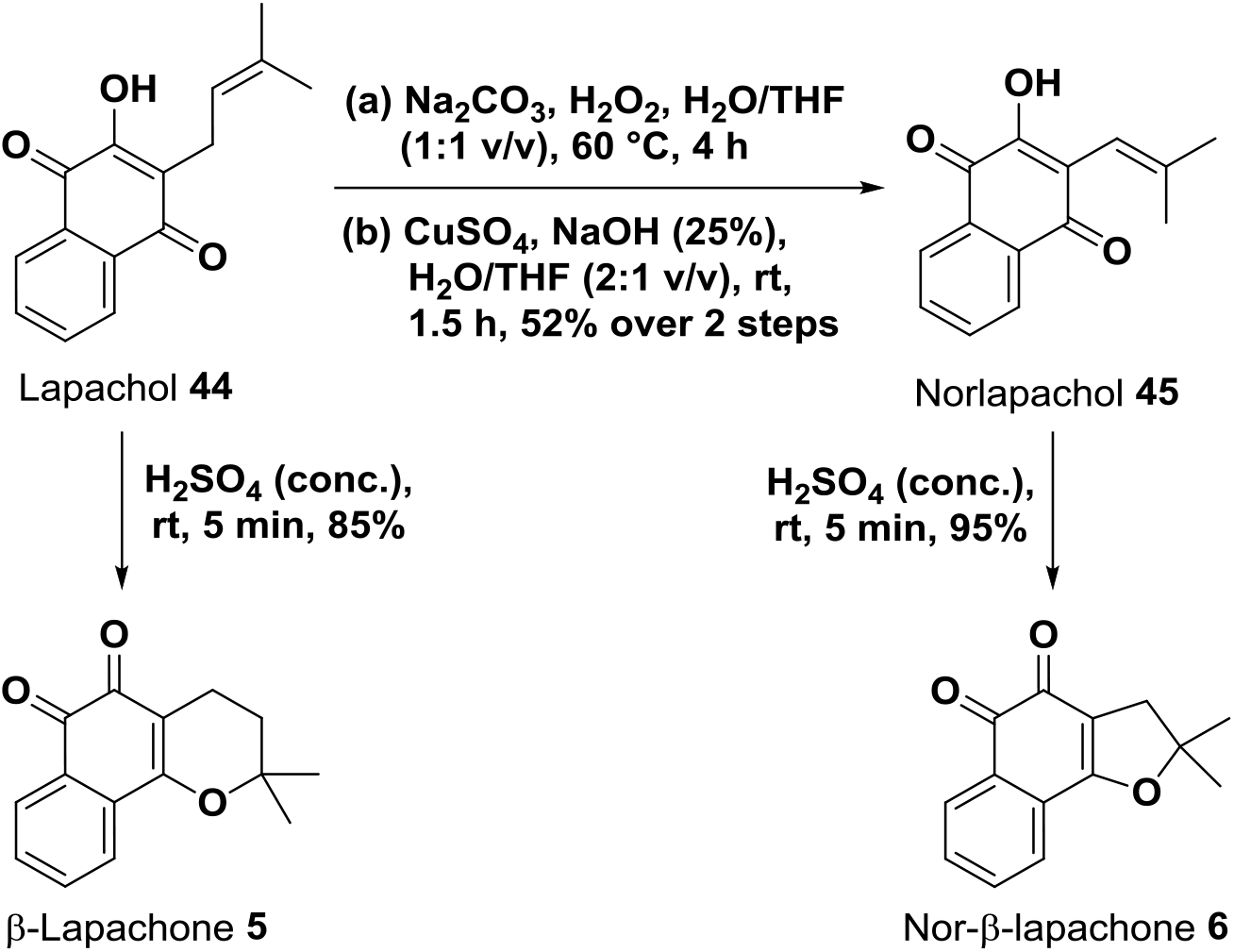
Synthesis of norlapachol **45** and β-lapachones **5, 6**.

Norlapachol **45** was also produced from lapachol, *via* a Hooker oxidation, in an unoptimized yield of 52% after recrystallisation. Finally, nor-β-lapachone **6** was synthesised from its corresponding precursor norlapachol **45**, again *via* acid-mediated cyclisation, in a near-quantitative yield of 95% after simple purification by trituration.

### Evaluation of the *in vivo* anti-inflammatory effects of TI and TIIA

To explore the anti-inflammatory effects of the different tanshinone sub-classes, TI **1** and TIIA **2** were first evaluated in a transgenic zebrafish model of inflammation (Figure 3), as previously described.^64,65^ For neutrophil recruitment experiments, the commercially available compound SP600125 (SP) was used as the positive control, as a selective and cell-permeable inhibitor of c-Jun N-terminal kinase (JNK), which prevents activation of various inflammatory genes.^80,81^

**Figure 3.**
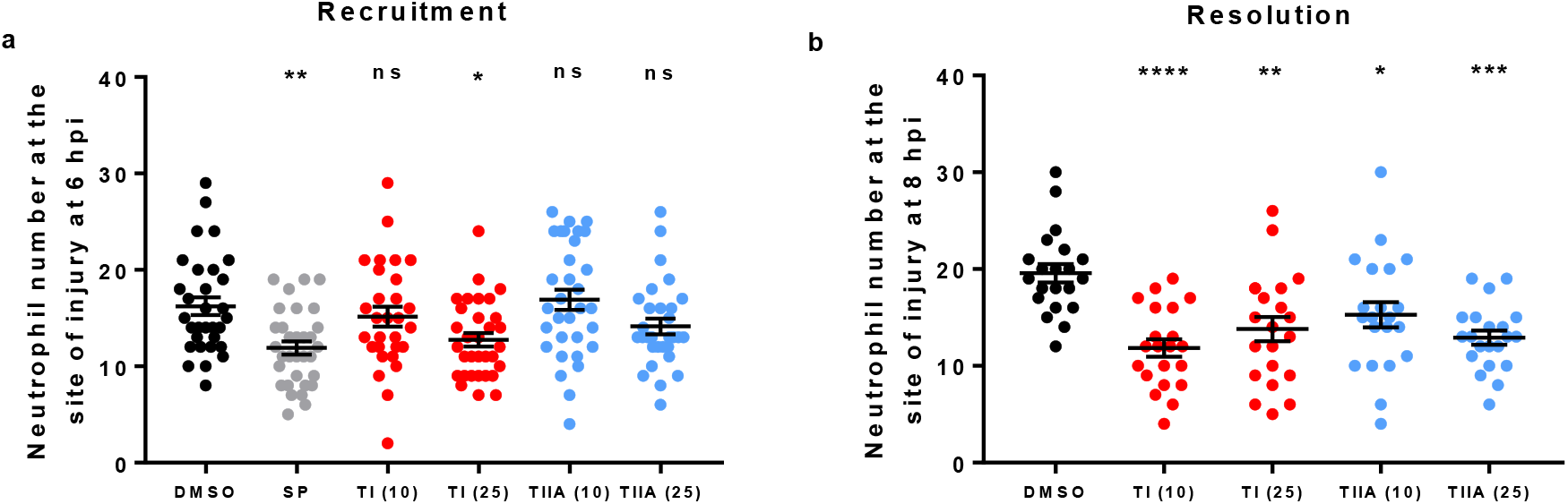
Effects of TI **1** and TIIA **2** on **a**) initial neutrophil recruitment and **b**) resolution of neutrophilic inflammation in a zebrafish inflammation model. For **a**), tailfin transection was performed on zebrafish larvae (3 dpf) which were subjected to compound treatments immediately: DMSO (0.5%), SP600125 (30 μM) and TI **1** and TIIA **2** (10, 25 μM as indicated in brackets). Neutrophil number at the site of injury was assessed at 6 hpi. For **b**), tailfin transection was performed on zebrafish larvae (3 dpf), and at 4 hpi, good responders were subjected to compound treatments: DMSO (0.5%), TI **1** and TIIA **2** (10 μM, 25 μM as indicated in brackets). Neutrophil number at the site of injury was assessed at 8 hpi. For both graphs, data shown as mean ± SEM; *n* = 21-32 larvae per group from 3-4 independent experiments per graph. * *P* < 0.05, ** *P* < 0.01, *** *P* < 0.001, **** *P* < 0.0001, compared to the DMSO vehicle control (one-way ANOVA with Dunnett’s multiple comparison post-test).

Investigation of the effects of the tanshinones on initial recruitment of neutrophils to the site of tailfin injury (Figure 3a) revealed that TI **1** (25 μM) reduced the number of neutrophils recruited at 6 hpi, although had no effect at 10 μM. TIIA **2**, however, did not significantly affect neutrophil recruitment at either 10 or 25 μM; these results were in agreement with previous findings for TIIA **2** using this model.^64^ For experiments investigating resolution of neutrophilic inflammation *in vivo* (Figure 3b), TIIA **2** (25 μM) was used as the positive control compound, as this has previously been shown to exhibit highly significant pro-resolution activity in this zebrafish model of inflammation.^64^ Both TI **1** and TIIA **2** accelerated resolution of neutrophilic inflammation, at 10 and 25 μM.

### Evaluation of the effects of TI analogues and isomers on neutrophil recruitment and resolution *in vivo*

The synthesised TI analogues and isomers **1**, **32**-**40**, **43** were evaluated for any effects on neutrophil recruitment in the same zebrafish inflammation model (Figure 4). As for previous neutrophil recruitment experiments, SP600125 was used as the positive control compound. In comparison to the DMSO vehicle control, TI **1** significantly reduced the number of neutrophils recruited to the site of injury in all three datasets (Figures 4b-d). Generally, treatment with iso-TI **32** also resulted in a significant decrease in the number of neutrophils recruited (Figures 4b-d); although this was not the case in one set (Figure 4c), this likely reflected natural variability, arising either in relation to this particular compound, or possibly in relation to the nature of the *in vivo* experiments performed. For the tanshinone analogues **33, 35, 37, 39** derived from the parent TI structure **1** (Figures 4b-c, data shown in red), replacement of the methyl group in the 6-position of TI **1** with a hydrogen atom (**33**) resulted in the loss of any effect on neutrophil recruitment to the injury site. When the 6-position comprised either a trifluoromethyl (**35**), fluorine (**37**) or methoxy (**39**) substituent, there was also no statistical difference in neutrophil recruitment compared to the DMSO control.

**Figure 4.**
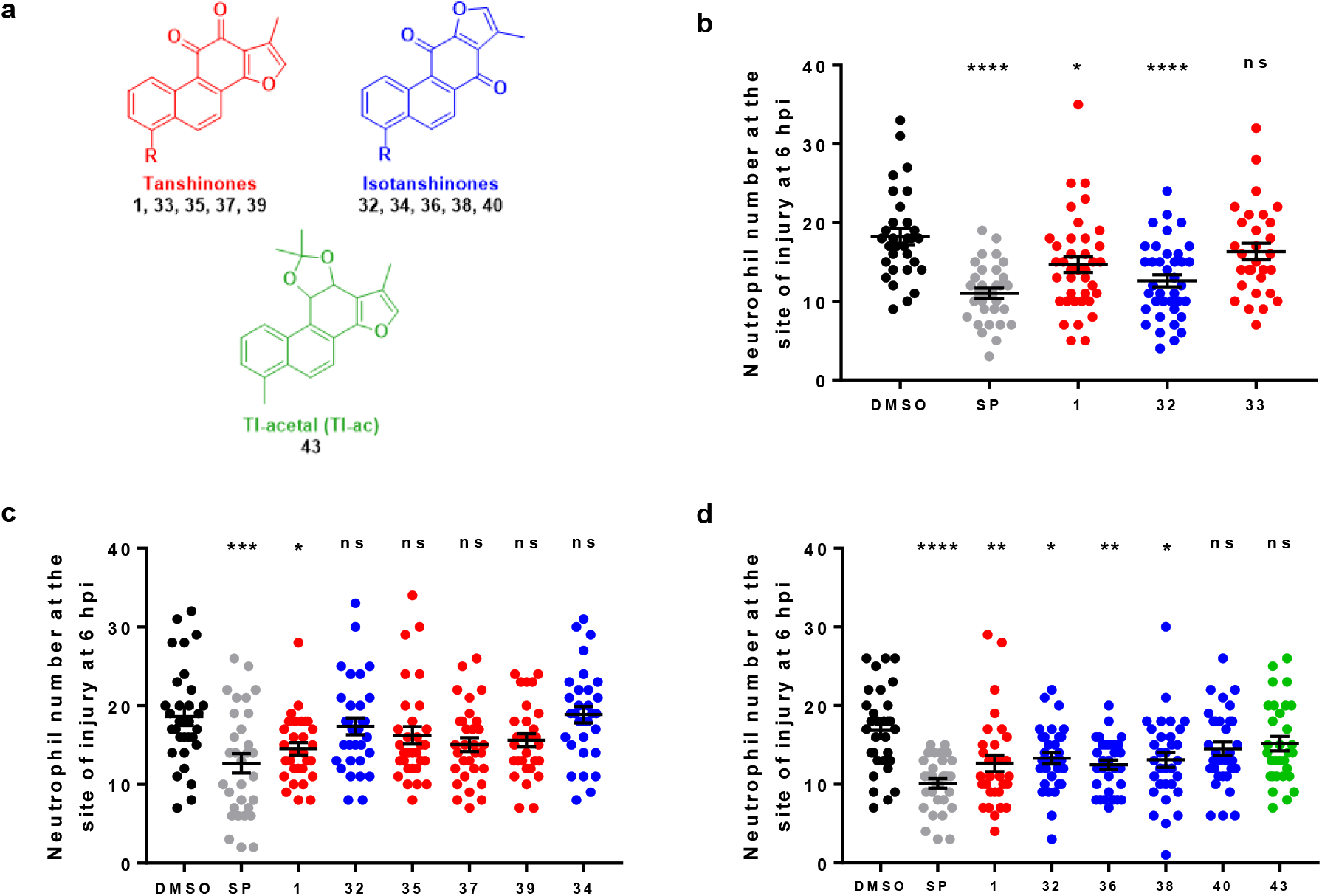
Effects of TI analogues and isomers on neutrophil recruitment. **a**) Structures of evaluated tanshinones (red), isotanshinones (blue) and the TI-acetal (green). **b**)-**d**) Tailfin transection was performed on zebrafish larvae (3 dpf) which were subjected to compound treatments immediately: DMSO (0.5%), SP600125 (30 μM) and compounds **1**, **32-40**, **43** (25 μM). Neutrophil number at the site of injury was assessed at 6 hpi. Data shown as mean ± SEM; *n* = 30-39 larvae from 4 independent experiments per graph. * *P* < 0.05, ** *P* < 0.01, *** *P* < 0.001, **** *P* < 0.0001, ns not significant, in comparison to the DMSO vehicle control (oneway ANOVA with Dunnett’s multiple comparison post-test).

For the isotanshinones **34, 36, 38, 40** (Figures 4c-d, relevant data depicted in blue), replacement of the 4-methyl group with a hydrogen atom (**34**) again resulted in the loss of any effect on neutrophil recruitment to the site of injury. However, replacement of this methyl group with either a trifluoromethyl group (**36**) or a fluorine atom (**38**) resulted in significantly fewer recruited neutrophils in each case, although the methoxy-substituted analogue **40** had no significant effect on recruitment. The TI-acetal **43** also had no significant effect on neutrophil recruitment to the site of injury (Figure 4d).

When the data for the various analogues are considered together, the results for compounds with each functional group show some consistency between tanshinones and isotanshinones. Compounds with a methyl substituent have a clear effect in reducing neutrophil recruitment to the site of injury; analogues bearing trifluoromethyl or fluorine substituents appear to exhibit some effect or trend. Removal of any functionality at this position, or replacement with a methoxy substituent, leads to a complete loss of biological effect. This suggests that the presence of a substituent in the modified position is required for activity, but the nature of the substituent may not be overly important.

The synthesised TI analogues and isomers **1, 32-40, 43** were then evaluated in a similar zebrafish model of inflammation resolution (Figure 5). For resolution experiments, TIIA **2** was again used as the positive control compound (Figures 5b-d). Although this did not quite show the expected effect in one set of compounds evaluated (Figure 5c), the results were very close to statistical significance (*P* = 0.052), and the compound otherwise performed as expected (Figures 5b and d).

**Figure 5.**
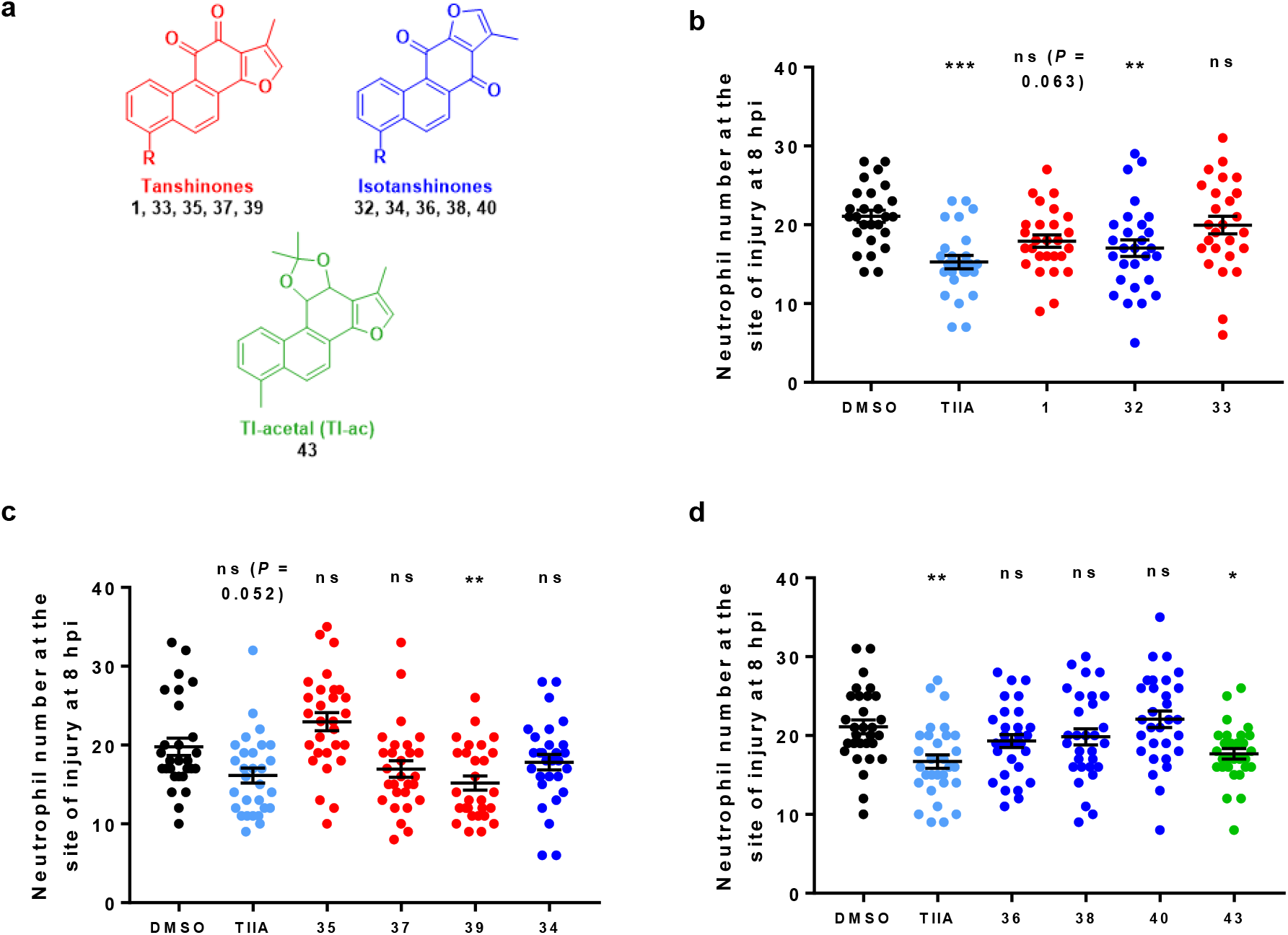
Effects of TI analogues and isomers on resolution of neutrophilic inflammation. **a**) Structures of tanshinones (red), isotanshinones (blue) and TI-acetal (green). **b**)-**d**) Tailfin transection was performed on zebrafish larvae (3 dpf), and at 4 hpi, good responders were subjected to compound treatments: DMSO (0.5%), TIIA **2** (25 μM) and compounds **1**, **32-40**, **43** (25 μM). Neutrophil number at the site of injury was assessed at 8 hpi. Data shown as mean ± SEM; *n* = 26-30 larvae per group from 4 independent experiments per graph. * *P* < 0.05, ** *P* < 0.01, *** *P* < 0.001, ns not significant, in comparison to the DMSO control (one-way ANOVA with Dunnett’s multiple comparison post-test).

Considering the *ortho*-quinone tanshinone compounds **1, 33, 35, 37, 39** first (Figures 5b-c, data shown in red), in larvae treated with TI **1**, there was a trend towards lower neutrophil numbers at the site of injury, compared to the DMSO-treated larvae (*P* = 0.063). Replacement of the methyl group in the 6-position (**1**) with a hydrogen atom (**33**) resulted in no effect on resolution *in vivo*. Related compounds containing a trifluoromethyl group (**35**) or fluorine atom (**37**) also showed no effect. Interestingly, for the 6-methoxy substituted tanshinone **39**, there was a clear reduction in neutrophil number at the site of injury at 8 hpi, in comparison to the DMSO control-treated larvae, indicating that this compound accelerates resolution of inflammation.

This compound **39** was of particular interest as it promoted resolution of neutrophilic inflammation but did not affect initial neutrophil recruitment, making this compound a more promising candidate for further investigation of specific pro-resolution treatments which do not impair host-defence.

For the isotanshinones **32, 34, 36, 38, 40**, (Figures 5b-d, data shown in blue), iso-TI **32** significantly accelerated neutrophilic inflammation resolution, whilst the unsubstituted compound **34** again had no effect. The remaining isotanshinones **36, 38, 40** had no effect on inflammation resolution, irrespective of the functional group in the 4-position.

Treatment with the TI-acetal **43** resulted in a reduction in neutrophil number at the site of injury (Figure 5d). One possible explanation is that the acetal **43** was (partially) hydrolysed and metabolised back to the parent TI **1** *in vivo*, which then exerted a pro-resolution effect as seen previously (Figure 3b). Thus, for inflammation resolution, the patterns for each functional group are less clear between tanshinones and isotanshinones. The tanshinones and isotanshinones containing a methyl group in the 6- or 4-position, respectively, generally show some effect (especially when considered with earlier experiments, see Figure 3b), and removal of functionality at this position results in loss of any effect. Compounds substituted with a trifluoromethyl group or a fluorine atom show no significant effect on resolution. When the substituent is a methoxy group, the tanshinone **39** has a pro-resolution effect, but the isotanshinone **40** has no effect. As for recruitment, this suggests that a substituent in this position is required for activity. However, in contrast to neutrophil recruitment, the nature of the substituent seems to be more important for pro-resolution activity.

Overall, some of these compounds affect both recruitment and resolution, whilst some affect neither, and others have an effect on only one of these inflammatory processes. This is not particularly surprising, as molecules which have an effect on neutrophil recruitment likely interact with a different molecular target to those which affect resolution of neutrophilic inflammation. Both of the parent compounds TI **1** and iso-TI **32** decreased neutrophil recruitment to the site of injury and accelerated inflammation resolution, whilst both of the unsubstituted molecules **33** and **34** had no effect on either process. The fluorine-containing tanshinones **35** and **37** also had no significant effect on either process. However, the analogous isotanshinones **36** and **38** both resulted in decreased neutrophil recruitment to the injury site yet had no effect on resolution of neutrophilic inflammation. The methoxy-substituted tanshinone **39** exhibited no effect on neutrophil recruitment, yet significantly promoted inflammation resolution, which was analogous to the findings observed for TIIA **2**.

This suggests that this particular compound may be an interesting candidate to focus on to investigate compounds which specifically target inflammation resolution. The same activity profile in these two types of experiment was also seen for the TI acetal **43**, although as discussed previously, this may have been due to the formation of products arising from acetal hydrolysis and metabolism. In contrast to the corresponding tanshinone, the methoxy-substituted isotanshinone **40** had no effect on either the recruitment or resolution stages of neutrophilic inflammation.

Comparison of the fluorine-containing compounds with their non-fluorine-containing analogues could provide insights into how *in vivo* metabolism may affect biological activity. Whereas TI **1** had an effect on both neutrophil recruitment and resolution, the trifluoromethyl-substituted variant **35** did not significantly affect either process. Similarly, iso-TI **32** also affected both recruitment and resolution, whilst the analogous compound **36** reduced neutrophil recruitment to the site of injury yet had no effect on resolution of neutrophilic inflammation. This could suggest that *in vivo* C-H bond metabolism (or some other factor relating to C-F bond substitution) is important for acceleration of inflammation resolution, but not for neutrophil recruitment. The unsubstituted tanshinone **33** did not affect either recruitment or resolution, and neither did the fluorine-substituted compound **37**. Meanwhile, the unsubstituted isotanshinone **34** also had no effect on neutrophil numbers for either recruitment or resolution, whilst treatment with the analogous fluorine-substituted isotanshinone **38** resulted in decreased neutrophil recruitment but did not affect resolution. In contrast to the results for the trifluoromethyl variants, this may suggest that replacement of a hydrogen atom with a fluorine atom here has more effect on neutrophil recruitment than on resolution of neutrophilic inflammation.

For all experiments, the general health of treated zebrafish larvae was also observed. At concentrations of 25 μM, none of the evaluated tanshinones and isotanshinones resulted in any larval toxicity; larvae were generally healthy in all treatments, as determined by general body shape, tail shape, presence of circulation, and heartbeat.

### Evaluation of the effects of lapachols and β-lapachones on neutrophil recruitment and resolution *in vivo*

Norlapachol **45**, lapachol **44**, β-lapachone **5**, and nor-β-lapachone **6** were evaluated in neutrophil recruitment experiments (Figure 6). To explore possible dose-response effects, norlapachol **45** was tested at 0.1, 1, and 10 μM. A higher dose of 25 μM was toxic to the larvae. The remaining three compounds were all evaluated at 1 μM; higher doses were also toxic. This stood in contrast to the use of the various tanshinones at 25 μM previously, in which larvae were healthy. These observations suggest that the structural differences between these compounds have important consequences on their toxicity profiles *in vivo*. This could be due to differences in molecular target interactions, differences in larval penetration, and/or differences in neutrophil penetration, possibly due to interactions with relevant drug transporter proteins.^82^ TI **1** (25 μM) was used as the positive control, based on the previous findings and to provide a synthetic control compound. Norlapachol **45** reduced the number of neutrophils recruited to the site of injury at both 10 μM and 0.1 μM (Figure 6a). Given the structural differences between norlapachol **45** and tanshinones, norlapachol **45** may have worked in a different way to tanshinones such as TI **1**, possibly by acting on a different protein target. Recruitment was not significantly reduced at 1 μM. The difference between the effects observed at 1 μM and 0.1 μM may reflect the natural variability in this type of *in vivo* experiment, despite all efforts to control for this as far as reasonably practicable.

**Figure 6.**
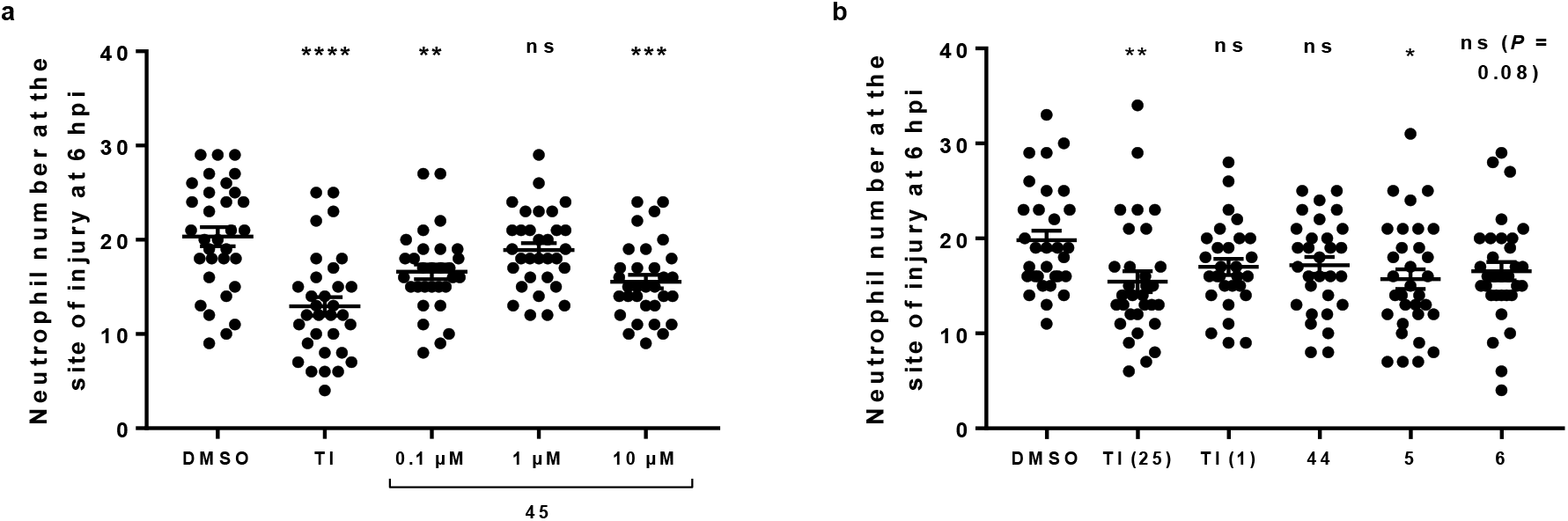
Effects of **a**) norlapachol **45** and **b**) lapachol **44**, β-lapachone **5** and nor-β-lapachone **6** on neutrophil recruitment. Tailfin transection was performed on zebrafish larvae (3 dpf) which were subjected to compound treatments immediately: DMSO (0.5%), TI **1** (25 μM; also 1 μM for **b** only, as indicated in brackets), norlapachol **45** (**a**; 0.1, 1, 10 μM as indicated) and lapachol **44**, β-lapachone **5** and nor-β-lapachone **6** (**b**; 1 μM). Neutrophil number at the site of injury was assessed at 6 hpi. Data shown as mean ± SEM; *n* = 29-32 larvae from 4 independent experiments per graph. * *P* < 0.05, ** *P* < 0.01, *** *P* < 0.001, **** *P* < 0.0001, ns not significant, in comparison to the DMSO control (one-way ANOVA with Dunnett’s multiple comparison post-test).

The remaining three compounds were evaluated in the same way but at 1 μM (Figure 6b). TI **1** was used at both 25 μM (as a known positive control) and 1 μM, to allow comparison of the effects of this compound to the other compounds tested at the same dosage. β-Lapachone **5** led to a significant reduction in the number of neutrophils recruited to the site of injury. However, neither nor-β-lapachone **6** nor lapachol **44** reduced neutrophil recruitment, nor did TI **1** at this concentration. No signs of toxicity were observed amongst the larvae of any of the treatment groups.

These results were particularly interesting, as β-lapachone **5** and nor-β-lapachone **6** are both *ortho*-quinone compounds which are structurally similar to TI **1**, and both led to reduced neutrophil numbers at the injury site (significant for β-lapachone **5** only), as observed with TI **1** at a 25 μM concentration. However, TI **1** showed no effect when used at a concentration of 1 μM, suggesting that β-lapachone **5** (and possibly also nor-β-lapachone **6**) may have been more efficacious than TI **1** *in vivo*. This would clearly require further investigation, but is of interest given the toxicity issues observed with the β-lapachones yet not with TI **1** in this study.

The same four compounds were evaluated for any effects on resolution of neutrophilic inflammation *in vivo* (Figure 7). Norlapachol **45** was tested at a range of different concentrations from 0.1 to 25 μM, as the compound was not toxic to larvae at 25 μM in these experiments. The higher threshold for toxicity in resolution experiments may be attributed to the shorter treatment time with compound (4 hours instead of 6 hours in recruitment studies) and the fact that the tissue has been allowed to heal somewhat before application of compounds. Lapachol **44**, β-lapachone **5** and nor-β-lapachone **6** were each tested at 1 μM, as higher doses were toxic. TIIA **2** (25 μM) was used as a positive control, as in previous resolution experiments. Norlapachol **45** had no effect on the number of neutrophils at 0.1 and 1 μM (Figure 7a). However, a reduced neutrophil number at the site of injury was observed at 10 μM, and a highly significant reduction was seen at a concentration of 25 μM, although some of the larvae did show slow or absent circulation. Overall, the resolution data for norlapachol **45** show a good dose-response relationship. Finally, the remaining compounds lapachol **44**, β-lapachone **5**, and nor-β-lapachone **6** were analysed in similar inflammation resolution experiments, at 1 μM (Figure 7b). In these experiments, TIIA **2** was used at 25 μM (as the positive control) and 1 μM, again to compare the effects of this compound to the other compounds tested at the same concentration. However, none of these experimental compounds had any effect on resolution of neutrophilic inflammation.

**Figure 7.**
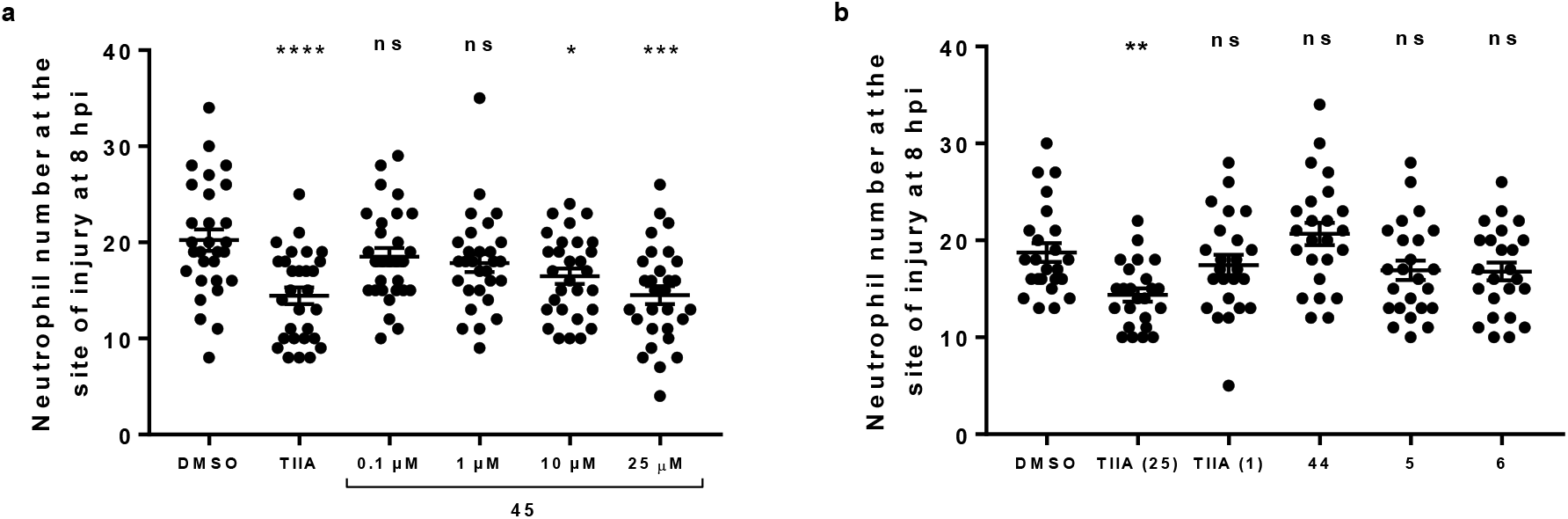
Effects of **a**) norlapachol **45** and **b**) lapachol **44**, β-lapachone **5** and nor-β-lapachone **6** on resolution of neutrophilic inflammation. Tailfin transection was performed on zebrafish larvae (3 dpf), and at 4 hpi, good responders were subjected to compound treatments: DMSO (0.5%), TIIA **2** (25 μM; also 1 μM for **b** only, as indicated in brackets), norlapachol **45** (**a**; 0.1, 1, 10, 25 μM as indicated) and lapachol **44**, β-lapachone **5** and nor-β-lapachone **6** (**b**; 1 μM). Neutrophil number at the site of injury was assessed at 8 hpi. Data shown as mean ± SEM; *n* = 24-29 larvae from 3-4 independent experiments per graph. * *P* < 0.05, ** *P* < 0.01, *** *P* < 0.001, **** *P* < 0.0001, ns not significant, in comparison to the DMSO control (one-way ANOVA with Dunnett’s multiple comparison post-test).

Overall, norlapachol **45** affected both neutrophil recruitment to the site of injury and resolution at higher concentrations, yet at lower concentrations, only initial neutrophil recruitment was significantly affected. Lapachol **44** exhibited no observable effect on either recruitment or resolution, and neither did nor-β-lapachone **6**, although the number of neutrophils recruited to the site of injury appeared to be lower for treatment with this compound. At a low concentration, β-lapachone **5**, affected only recruitment of neutrophils, and not resolution of neutrophilic inflammation. This may suggest a specific function of this compound (at this concentration) in earlier stages of the inflammatory response.

## Conclusions

A small library of analogues and isomers of TI **1** was synthesised from commercially available starting materials using an optimised route, which allowed access to novel tanshinone derivatives as well as isotanshinones, an underexplored class of compounds. Synthesis of an acetal variant of TI **1**, as well as structurally similar β-lapachones, was also undertaken. Evaluation of these compounds in a transgenic zebrafish model of inflammation revealed that various related molecules exhibited anti-inflammatory effects *in vivo*. Small changes in the molecular structures of these compounds resulted in varying effects on different stages of the inflammatory response, allowing for some broad structural activity relationships to be constructed. Three compounds – β-lapachone **5** and isotanshinones **36** and **38** – resulted in decreased neutrophil recruitment to the site of injury, but did not affect resolution of neutrophilic inflammation, whilst one tanshinone **39** accelerated inflammation resolution yet did not affect initial neutrophil recruitment. This compound is considered a particularly attractive candidate for potential pro-resolution therapeutics, as it exhibited an antiinflammatory effect without affecting organism host defence. The acetal **43** also affected resolution but not neutrophil recruitment. Three compounds – tanshinones **1** and **35**, and isotanshinone **32** – reduced initial neutrophil recruitment and accelerated inflammation resolution. This study also revealed differences in the toxicity profiles between tanshinones and the closely related β-lapachones, exemplifying the utility of using zebrafish phenotypic screening in identifying safety issues with potential drug treatments early in the drug discovery and development process. Overall, this work has enabled an increased insight into the anti-inflammatory activities of various substituted tanshinones, isotanshinones, and structurally related molecules, working towards identification of candidates for clinical treatment of dysregulated inflammation which act specifically on inflammatory neutrophils.

## Experimental Section

### Chemical synthesis

#### General reagents, materials and methods

All chemicals used were purchased from commercial suppliers and were used as received without further purification. Melting points were determined using a Gallenkamp melting point apparatus equipped with a thermometer. IR spectroscopy was performed on a PerkinElmer FT-IR Spectrum 65 or Spectrum 100 spectrometer, using either NaCl discs or a Universal diamond ATR. ^1^H, ^13^C and ^19^F NMR experiments were run on either a Bruker Avance 400 or Bruker Avance III HD 500 spectrometer at 298 K. Chemical shifts (δ) are reported in parts per million (ppm) relative to the deuterated lock solvent as an internal standard, where s = singlet, d = doublet, t = triplet, q = quartet, m = multiplet, br s = broad singlet, br d = broad doublet, br t = broad triplet, dd = doublet of doublets, ddd = double doublet of doublets, dt = doublet of triplets, td = triplet of doublets. All coupling constants are reported in hertz, Hz. Mass spectrometry was carried out on either an Agilent Technologies 6530 or 7200 spectrometer, using either electron impact (EI) or electrospray ionisation (ESI). TLC was performed on Merck silica gel 60 F254 aluminium-backed plates and visualised using ultraviolet light followed by staining with potassium permanganate dip. Column chromatography was carried out using silica gel obtained from VWR Chemicals, particle size 40-63 μm. Reactions using microwave conditions were carried out on a CEM Corporation Discover S-class microwave synthesiser, at pressure ≤ 17 bar and power ≤ 200 W. Synthesis of all intermediates for the tested compounds are provided in the supplementary information (SI).

#### General representative procedure A for reactions with chloroacetone to form tanshinones 1, 33, 35, 37, 39 and isotanshinones 32, 34, 36, 38, 40

A mixture of the alcohol **27-31** (100 mg, 0.42 mmol), ammonium acetate (32 mg, 0.42 mmol, 4 mmol g^-1^) and chloroacetone (10 mL g^-1^, 12 mmol) was heated in a sealed tube at 110 °C for 15 minutes under microwave conditions. The solution was cooled to room temperature, diluted with water (400 mL g^-1^) and extracted with DCM (3 x 400 mL g^-1^). The organic extracts were combined, dried (MgSO_4_), filtered and concentrated *in vacuo* to give the crude mixture which was purified by flash column chromatography (silica gel, DCM) to give the tanshinones **1, 33, 35, 37, 39** and isotanshinones **32, 34, 36, 38, 40**.

#### 1,6-Dimethylphenanthro[1,2-*b*]furan-10,11-dione, TI 1 and 4,8-dimethylphenanthro[3,2-*b*]furan-7,11-dione, iso-TI 32

General procedure **A** was followed, using the alcohol **27** (100 mg, 0.42 mmol) to give the dione **1** (21 mg, 12% over 3 steps) as a dark red solid; mp 232-234 °C (lit.^83^ 229-230 °C); *δ*_H_(400 MHz; CDCl_3_) 9.23 (1 H, d, *J* 8.8, ArC*H*), 8.27 (1 H, d, *J* 8.8, ArC*H*), 7.77 (1 H, d, *J* 8.8, ArC*H*), 7.54 (1 H, dd, *J* 8.8, 7.1, ArC*H*), 7.34 (1 H, d, *J* 7.1, ArC*H*), 7.30 (1 H, s, OC*H*), 2.68 (3 H, s, C*H*_3_), 2.30 (3 H, d, *J* 0.8, C*H*_3_); *δ*_C_(100 MHz; CDCl_3_) 183.4 (*C*=O), 175.6 (*C*=O), 161.2 (Ar*C*), 142.0 (Ar*C*H), 135.2 (Ar*C*), 133.6 (Ar*C*), 132.9 (Ar*C*H), 132.7 (Ar*C*), 130.6 (Ar*C*H), 129.6 (Ar*C*), 128.3 (Ar*C*H), 124.8 (Ar*C*H), 123.1 (Ar*C*), 121.8 (Ar*C*), 120.5 (Ar*C*), 118.7 (Ar*C*H), 19.9 (*C*H_3_), 8.8 (*C*H_3_); *m/z* (ESI^+^) 299 (10%, M+Na^+^), 277 (100, M+H^+^). All data were in general agreement with the literature.^67,74,83^ Also obtained was the dione **32** (22 mg, 12% over 3 steps) as an orange solid; mp 218-220 °C (lit.^72^ 219-220 °C); ν_max_(ATR)/cm^-1^ 1655 (C=O), 1588 (C=O), 1533 (C=C); *δ*_H_(400 MHz; CDCl_3_) 9.66 (1 H, d, *J* 8.9, ArC*H*), 8.40 (1 H, dd, *J* 8.9, 0.8, ArC*H*), 8.32 (1 H, d, *J* 8.9, ArC*H*), 7.63 (1 H, dd, *J* 8.9, 7.0, ArC*H*), 7.54-7.52 (1 H, m, OC*H*), 7.48 (1 H, dt, *J* 7.0, 0.8, ArC*H*), 2.76 (3 H, s, ArC*H*_3_), 2.41 (3 H, d, *J* 1.2, C*H*_3_); *δ*_C_(101 MHz; CDCl_3_) 182.2 (*C*=O), 177.2 (*C*=O), 153.9 (Ar*C*), 145.2 (Ar*C*H), 136.0 (Ar*C*), 134.7 (Ar*C*), 133.5 (Ar*C*), 131.2 (Ar*C*), 130.9 (Ar*C*H), 129.8 (Ar*C*H), 129.3 (Ar*C*H), 127.2 (Ar*C*), 126.5 (Ar*C*), 126.2 (Ar*C*H), 122.1 (Ar*C*H), 120.8 (Ar*C*), 20.0 (*C*H_3_), 8.7 (*C*H_3_); *m/z* (ESI^+^) 299 (25%, M+Na^+^), 277 (100, M+H^+^). No ^13^C NMR spectroscopy data were reported in the literature; all other data were in general agreement with the literature.^72,75,83^

#### 1-Methylphenanthro[1,2-*b*]furan-10,11-dione 33 and 8-methylphenanthro[3,2-*b*]furan-7,11-dione 34

General procedure **A** was followed, using the alcohol **28** (100 mg, 0.44 mmol) to give the dione **33** (24 mg, 15% over 3 steps) as a dark red solid; mp 229-233 °C (lit.^67^ 226-229 °C); ν_max_(NaCl discs)/cm^-1^ 2922 (C-H), 2852 (C-H), 1669 (C=O), 1593 (C=C); *δ*_H_(400 MHz; CDCl_3_) 9.42 (1 H, d, *J* 8.7, ArC*H*), 8.12 (1 H, d, *J* 8.7, ArC*H*), 7.83 (2 H, br d, *J* 8.7, 2 x ArC*H*), 7.71 (1 H, br t, *J* 7.7, ArC*H*), 7.55 (1 H, t, *J* 7.7, ArC*H*), 7.33 (1 H, s, ArC*H*), 2.32 (3 H, s, C*H*_3_). No IR spectroscopy data were reported in the literature. ^1^H NMR data were in broad agreement with the literature,^67^ although precise chemical shift values were slightly shifted due to the different solvent used for analysis. Also obtained was the dione **34** (28 mg, 17% over 3 steps) as an orange solid; mp 199-202 °C; ν_max_(NaCl discs)/cm^-1^ 2921 (C-H), 1766 (C=O), 1664 (C=O), 1536 (C=C); *δ*_H_(400 MHz; CDCl_3_) 9.77 (1 H, d, *J* 8.5, ArC*H*), 8.28 (1 H, d, *J* 8.5, ArC*H*), 8.18 (1 H, d, *J* 8.5, ArC*H*), 7.90 (1 H, d, *J* 8.5, ArC*H*), 7.78-7.72 (1 H, m, ArC*H*), 7.68-7.61 (1 H, m, ArC*H*), 7.53 (1 H, s, ArC*H*), 2.40 (3 H, s, C*H*_3_); *δ*_C_(101 MHz; CDCl_3_) 182.2 (*C*=O), 177.2 (*C*=O), 153.7 (Ar*C*), 145.3 (Ar*C*H), 136.7 (Ar*C*), 135.1 (Ar*C*H), 134.0 (Ar*C*), 130.9 (Ar*C*), 130.1 (Ar*C*H), 128.7 (Ar*C*H), 128.5 (Ar*C*H), 128.0 (Ar*C*H), 127.0 (Ar*C*), 126.5 (Ar*C*), 122.3 (Ar*C*H), 120.9 (Ar*C*), 8.7 (*C*H_3_); *m/z* (EI^+^) 262.0622 (100%, M^+^ C_17_H_10_O_3_ requires 262.0624), 234 (42), 206 (40), 176 (36), 151 (15), 126 (8), 87 (7), 76 (8).

#### 1-Methyl-6-(trifluoromethyl)phenanthro[1,2-*b*]furan-10,11-dione 35 and 8-methyl-4-(trifluoromethyl)phenanthro[3,2-*b*]furan-7,11-dione 36

General procedure **A** was followed, using the unpurified alcohol **29** (435 mg) to give the dione **35** (6 mg, 1% over 3 steps) as a dark red solid; mp 204-208 °C; ν_max_(NaCl discs)/cm^-1^ 1674 (C=O), 1550 (C=C); *δ*_H_(400 MHz; CDCl_3_) 9.65 (1 H, d, *J* 8.9, ArC*H*), 8.50 (1 H, d, *J* 8.9, ArC*H*), 7.98 (1 H, d, *J* 8.9, ArC*H*), 7.94 (1 H, d, *J* 7.2 ArC*H*), 7.75 (1 H, br t, *J* 8.1, ArC*H*), 7.39 (1 H, br d, *J* 1.1, OC*H*), 2.33 (3 H, d, *J* 1.1, C*H*_3_); *δ*_C_(100 MHz; CDCl_3_) 183.3(*C*=O), 175.2 (*C*=O), 160.2 (Ar*C*), 142.8 (Ar*C*H), 133.0 (Ar*C*), 132.8 (q, *J*_C-F_ 3.0, Ar*C*H), 131.0 (Ar*C*H), 130.5 (Ar*C*), 129.8 (Ar*C*), 129.0 (Ar*C*H), 127.0 (q, *J*_C-F_ 30.3, Ar*C*), 126.0 (q, *J*_C-F_ 5.9, Ar*C*H), 124.2 (q, *J*_C-F_ 273.9, *C*F_3_), 123.1 (Ar*C*), 122.1 (Ar*C*), 121.2 (Ar*C*), 120.8 (Ar*C*H), 8.8 (*C*H_3_); *δ*_F_(377 MHz; CDCl_3_) −58.8; *m/z* (ESI^+^) 353 (18%, M+Na^+^), 331.0580 (100, M+H^+^ C_18_H_10_O_3_F_3_ requires 331.0577). Also obtained was the dione **36** (20 mg, 2% over 3 steps) as a dark brown solid; mp 183-186 °C; ν_max_(NaCl discs)/cm^-1^ 1661 (C=O), 1601 (C=O), 1536 (C=C); *δ*_H_(400 MHz; CDCl_3_) 10.07 (1 H, d, *J* 9.0, ArC*H*), 8.58 (1 H, d, *J* 9.0, ArC*H*), 8.45 (1 H, d, *J* 9.0, ArC*H*), 8.05 (1 H, d, *J* 7.1 ArC*H*), 7.79 (1 H, dd, *J* 9.0, 7.1, ArC*H*), 7.57 (1 H, q, *J* 1.1, OC*H*), 2.42 (3 H, d, *J* 1.1, C*H*_3_); *δ*_C_(100 MHz; CDCl_3_) 181.4 (*C*=O), 176.6 (*C*=O), 153.5 (Ar*C*), 145.7 (Ar*C*H), 134.0 (Ar*C*), 132.4 (Ar*C*H), 132.2 (Ar*C*), 131.6 (Ar*C*), 130.8 (q, *J*_C-F_ 2.7, Ar*C*H), 128.2 (Ar*C*H), 127.2 (Ar*C*), 127.0 (q, *J*_C-F_ 5.9, Ar*C*H), 126.7 (d, *J*_C-F_ 3.0, Ar*C*), 126.5 (Ar*C*), 124.3 (q, *J*_C-F_ 274.0, *C*F_3_), 124.0 (Ar*C*H), 121.0 (Ar*C*), 8.7 (*C*H_3_); *δ*_F_(377 MHz; CDCl_3_) −59.0; *m/z* (EI^+^) 330.0496 (90%, M^+^ C_18_H_9_O_3_F_3_ requires 330.0498), 261 [100, (M-CF_3_)^+^].

#### 6-Fluoro-1-methylphenanthro[1,2-*b*]furan-10,11-dione 37 and 4-fluoro-8-methylphenanthro[3,2-*b*]furan-7,11-dione 38

General procedure **A** was followed, using the unpurified alcohol **30** (140 mg) to give the dione **37** (18 mg, 2% over 3 steps) as a dark red solid; mp 187-191 °C; ν_max_(NaCl discs)/cm^-1^ 1675 (C=O), 1664 (C=O), 1594 (C=C); *δ*_H_(400 MHz; CDCl_3_) 9.20 (1 H, d, *J* 8.8, ArC*H*), 8.45 (1 H, d, *J* 8.8, ArC*H*), 7.89 (1 H, d, *J* 8.8, ArC*H*), 7.67-7.60 (1 H, m, ArC*H*), 7.36 (1 H, q, *J* 1.2, OC*H*), 7.22 (1 H, ddd, *J* 10.0, 7.8, 0.7, ArC*H*), 2.32 (3 H, d, *J* 1.2, C*H*_3_); *δ*_C_(100 MHz; CDCl_3_) 183.0 (*C*=O), 175.0 (*C*=O), 160.5 (Ar*C*), 158.8 (d, *J*_C-F_ 253.6, Ar*C*), 142.5 (Ar*C*H), 133.2 (d, *J*_C-F_ 2.5, Ar*C*), 131.0 (d, *J*_C-F_ 8.5, Ar*C*H), 130.9 (Ar*C*), 129.4 (d, *J*_C-F_ 7.3, Ar*C*H), 124.7 (d, *J*_C-F_ 15.4, Ar*C*), 122.5 (d, *J*_C-F_ 4.7, Ar*C*H), 122.3 (d, *J*_C-F_ 2.3, Ar*C*), 122.0 (Ar*C*), 121.0 (Ar*C*), 119.3 (d, *J*_C-F_ 1.6, Ar*C*H), 111.1 (d, *J*_C-F_ 19.3, Ar*C*H), 8.8 (*C*H_3_); *δ*_F_(377 MHz; CDCl_3_) −120.7; *m/z* (ESI^+^) 326 (34%, M+2Na^+^), 303 (20, M+Na^+^), 281.0612 (100, M+H^+^ C_17_H_10_O_3_F requires 281.0608). Also obtained was the dione **38** (12 mg, 1% over 3 steps) as a pale orange solid; mp 185-189 °C; ν_max_(NaCl discs)/cm^-1^ 1662 (C=O), 1591 (C=O), 1535 (C=C); *δ*_H_(400 MHz; CDCl_3_) 9.56 (1 H, d, *J* 8.9, ArC*H*), 8.50 (1 H, d, *J* 8.9, ArC*H*), 8.34 (1 H, d, *J* 8.9, ArC*H*), 7.71-7.64 (1 H, m, ArC*H*), 7.55 (1 H, d, *J* 1.0, OC*H*), 7.32 (1 H, dd, *J* 9.9, 7.8, ArC*H*), 2.41 (3 H, d, *J* 1.0, C*H*_3_); *δ*_C_(100 MHz; CDCl_3_) 181.8 (*C*=O), 176.7 (*C*=O), 158.4 (d, *J*_C-F_ 253.0, Ar*C*), 153.6 (Ar*C*), 145.5 (Ar*C*H), 134.5 (Ar*C*), 132.0 (d, *J*_C-F_ 2.8, Ar*C*), 130.0 (d, *J_C-F_* 8.1, Ar*C*H), 127.5 (d, *J*_C-F_ 6.9, Ar*C*H), 127.1 (d, *J*_C-F_ 15.8, Ar*C*), 126.8 (d, *J*_C-F_ 2.6, Ar*C*), 126.7 (Ar*C*), 124.0 (d, *J*_C-F_ 4.6, Ar*C*H), 122.6 (d, *J*_C-F_ 1.6, Ar*C*H), 121.0 (Ar*C*), 112.1 (d, *J*_C-F_ 19.3, Ar*C*H), 8.7 (*C*H_3_); *δ*_F_(377 MHz; CDCl_3_) −120.9; *m/z* (EI^+^) 280.0532 (100%, M^+^ C_17_H_9_O_3_F requires 280.0530).

#### 6-Methoxy-1-methylphenanthro[1,2-*b*]furan-10,11-dione 39 and 4-methoxy-8-methylphenanthro[3,2-*b*]furan-7,11-dione 40

General procedure **A** was followed, using the alcohol **31** (100 mg, 0.39 mmol) to give the dione **39** (22 mg, 5% over 3 steps) as a dark brown solid; mp 249-253 °C; ν_max_(NaCl discs)/cm^-1^ 1670 (C=O), 1660 (C=O), 1587 (C=C), 1548 (C=C); *δ*_H_(400 MHz; CDCl_3_) 9.00 (1 H, d, *J* 8.8, ArC*H*), 8.66 (1 H, d, *J* 8.8, ArC*H*), 7.82 (1 H, d, *J* 8.8, ArC*H*), 7.62 (1 H, br t, *J* 8.4, ArC*H*), 7.34 (1 H, s, ArC*H*), 6.89 (1 H, d, *J* 7.8, ArC*H*), 4.04 (3 H, s, OC*H*_3_), 2.32 (3 H, s, C*H*_3_); *δ*_C_(126 MHz; CDCl_3_) 183.4 (*C*=O), 175.7 (*C*=O), 161.2 (Ar*C*), 155.7 (Ar*C*), 142.1 (Ar*C*H), 133.5 (Ar*C*), 131.5 (Ar*C*H), 131.2 (Ar*C*H), 130.5 (Ar*C*), 126.8 (Ar*C*), 122.4 (Ar*C*), 121.8 (Ar*C*), 120.7 (Ar*C*), 118.5 (Ar*C*H), 118.2 (Ar*C*H), 105.5 (Ar*C*H), 55.7 (O*C*H_3_), 8.8 (*C*H_3_); *m/z* (ESI^+^) 315 (11%, M+Na^+^), 293.0813 (100, M+H^+^ C_18_H_13_O_4_ requires 293.0808). Also obtained was the dione **40** (21 mg, 5% over 3 steps) as a red solid; mp 231-234 °C; ν_max_(ATR)/cm^-1^ 1662 (C=O), 1582 (C=O), 1535 (C=C); *δ*_H_(400 MHz; CDCl_3_) 9.32 (1 H, d, *J* 8.8, ArC*H*), 8.68 (1 H, d, *J* 8.8, ArC*H*), 8.24 (1 H, d, *J* 8.8, ArC*H*), 7.63 (1 H, dd, *J* 8.8, 7.9, ArC*H*), 7.51 (1 H, d, *J* 0.8, ArC*H*), 6.95 (1 H, d, *J* 7.9, ArC*H*), 4.04 (3 H, s, OC*H*_3_), 2.40 (3 H, d, *J* 0.8, C*H*_3_); *δ*_C_(101 MHz; CDCl_3_) 182.3 (*C*=O), 177.2 (*C*=O), 155.3 (Ar*C*), 153.9 (Ar*C*), 145.1 (Ar*C*H), 134.4 (Ar*C*), 132.0 (Ar*C*), 130.5 (Ar*C*H), 129.3 (Ar*C*), 129.2 (Ar*C*H), 126.51 (Ar*C*), 126.49 (Ar*C*), 121.6 (Ar*C*H), 120.8 (Ar*C*), 119.8 (Ar*C*H), 106.2 (Ar*C*H), 55.7 (O*C*H_3_), 8.8 (*C*H_3_); *m/z* (ESI^+^) 315 (43%, M+Na^+^), 293.0810 (100, M+H^+^ C_18_H_13_O_4_ requires 293.0808).

#### 1,6-Dimethylphenanthro[1,2-*b*]furan-10,11-dimethyldioxole, TI-acetal 43

TI **1** (27 mg, 0.10 mmol) was dissolved in anhydrous THF (7 mL) and stirred under a nitrogen atmosphere at room temperature. Sodium borohydride (4.0 mg, 0.10 mmol) was added in a single portion, and the solution stirred for 30 minutes. The solution was poured onto ice/water (20 mL), acidified with aqueous HCl solution (1 M, 3 mL), and extracted with DCM (3 x 20 mL). The organic extracts were combined, dried (MgSO_4_), filtered and concentrated *in vacuo* to give the crude diol **42** as an olive-green solid (29 mg), which was immediately dissolved in anhydrous toluene (10 mL). 2,2-Dimethoxypropane (0.025 mL, 0.20 mmol) and *poro*-toluenesulfonic acid monohydrate (25 mg, 0.13 mmol) were added, and the reaction was stirred under a nitrogen atmosphere at reflux for 2 h. The mixture was cooled to room temperature, aqueous saturated NaHCO3 solution (20 mL) added, and extracted with diethyl ether (3 x 20 mL). The organic extracts were combined, dried (MgSO_4_), filtered and concentrated *in vocuo* to give an off-white solid (38 mg), which was purified twice by flash column chromatography (silica gel, DCM, and then silica gel, 19:1 40/60 petroleum ether/ethyl acetate) to give the acetal **43** as a beige solid (11 mg, 34%); mp 176-180 °C; νmax(ATR)/cm^-1^ 2917 (C-H); *δ*_H_(400 MHz; CDCl_3_) 9.13 (1 H, d, *J* 8.4, ArC*H*), 8.22 (1 H, d, *J* 9.2, ArC*H*), 7.89 (1 H, dd, *J* 9.2, 0.8, ArC*H*), 7.57-7.52 (2 H, m, 2 x ArC*H*), 7.45 (1 H, dt, *J* 7.1, 0.9, ArC*H*), 2.79 (3 H, s, ArC*H*_3_), 2.46 (3 H, d, *J* 1.3, ArC*H*_3_), 1.91 (6 H, s, 2 x C*H*_3_); *δ*_C_(126 MHz; CDCl_3_) 148.3 (Ar*C*), 141.4 (Ar*C*H), 138.2 (Ar*C*), 137.8 (Ar*C*), 134.1 (Ar*C*), 130.7 (Ar*C*), 128.9 (Ar*C*), 127.3 (Ar*C*H), 125.6 (Ar*C*H), 125.4 (Ar*C*H), 120.4 (Ar*C*H), 119.2 (Ar*C*H), 118.5 (Ar*C*), 114.5 (Ar*C*), 114.3 (Ar*C*), 113.0 (Ar*C*), 112.7 (Ar*C*), 26.1 (2 x *C*H_3_), 20.2 (Ar*C*H_3_), 9.2 (Ar*C*H_3_); *m/z* (ESI^+^) 319.1322 (100%, M+H^+^ C21H19O3 requires 319.1329), 383 (8), 359 (28), 261 (22).

#### β-Lapachone 5

Concentrated H_2_SO_4_ (1.0 mL) was added slowly to lapachol **44** (25 mg, 0.10 mmol), with stirring, until the solid completely dissolved. The solution was stirred for a further 5 minutes and poured onto ice (20 g). The precipitate was filtered off by vacuum filtration and washed with cold water (5 mL) to give an orange solid. The crude product was purified by flash column chromatography (silica gel, DCM – 19:1 DCM/MeOH) to give the dione **5** (21 mg, 85%) as an orange solid; mp 151-153 °C (lit.^84^ 152-154 °C); *δ*_H_(400 MHz; CDCl_3_) 8.07 (1 H, dd, *J* 7.7, 1.0, ArC*H*), 7.83 (1 H, d, *J* 7.7, ArC*H*), 7.66 (1 H, td, *J* 7.7, 1.0, ArC*H*), 7.52 (1 H, td, *J* 7.7, 1.0, ArC*H*), 2.58 (2 H, t, *J* 6.7, C*H*2), 1.87 (2 H, t, *J* 6.7, C*H*2), 1.48 (6 H, s, 2 x C*H*_3_); *δ*_C_(101 MHz; CDCl_3_) 179.9 (*C*=O), 178.6 (*C*=O), 162.1 (Ar*C*), 134.8 (Ar*C*H), 132.6 (Ar*C*), 130.7 (Ar*C*H), 130.1 (Ar*C*), 128.6 (Ar*C*H), 124.1 (Ar*C*H), 112.7 (Ar*C*), 79.3 [*C*(CH_3_)_2_], 31.6 (*C*H_2_), 26.8 (2 x *C*H_3_), 16.2 (*C*H_2_). All data were in general agreement with the literature.^84,85^

#### Norlapachol 45

Lapachol **44** (150 mg, 0.62 mmol) was dissolved in THF (5 mL), sodium carbonate (72 mg, 0.68 mmol) in water (5 mL) was added, and the solution was heated to 60 °C. Aqueous hydrogen peroxide (30%, 1.0 mL, 13 mmol) was added and the solution was heated at 60 °C for 4 h. The reaction was cooled to rt, acidified with concentrated HCl (0.2 mL), and quenched with sodium sulfite (3.0 g). Aqueous NaOH (25%, 5 mL) and copper(II) sulfate (625 mg, 3.9 mmol) in water (5 mL) were added and the mixture was stirred at room temperature for 1.5 h. The solution was filtered through a pad of Celite, acidified with concentrated HCl (7 mL) and extracted with diethyl ether (3 x 75 mL). The organic extracts were combined, washed with brine (200 mL), dried (MgSO_4_), filtered and concentrated *in vacuo* to give an orange solid. The crude product was purified by recrystallisation from *n*-hexane to give the dione **45** (73 mg, 52%) as an orange solid; mp 123-125 °C (from *n*-hexane), (lit.^85^ 121-122 °C); ν_max_(NaCl discs)/cm^-1^ 3364 (O-H), 2928 (C-H), 2910 (C-H), 1662 (C=O), 1645 (C=O), 1627 (C=C), 1593 (C=C); *δ*_H_(400 MHz; CDCl_3_) 8.15 (1 H, d, *J* 7.6, ArC*H*), 8.11 (1 H, d, *J* 7.6, ArC*H*), 7.78 (1 H, td, *J* 7.6, 1.1, ArC*H*), 7.71 (1 H, td, *J* 7.6, 1.1, ArC*H*), 7.55 (1 H, br s, O*H*), 6.02 (1 H, br s, C*H*), 2.01 (3 H, s, C*H*_3_), 1.70 (3 H, s, C*H*_3_); *δ*_C_(101 MHz; CDCl_3_) 184.8 (*C*=O), 181.6 (*C*=O), 151.2 (*C*), 143.6 (*C*), 134.9 (Ar*C*H), 133.0 (Ar*C*H), 132.9 (*C*), 129.5 (*C*), 126.9 (Ar*C*H), 126.1 (Ar*C*H), 120.9 (*C*), 113.7 (*C*H), 26.6 (*C*H_3_), 21.8 (*C*H_3_); *m/z* (ESI^+^) 251 (5%, M+Na^+^), 229.0862 (100, M+H^+^ C_14_H_13_O_3_ requires 229.0859). All data were in general agreement with the literature.^85–87^

#### Nor-β-lapachone 6

Concentrated H_2_SO_4_ (1.5 mL) was added slowly to norlapachol **45** (30 mg, 0.13 mmol), with stirring, until the solid completely dissolved. The solution was stirred at room temperature for a further 5 minutes, poured onto ice (20 g) and rinsed with water (20 mL). The solution was extracted with DCM (3 x 40 mL), and the organic extracts were combined, washed with brine (100 mL), dried (MgSO_4_), filtered, and concentrated *in vocuo* to give an orange/pink solid. The crude product was purified by trituration with cold hexane (2 x 2 mL) and dried under high vacuum to give the dione **6** (29 mg, 95%) as an orange solid; mp 190-193 °C (lit.^88^ 188-189 °C); *δ*_H_(400 MHz; CDCl_3_) 8.10 (1 H, d, *J* 7.6, ArC*H*), 7.68-7.64 (2 H, m, 2 x ArC*H*), 7.63-7.57 (1 H, m, ArC*H*), 2.97 (2 H, s, C*H*_2_), 1.63 (6 H, s, 2 x C*H*_3_); *δ*_C_(101 MHz; CDCl_3_) 181.4 (*C*=O), 175.7 (*C*=O), 168.8 (Ar*C*), 134.5 (Ar*C*H), 131.9 (Ar*C*H), 130.9 (Ar*C*), 129.3 (Ar*C*H), 127.9 (Ar*C*), 124.6 (Ar*C*H), 115.0 (Ar*C*), 93.8 [*C*(CH_3_)_2_], 39.3 (*C*H_2_), 28.4 (2 x *C*H_3_). All data were in general agreement with the literature.^85,88^

### Biological evaluation

#### Reagents

All reagents were obtained from Sigma Aldrich/Merck (Darmstadt, Germany) unless stated otherwise. The c-Jun N-terminal kinase inhibitor SP600125 was obtained from StressMarq Biosciences (Victoria, Canada), and tanshinone IIA (TIIA) **2** was obtained from Generon (Slough, UK). All synthetic compounds were synthesised using chemistry as outlined in the previous section. All compounds were dissolved in dimethyl sulfoxide (DMSO).

#### Zebrafish husbandry

Zebrafish (*Donio rerio*) were raised and maintained in UK Home Office-approved aquaria at the Bateson Centre at The University Of Sheffield, in accordance with standard protocols and procedures.^89^ Transgenic zebrafish larvae which express GFP specifically in neutrophils, *Tg(mpx:GFP)i114*,^90^ 3 days post-fertilisation (dpf), were used for all experiments.

#### Zebrafish experiments

All compounds were diluted from the stock solutions and administered by immersion in aqueous solutions. DMSO (0.5% concentration) was used as the negative control. In neutrophil recruitment experiments, SP600125 (30 μM) or tanshinone I (TI) **1** (25 μM) was the positive control, whilst in inflammation resolution experiments, TIIA **2** (25 μM) was used.

Zebrafish larvae were injured by transection of the tailfin using a micro-scalpel blade, as posterior as possible in the region where the pigment pattern was disrupted, inducing an inflammatory response. Neutrophil counting was performed according to standard procedures used previously.^64,65,90^ To assess neutrophil recruitment, zebrafish larvae were injured and treated immediately with test compounds at the stated concentrations. At 4 or 6 hours post-injury (hpi), larvae were anaesthetised and the number of neutrophils at the site of injury was counted. To study the resolution of neutrophilic inflammation, zebrafish larvae were injured and allowed to recover. Larvae that mounted a good inflammatory response (around 20-25 neutrophils at the site of injury; termed ‘good responders’) were selected for compound treatment at the stated concentrations at 4 hpi. At 8 hpi, larvae were anaesthetised and the number of neutrophils at the site of injury was counted. All experiments were performed blind to treatment groups.

#### Statistical analysis

All statistical analysis was carried out using Prism 7.0 (GraphPad, San Diego, USA), version 7.00. For all experiments, at least three biological repeats were performed, from which data were pooled and then analysed. In all figures, data are shown as mean ± SEM of all data points from individual larvae. Statistical analysis was performed using a one-way analysis of variance (ANOVA) test with Dunnett’s multiple comparison post-test for comparing treatment neutrophil numbers to those of the DMSO negative control.

## Acknowledgements

The authors thank Dr. Anne L. Robertson for initial assistance with zebrafish techniques, Catherine A. Loynes for advice, and the Bateson Centre aquarium staff for providing expert zebrafish care. This work was supported by a Medical Research Council (MRC) Programme Grant to S.A.R. (MR/M004864/1) and an MRC Centre Grant G0700091. The authors also thank The University Of Sheffield for funding (M.J.F.).

## Supplementary Information (SI) available

Chemical synthesis of compounds **8-9**, **11**, **13-31**; information on DoE studies; and copies of ^1^H, ^13^C and ^19^F NMR spectra.

