## Supplementary Information for "Evaluation of the anti-inflammatory effects of synthesised tanshinone I and isotanshinone I analogues in zebrafish"

#### Supplementary Information (SI)

Experimental - chemical synthesis of compounds **8-9**, **11**, **13-31**

Information on Design of Experiments (DoE) studies

NMR spectra of synthesised compounds

References

#### Experimental

##### General reagents, materials and methods

All chemicals used were purchased from commercial suppliers and were used as received without further purification. Melting points were determined using a Gallenkamp melting point apparatus equipped with a thermometer. IR spectroscopy was performed on a PerkinElmer FT-IR Spectrum 65 or Spectrum 100 spectrometer, using NaCl discs. $^{1}$H and $^{13}$C and NMR experiments were run on either a Bruker Avance 400 or Bruker Avance III HD 500 spectrometer at 298 K. Chemical shifts (δ) are reported in parts per million (ppm) relative to the deuterated lock solvent as an internal standard, where s = singlet, d = doublet, t = triplet, q = quartet, m = multiplet, br s = broad singlet, br d = broad doublet, br t = broad triplet, app d = apparent doublet, app t = apparent triplet, dd = doublet of doublets, ddd = double doublet of doublets, td = triplet of doublets, dq = doublet of quartets. All coupling constants are reported in hertz, Hz. TLC was performed on Merck silica gel 60 $F_{254}$ aluminium-backed plates and visualised using ultraviolet light followed by staining with potassium permanganate dip. Column chromatography was carried out using silica gel obtained from VWR Chemicals, particle size 40-63 µm.

##### 2-Bromo-6-methoxy-1,4-hydroquinone 8

Sodium percarbonate (18.71 g, 119.2 mmol) was added to a solution of 5-bromovanillin **7** (25.03 g, 108.4 mmol) in THF (300 mL) and water (120 mL) and stirred at room temperature for 5 h. The reaction was quenched with sodium sulfite (15.00 g), filtered, and concentrated *in vacuo*. The organic product was extracted with ethyl acetate (3 x 300 mL), dried ($MgSO_4$), filtered and concentrated *in vacuo* to give the hydroquinone **8** (20.34 g, 86%) as a grey solid which was used without further purification; mp 142-146 °C (lit.[1] 139-141 °C); $\delta_H$[400 MHz; $(CD_3)_2CO$] 8.13 (1 H, s, O*H*), 7.57 (1H, s, O*H*), 6.59 (1 H, d, *J* 2.6, ArC*H*), 6.50 (1 H, d, *J* 2.6, ArC*H*), 3.82 (3 H, s, OC*H*$_3$); $\delta_C$[100 MHz; $(CD_3)_2CO$] 150.7 (Ar*C*), 148.6 (Ar*C*), 137.3 (Ar*C*), 110.0 (Ar*C*H), 108.1 (Ar*C*), 99.6 (Ar*C*H), 55.6 (O*C*H$_3$). All data were in agreement with the literature.[1]

##### 2-Bromo-6-methoxy-[1,4]-benzoquinone 9

A solution of iron(III) chloride hexahydrate (125.4 g, 463.9 mmol) in water (600 mL) was added to a solution of 2-bromo-6-methoxy-1,4-hydroquinone **8** (20.32 g, 92.77 mmol) in methanol

(80 mL) with stirring. The resulting mixture was stirred at room temperature for 5 h. The organic layer was separated and extracted with DCM (3 x 250 mL), combined, washed with water (500 mL) and brine (500 mL), dried ($MgSO_4$), filtered and concentrated *in vacuo* to give the benzoquinone **9** (19.84 g, 99%) as an orange solid which was used without further purification; mp 152-156 °C (lit.[1] 160-162 °C); $\delta_H$(400 MHz; $CDCl_3$) 7.23 (1 H, d, *J* 2.2, C*H*CO), 5.98 (1 H, d, *J* 2.2, C*H*CO), 3.88 (3 H, s, OC*H*$_3$); $\delta_C$(100 MHz; $CDCl_3$) 184.6 (*C*), 174.5 (*C*), 158.3 (*C*), 138.5 (=*C*H), 134.3 (*C*), 107.7 (=*C*H), 56.9 (O*C*H$_3$). All NMR data were in agreement with the literature.[1]

**General representative procedure A for radical decarboxylative alkylation reactions to form bromides 11, 13-16**

A mixture of the quinone **9** (2.00 g, 9.26 mmol), carboxylic acid (11.1 mmol) and silver(I) phosphate (1.94 g, 4.63 mmol) in acetonitrile (70 mL) was stirred under a nitrogen atmosphere in the dark and heated at reflux. A solution of ammonium persulfate (4.23 g, 18.5 mmol) in water (70 mL) was added slowly over 20 minutes *via* dropping funnel, and the resulting mixture was stirred at reflux for 2.5 h. The solution was cooled to room temperature and poured onto ice (50 g). Aqueous NaOH (1 M, 10 mL) and water (50 mL) were added and the organic layer was separated and extracted with DCM (3 × 150 mL). The organic extracts were combined, washed with brine (400 mL), dried ($MgSO_4$), filtered and concentrated *in vacuo* to give material which was purified by flash column chromatography (silica gel, DCM) to give the bromides **11, 13-16**.

**3-Bromo-5-methoxy-2-[2-(2-methylphenyl)ethyl]-2,5-cyclohexadiene-1,4-dione 11**

General procedure **A** was followed, using 3-(2-methylphenyl)propionic acid **10** (1.82 g), to give the bromide **11** (1.67 g, 54 %) as a yellow solid; mp 179-184 °C (lit.[2] mp 149-152 °C); $\delta_H$(400 MHz; $CDCl_3$) 7.24-7.14 (4 H, m, 4 × ArC*H*), 5.99 (1 H, s, C*H*CO), 3.88 (3 H, s, OC*H*$_3$), 2.98-2.92 (2 H, m, C*H*$_2$), 2.82-2.76 (2 H, m, C*H*$_2$), 2.45 (3 H, s, ArC*H*$_3$); $\delta_C$(100 MHz; $CDCl_3$) 183.7 (*C*=O), 174.8 (*C*=O), 158.3 (*C*), 149.0 (*C*), 138.6 (*C*), 136.2 (*C*), 133.2 (*C*), 130.4 (ArCH), 129.1 (Ar*C*H), 126.6 (ArCH), 126.2 (Ar*C*H), 107.4 (*C*HCO), 56.7 (O*C*H$_3$), 32.0 (*C*H$_2$), 31.3 (*C*H$_2$), 19.3 (*C*H$_3$). All NMR data were in agreement with the literature.[2]

**3-Bromo-5-methoxy-2-(2-phenylethyl)-2,5-cyclohexadiene-1,4-dione 13**

General procedure **A** was followed, but on a larger scale, using the quinone **9** (3.00 g, 13.9 mmol), 3-phenylpropionic acid **12** (2.50 g, 16.7 mmol) and silver(I) phosphate (4.82 g, 11.5 mmol) in acetonitrile (150 mL), and ammonium persulfate (6.34 g, 27.8 mmol) in water (100 mL), to give the bromide **13** (2.04 g, 46%) as a yellow solid; mp 138-141 °C (lit.[2] 138-140 °C); $\delta_H$(400 MHz; $CDCl_3$) 7.36-7.22 (5 H, m, 5 x ArC*H*), 5.98 (1 H, s, C*H*CO), 3.87 (3 H, s, OC$H_3$), 3.03-2.97 (2 H, m, C$H_2$), 2.83-2.77 (2 H, m, C$H_2$); $\delta_C$(100 MHz; $CDCl_3$) 183.6 (*C*=O), 174.8 (*C*=O), 158.3 (*C*), 149.0 (*C*), 140.4 (*C*), 133.3 (*C*), 128.55 (2 x Ar*C*H), 128.49 (2 x Ar*C*H), 126.5 (Ar*C*H), 107.4 (=*C*H), 56.7 (O*C*$H_3$), 33.8 (*C*$H_2$), 33.2 (*C*$H_2$). All data were in agreement with the literature.[2]

**3-Bromo-5-methoxy-2-[2-(trifluoromethyl)phenethyl]-2,5-cyclohexadiene-1,4-dione 14**

General procedure **A** was followed, using 3-[2-(trifluoromethyl)phenyl]propionic acid (2.42 g), to give the bromide **14** (1.35 g, 38%) as a yellow solid; mp 144-147 °C (lit.[2] 143-145 °C); $\delta_H$(400 MHz; $CDCl_3$) 7.66 (1 H, d, *J* 7.6, ArC*H*), 7.52 (1 H, t, *J* 7.6, ArC*H*), 7.46 (1 H, d, *J* 7.6, ArC*H*), 7.35 (1 H, t, *J* 7.6, ArC*H*), 6.00 (1 H, s, C*H*CO), 3.88 (3 H, s, OC$H_3$), 3.08-2.95 (4 H, m, 2 x C$H_2$); $\delta_C$(100 MHz; $CDCl_3$) 183.5 (*C*=O), 174.7 (*C*=O), 158.3 (*C*), 148.4 (*C*), 139.0 (*C*), 133.6 (*C*), 131.9 (Ar*C*H), 131.4 (Ar*C*H), 128.6 (q, $J_{C\text{-}F}$ 30.0, Ar*C*), 126.6 (Ar*C*H), 126.0 (q, $J_{C\text{-}F}$ 5.7, Ar*C*H), 124.5 (q, $J_{C\text{-}F}$ 273.6, *C*$F_3$), 107.4 (=*C*H), 56.7 (O*C*$H_3$), 32.8 (*C*$H_2$), 30.3 (*C*$H_2$). All data were in agreement with the literature.[2]

**3-Bromo-2-(2-fluorophenethyl)-5-methoxy-2,5-cyclohexadiene-1,4-dione 15**

General procedure **A** was followed, using 3-(2-fluorophenyl)propionic acid (1.87 g), to give the bromide **15** (1.44 g, 46%) as a yellow solid; mp 144-148 °C (lit.[2] 134-137 °C); $\delta_H$(400 MHz; $CDCl_3$) 7.25-7.19 (2 H, m, 2 x ArC*H*), 7.11-6.99 (2 H, m, 2 x ArC*H*), 5.97 (1 H, s, C*H*CO), 3.87 (3 H, s, OC$H_3$), 3.06-2.99 (2 H, m, C$H_2$), 2.92-2.85 (2 H, m, C$H_2$); $\delta_C$(100 MHz; $CDCl_3$) 183.6 (*C*=O), 174.8 (*C*=O), 161.2 (d, $J_{C\text{-}F}$ 245.4, Ar*C*F), 158.2 (*C*), 148.6 (*C*), 133.5 (*C*), 130.8 (d, $J_{C\text{-}F}$ 4.6, Ar*C*H), 128.3 (d, $J_{C\text{-}F}$ 8.1, Ar*C*H), 127.1 (d, $J_{C\text{-}F}$ 16.2, Ar*C*), 124.1 (d, $J_{C\text{-}F}$ 3.4, Ar*C*H), 115.3 (d, $J_{C\text{-}F}$ 22.0, Ar*C*H), 107.3 (=*C*H), 56.7 (O*C*$H_3$), 31.5 (*C*$H_2$), 27.2 (*C*$H_2$). All NMR data were in agreement with the literature.[2]

**3-Bromo-5-methoxy-2-(2-methoxyphenethyl)-2,5-cyclohexadiene-1,4-dione 16**

General procedure **A** was followed, using 3-(2-methoxyphenyl)propionic acid (2.00 g), to give an inseparable mixture of the bromide **16** and the benzoquinone **9** (1.07 g) as a yellow solid in a 60:40 ratio which was used without further purification. Analysis of the $^1$H NMR spectrum of this mixture indicated a 33% yield of the bromide **16**. A sample of this mixture was further purified for analytical purposes to give the bromide **16** as a yellow solid; mp 160-163 °C (lit.[2] 161-163 °C); $\delta_H$(400 MHz; $CDCl_3$) 7.22 (1 H, td, *J* 7.7, 1.6, ArC*H*), 7.15 (1 H, dd, *J* 7.7, 1.6, ArC*H*), 6.89 (1 H, td, *J* 7.7, 0.7, ArC*H*), 6.84 (1 H, d, *J* 7.7, ArC*H*), 5.96 (1 H, s, C*H*CO), 3.86 (3 H, s, $OCH_3$), 3.84 (3 H, s, $OCH_3$), 3.04-2.98 (2 H, m, $CH_2$), 2.88-2.82 (2 H, m, $CH_2$); $\delta_C$(100 MHz; $CDCl_3$) 183.7 (*C*=O), 174.9 (*C*=O), 158.1 (*C*), 157.6 (*C*), 149.6 (*C*), 132.9 (*C*), 130.1 (ArCH), 128.6 (*C*), 127.8 (ArCH), 120.5 (Ar*C*H), 110.2 (Ar*C*H), 107.3 (ArCH), 56.7 ($OCH_3$), 55.3 ($OCH_3$), 31.3 ($CH_2$), 28.7 ($CH_2$). All data were in general agreement with the literature.[2]

**General representative procedure B for Heck reactions to form diones 17-26**

A solution of the bromide **11, 13-16** (2.24 mmol), palladium(II) acetate (25 mg, 0.111 mmol), triphenylphosphine (59 mg, 0.225 mmol) and potassium carbonate (930 mg, 6.73 mmol) in degassed toluene (80 mL $mmol^{-1}$) was heated at reflux in the dark for 17 h under a nitrogen atmosphere, with stirring. The mixture was cooled to room temperature, concentrated *in vacuo*, diluted with water (40 mL $mmol^{-1}$), and extracted with DCM (3 x 40 mL $mmol^{-1}$). The organic extracts were combined, washed with brine (80 mL $mmol^{-1}$), dried ($MgSO_4$), filtered and concentrated *in vacuo* to give material which was separated from major impurities by flash column chromatography (silica gel, DCM) to give an inseparable mixture of the fully aromatised and non-aromatised diones **17-26**.

**3-Methoxy-8-methylphenanthrene-1,4-dione 17 and 9,10-dihydro-3-methoxy-8-methylphenanthrene-1,4-dione 18**

General procedure **B** was followed, using the bromide **11** (750 mg, 2.24 mmol), to give a bright orange solid (378 mg, 67%) as an inseparable mixture of diones **17** and **18** in a 75:25 ratio which was used without further purification. Selected $^1$H NMR data from product mixture corresponding to the dione **17**: $\delta_H$(400 MHz; $CDCl_3$) 9.42 (1 H, d, *J* 8.8, ArC*H*), 8.43 (1 H, br d, *J* 8.8, ArC*H*), 8.25 (1 H, br d, *J* 8.8, ArC*H*), 7.65 (1 H, dd, *J* 8.8, 6.9, ArC*H*), 7.50 (1 H, br d, *J* 6.9, ArC*H*), 6.17 (1 H, br s, C*H*CO), 3.97 (3 H, br s, $OCH_3$), 2.77 (3 H, s, $CH_3$). All data were in general agreement with the literature.[3] Selected $^1$H NMR data from product mixture corresponding to the dione **18**: $\delta_H$(400 MHz; $CDCl_3$) 7.92-7.86 (1 H, app t, *J* 4.5, ArC*H*), 7.23 (2 H, d, *J* 4.5, ArC*H*), 5.99 (1 H, br s, C*H*CO), 3.88 (3 H, br s, $OCH_3$), 2.79-2.74 (2 H, m, $CH_2$), 2.74-2.70 (2 H, m, $CH_2$), 2.36 (3 H, s, $CH_3$). No data were reported in the literature.

**3-Methoxyphenanthrene-1,4-dione 19 and 9,10-dihydro-3-methoxyphenanthrene-1,4-dione 20**

General procedure **B** was followed, using the bromide **13** (200 mg, 0.622 mmol), to give a pale red solid (109 mg, 73%) as an inseparable mixture of diones **19** and **20** in a 96:4 ratio which was used without further purification. Selected $^1$H NMR data from product mixture corresponding to the dione **19**: $\delta_H$(400 MHz; $CDCl_3$) 9.55 (1 H, d, *J* 8.8, ArC*H*), 8.20 (2 H, app d, *J* 1.2, 2 x ArC*H*), 7.92 (1 H, d, *J* 8.1, ArC*H*), 7.77 (1 H, ddd, *J* 8.8, 6.8, 1.2, ArC*H*), 7.66 (1 H, ddd, *J* 8.1, 6.8, 1.2, ArC*H*), 6.17 (1 H, s, C*H*CO), 3.96 (3 H, s, $OCH_3$). All data were in general agreement with the literature.[4,5] Selected $^1$H NMR data from product mixture corresponding to the dione **20**: $\delta_H$(400 MHz; $CDCl_3$) 5.99 (1 H, s, C*H*CO), 3.88 (3 H, s, $OCH_3$), 2.83-2.78 (2 H, m, $CH_2$), 2.76-2.72 (2 H, m, $CH_2$). No data were reported in the literature.

**3-Methoxy-8-(trifluoromethyl)phenanthrene-1,4-dione 21 and 9,10-dihydro-3-methoxy-8-trifluoromethyl)phenanthrene-1,4-dione 22**

General procedure **B** was followed, using the bromide **14** (1.35 g, 3.47 mmol), to give a dark orange solid (500 mg) as an inseparable mixture including diones **21** and **22** in a 65:35 ratio alongside an additional unidentified compound, which was used without further purification. Selected $^1$H NMR peaks from product mixture corresponding to the dione **21**: $\delta_H$(400 MHz; $CDCl_3$) 9.81 (1 H, d, *J* 8.9, ArC*H*), 8.59 (1 H, dq, *J* 8.9, 0.9, ArC*H*), 8.37 (1 H, d, *J* 8.9, ArC*H*), 8.06 (1 H, d, *J* 7.2, ArC*H*), 7.86-7.81 (1 H, m, ArC*H*), 6.23 (1 H, s, C*H*CO), 3.99 (3 H, s, $OCH_3$). Selected $^1$H NMR peaks from product mixture corresponding to the dione **22**: $\delta_H$(400 MHz; $CDCl_3$) 5.97 (1 H, s, C*H*CO), 3.85 (3 H, s, $OCH_3$), 3.04-2.97 (2 H, m, $CH_2$), 2.80-2.74 (2 H, m, $CH_2$). Selected $^1$H NMR peaks from product mixture corresponding to the unidentified compound: $\delta_H$(400 MHz; $CDCl_3$) 6.01(1 H, s, =C*H*), 3.89 (3 H, s, $OCH_3$). No data were reported in the literature.

**8-Fluoro-3-methoxyphenanthrene-1,4-dione 23 and 9,10-dihydro-8-fluoro-3-methoxyphenanthrene-1,4-dione 24**

General procedure **B** was followed, using the bromide **15** (1.34 g, 3.95 mmol), to give a dark orange solid (197 mg) as an inseparable mixture including diones **23** and **24** in an 80:20 ratio alongside an additional unidentified compound, which was used without further purification. Selected $^1$H NMR peaks from product mixture corresponding to the dione **23**: $\delta_H$(400 MHz; $CDCl_3$) 9.35 (1 H, d, *J* 8.8, ArC*H*), 8.54 (1 H, d, *J* 8.8, ArC*H*), 8.29 (1 H, d, *J* 8.8, ArC*H*), 7.74-7.67 (1 H, m, ArC*H*), 7.37-7.32 (1 H, m, ArC*H*), 6.21 (1 H, s, C*H*CO), 3.98 (3 H, s, $OCH_3$). Selected $^1$H NMR peaks from product mixture corresponding to the dione **24**: $\delta_H$(400 MHz; $CDCl_3$) 5.95 (1 H, s, =C*H*), 3.84 (3 H, s, $OCH_3$), 2.93-2.87 (2 H, m, $CH_2$), 2.80-2.73 (2 H, m, $CH_2$). Selected $^1$H NMR peaks from product mixture corresponding to the unidentified compound: $\delta_H$(400 MHz; $CDCl_3$) 6.90 (1 H, s, ArC*H*), 5.99 (1 H, s, C*H*CO), 3.88 (3 H, s, $OCH_3$). No data were reported in the literature.

**3,8-Dimethoxyphenanthrene-1,4-dione 25 and 9,10-dihydro-3,8-dimethoxyphenanthrene-1,4-dione 26**

General procedure **B** was followed, using the bromide **16** (1.02 g, 2.91 mmol), to give a red solid (218 mg, 28%) as an inseparable mixture of diones **25** and **26** in an 82:18 ratio which was used without further purification. Selected $^1$H NMR data from product mixture corresponding to the dione **25**: $\delta_H$(400 MHz; $CDCl_3$) 9.10 (1 H, d, *J* 8.8, ArC*H*), 8.71 (1 H, d, *J* 8.8, ArC*H*), 8.18 (1 H, d, *J* 8.8, ArC*H*), 7.66 (1 H, br t, *J* 8.4, ArC*H*), 6.98 (1 H, d, *J* 7.7, ArC*H*), 6.15 (1 H, s, C*H*CO), 4.04 (3 H, s, $OCH_3$), 3.95 (3 H, s, $OCH_3$). All data were in agreement with the literature.[5] Selected $^1$H NMR data from product mixture corresponding to the dione **26**: $\delta_H$(400 MHz; $CDCl_3$) 5.94 (1 H, s, C*H*CO), 3.83 (3 H, s, $OCH_3$), 3.80 (3 H, s, $OCH_3$), 2.88-2.82 (2 H, m, $CH_2$), 2.77-2.72 (2 H, m, $CH_2$). No data were reported in the literature.

**General representative procedure C for demethylation to form alcohols 27-31**

Aqueous NaOH (2 M, 50 mL $g^{-1}$, 40.0 mmol) was added to a mixture of the ethers **17-26** (378 mg, 1.50 mmol) in ethanol (50 mL $g^{-1}$), and the resulting mixture was stirred at reflux for 1 h. The solution was cooled to room temperature, acidified with aqueous HCl (1 M, 100 mL $g^{-1}$), water was added (50 mL $g^{-1}$), and the solution was extracted with ethyl acetate (3 × 63 mL $g^{-}$

$^{1}$). The organic extracts were combined, washed with brine (250 mL $g^{-1}$), dried ($MgSO_4$), filtered and concentrated *in vacuo* to give the alcohols **27-31**.

**3-Hydroxy-8-methylphenanthrene-1,4-dione 27**

General procedure **C** was followed, using the mixture of diones **17** and **18** (378 mg, 1.50 mmol) to give the alcohol **27** (347 mg, 97%) as a pale red solid that did not require further purification; mp 208-214 °C (lit.[2] mp 209-212 °C); $\delta_H$[400 MHz; $(CD_3)_2SO$] 11.67 (1 H, br s, ArO*H*), 9.32 (1 H, d, *J* 8.8, ArC*H*), 8.51 (1 H, d, *J* 8.8, ArC*H*), 8.12 (1 H, d, *J* 8.8, ArC*H*), 7.69 (1 H, dd, *J* 8.8, 7.0, ArC*H*), 7.57 (1 H, d, *J* 7.0, ArC*H*), 6.15 (1 H, s, C*H*CO), 2.72 (3 H, s, ArC*H*$_3$); $\delta_C$[100 MHz; $(CD_3)_2SO$] 185.5 (*C*=O), 184.3 (*C*=O), 160.6 (*C*), 135.6 (*C*), 135.1 (*C*), 132.6 (*C*), 132.1 (Ar*C*H), 130.4 (Ar*C*H), 130.1 (*C*), 129.4 (Ar*C*H), 125.8 (*C*), 125.2 (Ar*C*H), 121.7 (Ar*C*H), 108.4 (Ar*C*H), 19.9 (*C*H$_3$). All data were in general agreement with the literature.[2]

**3-Hydroxyphenanthrene-1,4-dione 28**

General procedure **C** was followed, using the mixture of diones **19** and **20** (222 mg, 0.931 mmol) to give the alcohol **28** (200 mg, 97%) as a dark orange solid which was used without further purification; mp 199-202 °C (lit.[6] 200 °C); $\delta_H$[400 MHz; $(CD_3)_2CO$] 9.83 (1 H, br s, O*H*), 9.58 (1 H, d, *J* 8.7, ArC*H*), 8.41 (1 H, d, *J* 8.7, ArC*H*), 8.19 (1 H, dd, *J* 8.7, 1.7, ArC*H*), 8.10 (1 H, d, *J* 8.2, ArC*H*), 7.86-7.80 (1 H, m, ArC*H*), 7.77-7.71 (1 H, m, ArC*H*), 6.25 (1 H, d, *J* 1.7, C*H*CO). All $^1$H NMR data were in broad agreement with the literature,[2] although precise values were slightly shifted due to the different solvent used for analysis.

**3-Hydroxy-8-(trifluoromethyl)phenanthrene-1,4-dione 29**

General procedure **C** was followed, using the mixture of diones **21** and **22** (500 mg) to give the crude alcohol **29** (447 mg) as an orange solid which could not be purified further by either flash column chromatography or recrystallisation. Selected peaks from $^1$H NMR spectrum of crude material corresponding to the alcohol **29**: $\delta_H$[400 MHz; $(CD_3)_2CO$] 9.89 (1 H, d, *J* 9.0, ArC*H*), 8.64 (1 H, br d, *J* 9.0, ArC*H*), 8.40 (1 H, d, *J* 9.0, ArC*H*), 8.21 (1 H, d, *J* 7.3, ArC*H*), 8.00-7.94 (1 H, m, ArC*H*), 6.31 (1 H, s, C*H*CO). All $^1$H NMR data were in broad agreement with the literature,[2] although precise values were slightly shifted due to the different solvent used for analysis.

**8-Fluoro-3-hydroxyphenanthrene-1,4-dione 30**

General procedure **C** was followed, using the mixture of diones **23** and **24** (194 mg) to give the crude alcohol **30** (170 mg) as a dark red solid which could not be purified further by either flash column chromatography or recrystallisation. Selected peaks from $^{1}$H NMR spectrum of crude material corresponding to the alcohol **30**: $\delta_H$[400 MHz; $(CD_3)_2CO$] 9.95 (1 H, br s, O*H*), 9.40 (1 H, d, *J* 8.8, ArC*H*), 8.60 (1 H, d, *J* 8.8, ArC*H*), 8.29 (1 H, d, *J* 8.8, ArC*H*), 7.86-7.79 (1 H, m, ArC*H*), 7.51 (1 H, ddd, *J* 10.4, 7.8, 0.7, ArC*H*), 6.28 (1 H, s, C*H*CO). All $^{1}$H NMR data were in broad agreement with the literature,[2] although precise values were slightly shifted due to the different solvent used for analysis.

**3-Hydroxy-8-methoxyphenanthrene-1,4-dione 31**

General procedure **C** was followed, using the mixture of diones **25** and **26** (218 mg, 0.813 mmol) to give the alcohol **31** (192 mg, 93%) as a dark red solid which was used without further purification; mp 204-207 °C (lit.[2] 185-187 °C); $\nu_{max}$(NaCl discs)/cm$^{-1}$ 3281 (O-H), 1657 (C=O), 1634 (C=O), 1582 (C=C); $\delta_H$[400 MHz; $(CD_3)_2SO$] 11.65 (1 H, br s, O*H*), 8.97 (1 H, d, *J* 8.8, ArC*H*), 8.59 (1 H, d, *J* 8.8, ArC*H*), 8.02 (1 H, d, *J* 8.8, ArC*H*), 7.70 (1 H, t, *J* 7.8, ArC*H*), 7.15 (1 H, d, *J* 7.8, ArC*H*), 6.13 (1 H, s, C*H*CO), 4.00 (3 H, s, OC*H*$_3$). IR spectroscopy data were not reported in the literature. All other spectroscopic data were in agreement with the literature.[2]

#### Design of Experiments (DoE) studies

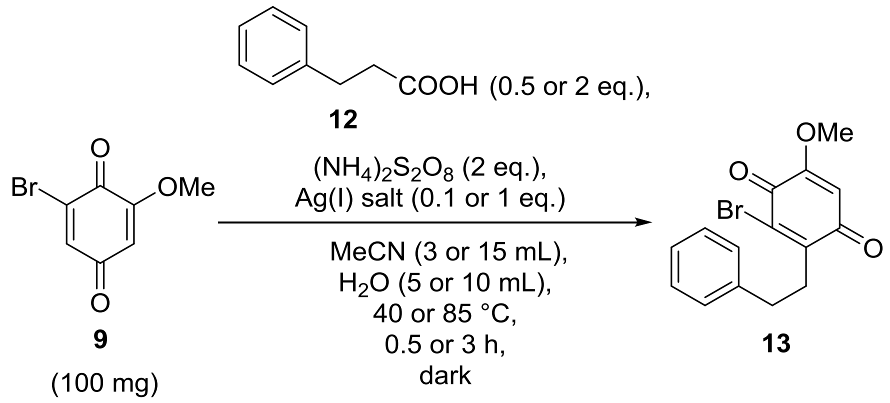


**Scheme 1**. General radical alkylation reaction for optimisation studies using DoE.

**Table 1**. The ten radical alkylation experiments carried out, as determined by the JMP DoE software.

| Entry | Eq. acid 12 | Eq. Ag(I) salt | Ag(I) salt | Vol. MeCN / mL | Vol. $H_2O$ / mL | Time / h | Temp. / °C | Metal needle used? | % yield product 13* |
|---|---|---|---|---|---|---|---|---|---|
| 1 | 0.5 | 1 | $Ag_3PO_4$ | 3 | 5 | 0.5 | 40 | No | 25% |
| 2 | 0.5 | 0.1 | $AgNO_3$ | 3 | 5 | 0.5 | 85 | Yes | 10% |
| 3 | 0.5 | 0.1 | $Ag_3PO_4$ | 15 | 10 | 0.5 | 85 | Yes | 11% |
| 4 | 0.5 | 1 | $AgNO_3$ | 3 | 10 | 3 | 40 | Yes | 18% |
| 5 | 0.5 | 1 | $Ag_2CO_3$ | 15 | 5 | 3 | 85 | No | 26% |
| 6 | 2 | 0.1 | $Ag_2CO_3$ | 3 | 10 | 0.5 | 40 | No | 6% |
| 7 | 2 | 1 | $Ag_2CO_3$ | 15 | 5 | 0.5 | 40 | Yes | 0% |
| 8 | 2 | 1 | $AgNO_3$ | 15 | 10 | 0.5 | 85 | No | 51% |
| 9 | 2 | 0.1 | $AgNO_3$ | 15 | 5 | 3 | 40 | No | 18% |
| 10 | 2 | 1 | $Ag_3PO_4$ | 3 | 5 | 3 | 85 | Yes | 24% |

* Represents isolated product after chromatographic purification.

### NMR spectra of synthesised compounds

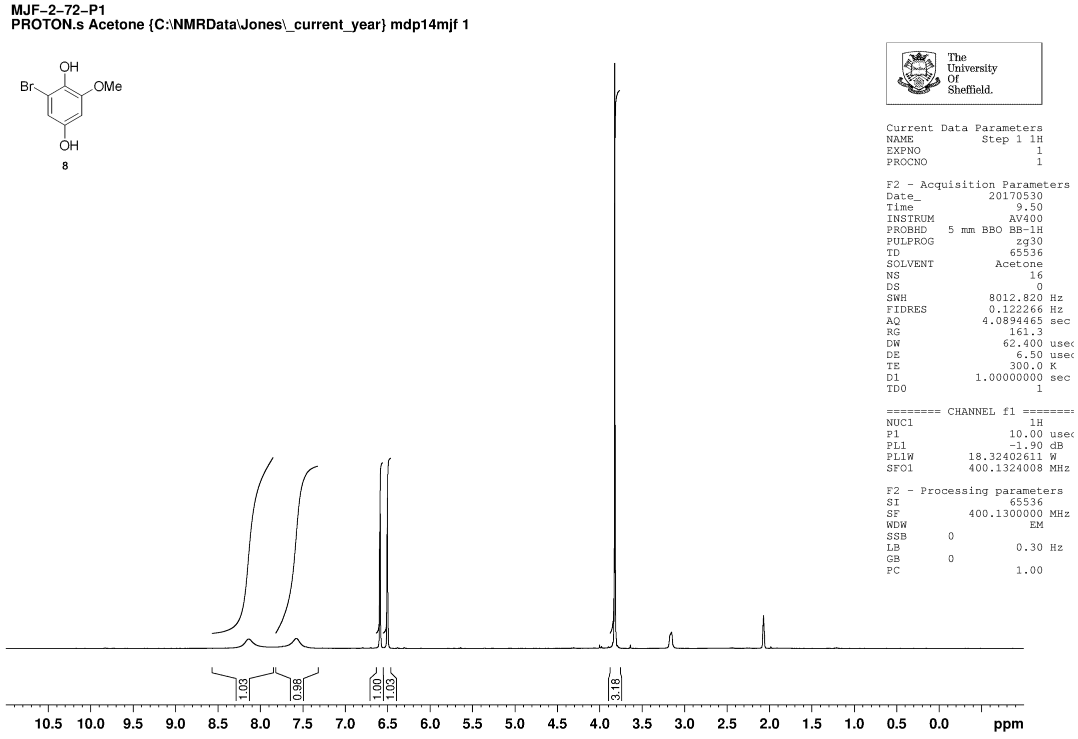


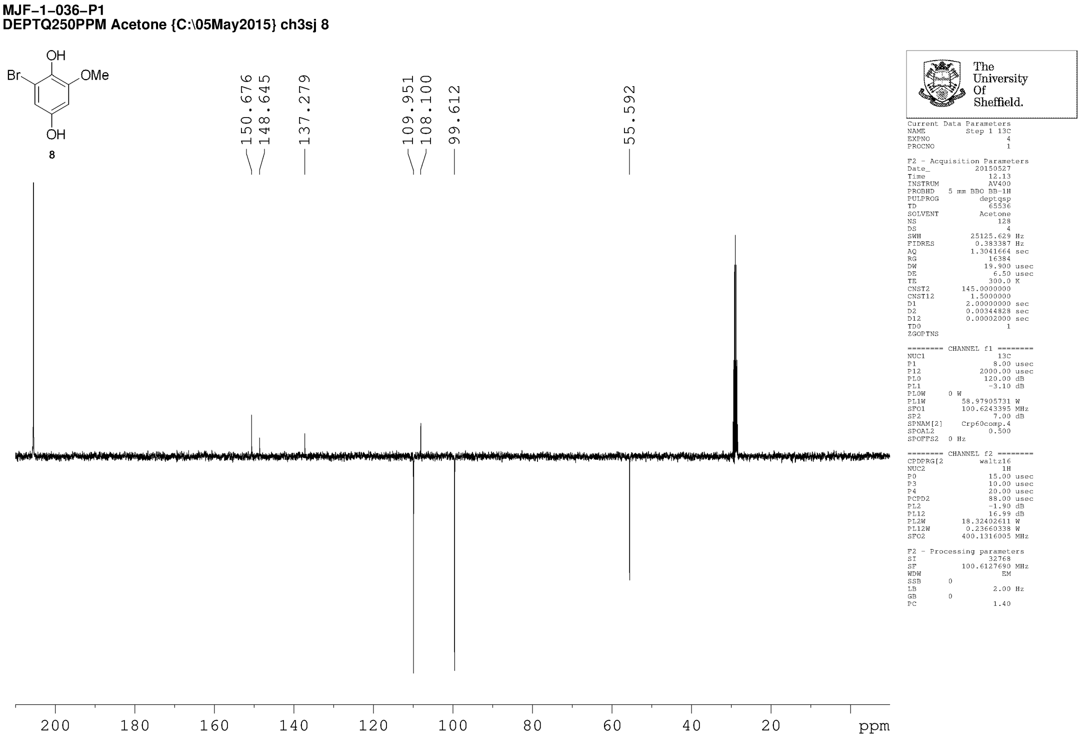

**MJF-2-73-P1**
**PROTON.s CDCl3 {C:\NMRData\Jones_current_year} mdp14mjf 9**

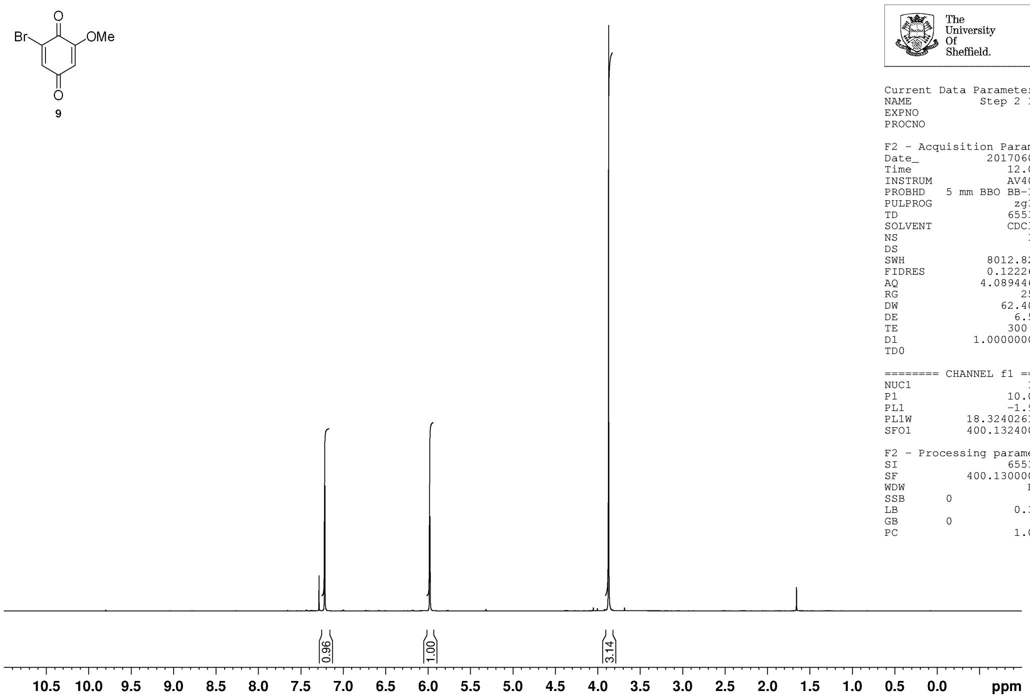


The University Of Sheffield.

| Current Data Parameters | |
|---|---|
| NAME | Step 2 1H |
| EXPNO | 1 |
| PROCNO | 1 |

| F2 - Acquisition Parameters | |
|---|---|
| Date_ | 20170601 |
| Time | 12.02 |
| INSTRUM | AV400 |
| PROBHD | 5 mm BBO BB-1H |
| PULPROG | zg30 |
| TD | 65536 |
| SOLVENT | CDCl3 |
| NS | 16 |
| DS | 0 |
| SWH | 8012.820 Hz |
| FIDRES | 0.122266 Hz |
| AQ | 4.0894465 sec |
| RG | 256 |
| DW | 62.400 usec |
| DE | 6.50 usec |
| TE | 300.0 K |
| D1 | 1.00000000 sec |
| TD0 | 1 |

| ======== CHANNEL f1 ======== | |
|---|---|
| NUC1 | 1H |
| P1 | 10.00 usec |
| PL1 | -1.90 dB |
| PL1W | 18.32402611 W |
| SFO1 | 400.1324008 MHz |

| F2 - Processing parameters | |
|---|---|
| SI | 65536 |
| SF | 400.1300000 MHz |
| WDW | EM |
| SSB | 0 |
| LB | 0.30 Hz |
| GB | 0 |
| PC | 1.00 |

**MJF-1-005-P1**
**JMOD250ppm CDCl3 {C:\NMRData\11Nov2014} ch3sj 25**

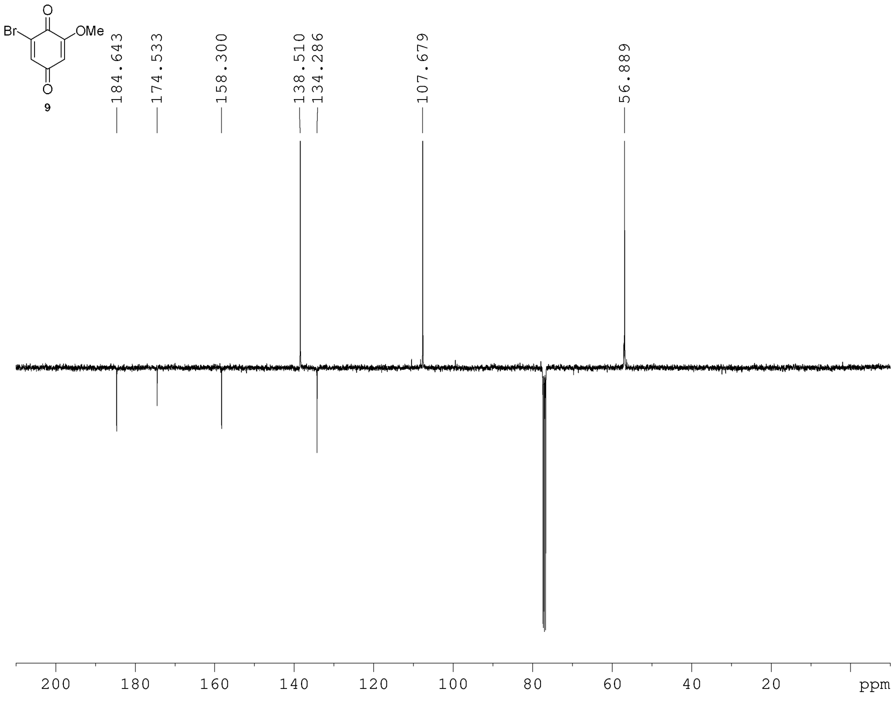


The University Of Sheffield.

| Current Data Parameters | |
|---|---|
| NAME | Step 2 13C |
| EXPNO | 4 |
| PROCNO | 1 |

| F2 - Acquisition Parameters | |
|---|---|
| Date_ | 20141104 |
| Time | 23.52 |
| INSTRUM | AVIIIHD400 |
| PROBHD | 5 mm PABBO BB/ |
| PULPROG | jmod |
| TD | 65536 |
| SOLVENT | CDCl3 |
| NS | 512 |
| DS | 4 |
| SWH | 25252.525 Hz |
| FIDRES | 0.385323 Hz |
| AQ | 1.2976128 sec |
| RG | 195.75 |
| DW | 19.800 usec |
| DE | 6.50 usec |
| TE | 298.0 K |
| CNST2 | 145.0000000 |
| CNST11 | 1.0000000 |
| D1 | 3.70000005 sec |
| D20 | 0.00689655 sec |
| TD0 | 1 |

| ======== CHANNEL f1 ======== | |
|---|---|
| SFO1 | 100.6494869 MHz |
| NUC1 | 13C |
| P1 | 10.00 usec |
| P2 | 20.00 usec |
| PLW1 | 71.00000000 W |

| ======== CHANNEL f2 ======== | |
|---|---|
| SFO2 | 400.2316009 MHz |
| NUC2 | 1H |
| CPDPRG[2 | waltz16 |
| PCPD2 | 90.00 usec |
| PLW2 | 16.00000000 W |
| PLW12 | 0.19753000 W |

| F2 - Processing parameters | |
|---|---|
| SI | 32768 |
| SF | 100.6379135 MHz |
| WDW | EM |
| SSB | 0 |
| LB | 2.00 Hz |
| GB | 0 |
| PC | 1.40 |

**MJF-1-019-P1**
**PRO CDCl3 {C:\02Feb2015} ch3sj 29**

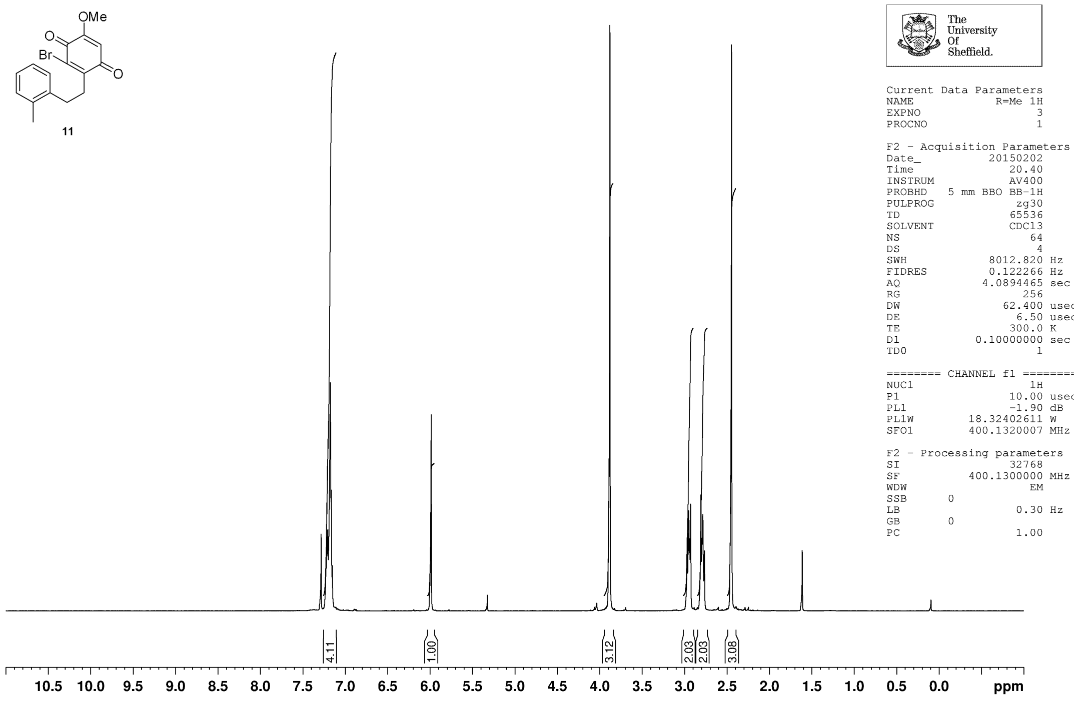


**MJF-1-008-P2**
**JMOD250ppm CDCl3 {C:\NMRData\11Nov2014} ch3sj 44**

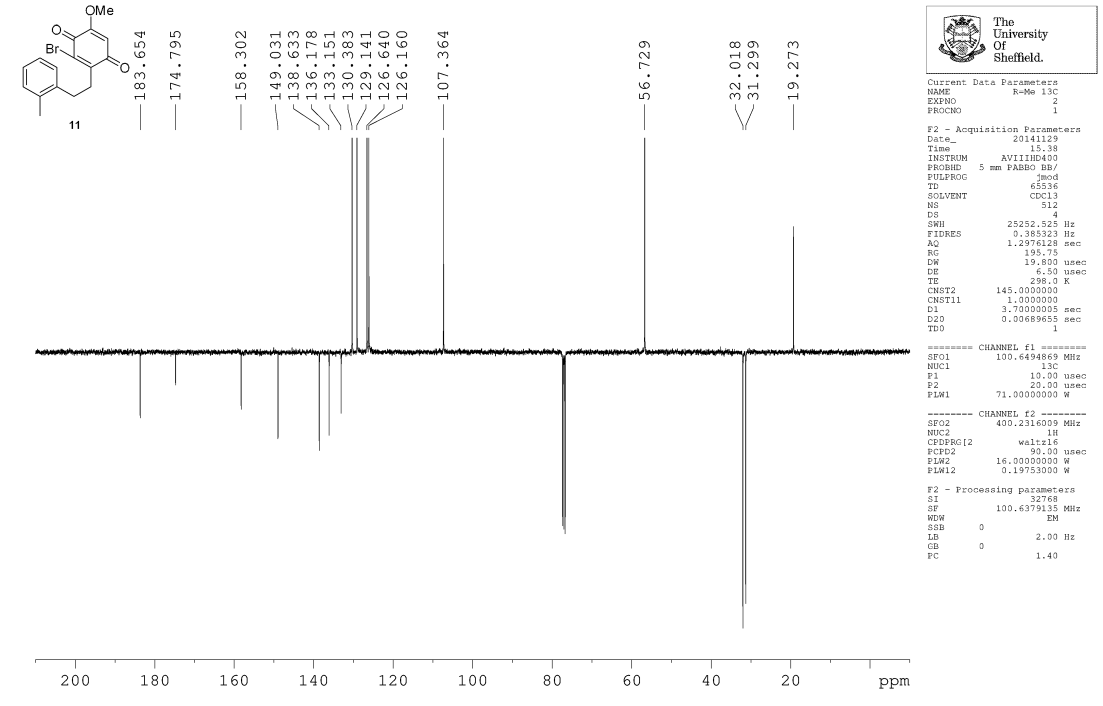

**MJF-1-038-P1**
**PRO CDCl3 {C:\05May2015} ch3sj 48**

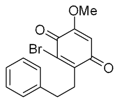


13

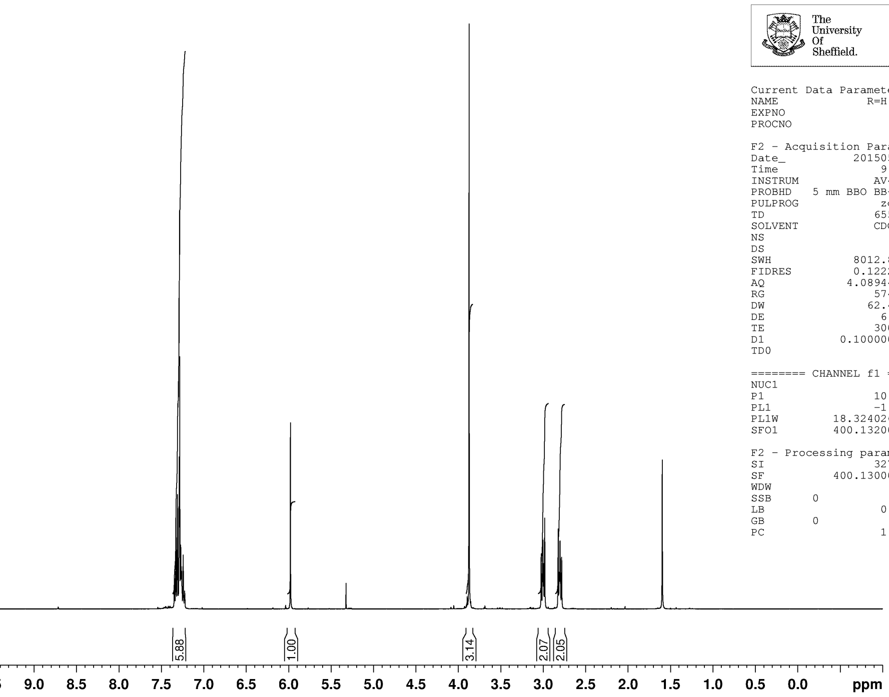


The University Of Sheffield.

```
Current Data Parameters
NAME              R=H 1H
EXPNO                  4
PROCNO                 1

F2 - Acquisition Parameters
Date_           20150527
Time                9.38
INSTRUM            AV400
PROBHD   5 mm BBO BB-1H
PULPROG             zg30
TD                 65536
SOLVENT            CDCl3
NS                    64
DS                     4
SWH             8012.820 Hz
FIDRES          0.122266 Hz
AQ             4.0894465 sec
RG                 574.7
DW                62.400 usec
DE                  6.50 usec
TE                 300.0 K
D1            0.10000000 sec
TD0                    1

======== CHANNEL f1 ========
NUC1                  1H
P1                 10.00 usec
PL1                -1.90 dB
PL1W         18.32402611 W
SFO1         400.1320007 MHz

F2 - Processing parameters
SI                 32768
SF           400.1300000 MHz
WDW                   EM
SSB      0
LB                  0.30 Hz
GB       0
PC                  1.00
```

**MJF-1-038-P1**
**DEPTQ250PPM CDCl3 {C:\05May2015} ch3sj 10**

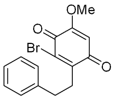


13

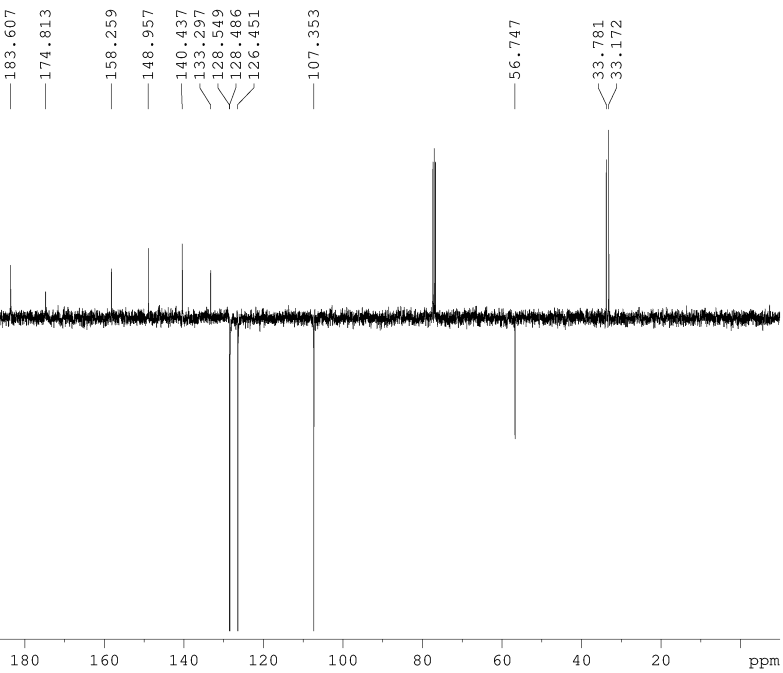


The University Of Sheffield.

```
Current Data Parameters
NAME             R=H 13C
EXPNO                  6
PROCNO                 1

F2 - Acquisition Parameters
Date_           20150527
Time               12.46
INSTRUM            AV400
PROBHD   5 mm BBO BB-1H
PULPROG          deptqsp
TD                 65536
SOLVENT            CDCl3
NS                   128
DS                     4
SWH            25125.629 Hz
FIDRES          0.383387 Hz
AQ             1.3041664 sec
RG                 16384
DW                19.900 usec
DE                  6.50 usec
TE                 300.0 K
CNST2        145.0000000
CNST12         1.5000000
D1            2.00000000 sec
D2            0.00344828 sec
D12           0.00002000 sec
TD0                    1
ZGOPTNS

======== CHANNEL f1 ========
NUC1                 13C
P1                  8.00 usec
P12              2000.00 usec
PL0               120.00 dB
PL1                -3.10 dB
PL0W      0 W
PL1W         58.97905731 W
SFO1         100.6243395 MHz
SP2                 7.00 dB
SPNAM[2]     Crp60comp.4
SPOAL2             0.500
SPOFFS2   0 Hz

======== CHANNEL f2 ========
CPDPRG[2         waltz16
NUC2                  1H
P0                 15.00 usec
P3                 10.00 usec
P4                 20.00 usec
PCPD2              88.00 usec
PL2                -1.90 dB
PL12               16.99 dB
PL2W         18.32402611 W
PL12W         0.23660338 W
SFO2         400.1316005 MHz

F2 - Processing parameters
SI                 32768
SF           100.6127690 MHz
WDW                   EM
SSB      0
LB                  2.00 Hz
GB       0
PC                  1.40
```

**MJF-1-067-P1**
**PRO CDCl3 {C:\01Jan2016} ch3sj 21**

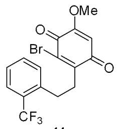


**14**

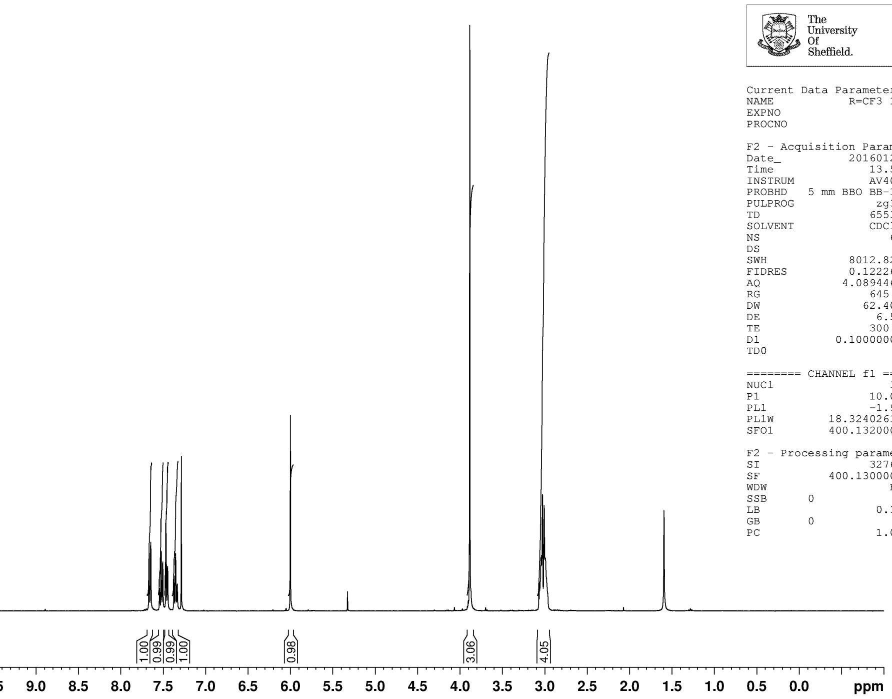


The University Of Sheffield.

```
Current Data Parameters
NAME             R=CF3 1H
EXPNO                   1
PROCNO                  1

F2 - Acquisition Parameters
Date_            20160126
Time                13.55
INSTRUM             AV400
PROBHD   5 mm BBO BB-1H
PULPROG              zg30
TD                  65536
SOLVENT             CDCl3
NS                     64
DS                      4
SWH            8012.820 Hz
FIDRES         0.122266 Hz
AQ            4.0894465 sec
RG                  645.1
DW               62.400 usec
DE                 6.50 usec
TE                300.0 K
D1           0.10000000 sec
TD0                     1

======== CHANNEL f1 ========
NUC1                   1H
P1                10.00 usec
PL1               -1.90 dB
PL1W        18.32402611 W
SFO1        400.1320007 MHz

F2 - Processing parameters
SI                  32768
SF          400.1300000 MHz
WDW                    EM
SSB      0
LB                 0.30 Hz
GB       0
PC                   1.00
```

**MJF-1-067-P1**
**JMOD250PPM CDCl3 {C:\01Jan2016} ch3sj 40**

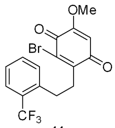


**14**

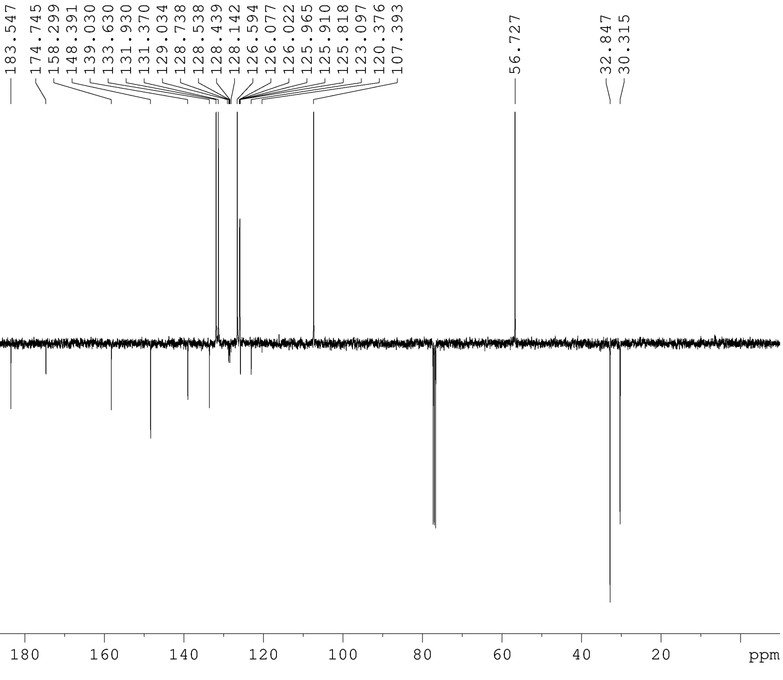


The University Of Sheffield.

```
Current Data Parameters
NAME             R=CF3 13C
EXPNO                   6
PROCNO                  1

F2 - Acquisition Parameters
Date_            20160126
Time                18.27
INSTRUM             AV400
PROBHD   5 mm BBO BB-1H
PULPROG            jmodsp
TD                  65536
SOLVENT             CDCl3
NS                    512
DS                      4
SWH           25125.629 Hz
FIDRES         0.383387 Hz
AQ            1.3041664 sec
RG                  16384
DW               19.900 usec
DE                 6.50 usec
TE                300.0 K
CNST2       145.0000000
CNST11        1.0000000
D1           4.00000000 sec
D20          0.00689655 sec
P2                16.00 usec
TD0                     1

======== CHANNEL f1 ========
NUC1                  13C
P1                 8.00 usec
P12             2000.00 usec
PL1               -3.10 dB
PL1W        58.97905731 W
SFO1        100.6243395 MHz
SP2                7.00 dB
SPNAM[2]     Crp60comp.4
SPOAL2              0.500
SPOFFS2  0 Hz

======== CHANNEL f2 ========
CPDPRG[2          waltz16
NUC2                   1H
PCPD2             88.00 usec
PL2               -1.90 dB
PL12              16.99 dB
PL2W        18.32402611 W
PL12W        0.23660338 W
SFO2        400.1316005 MHz

F2 - Processing parameters
SI                  32768
SF          100.6127690 MHz
WDW                    EM
SSB      0
LB                 2.00 Hz
GB       0
PC                   1.40
```

**MJF−2−27−P1**
**PRO CDCl3 {C:\07Jul2016} ch3sj 2**

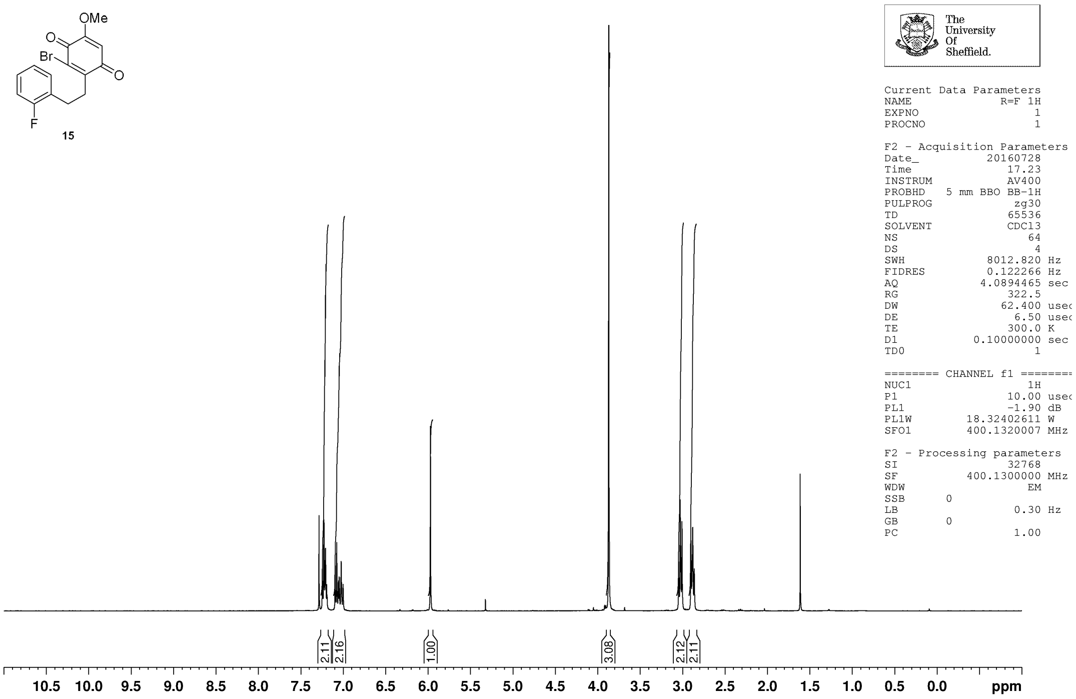


**MJF−1−66−P1**
**JMOD250ppm CDCl3 {C:\NMRData\01Jan2016} ch3sj 22**

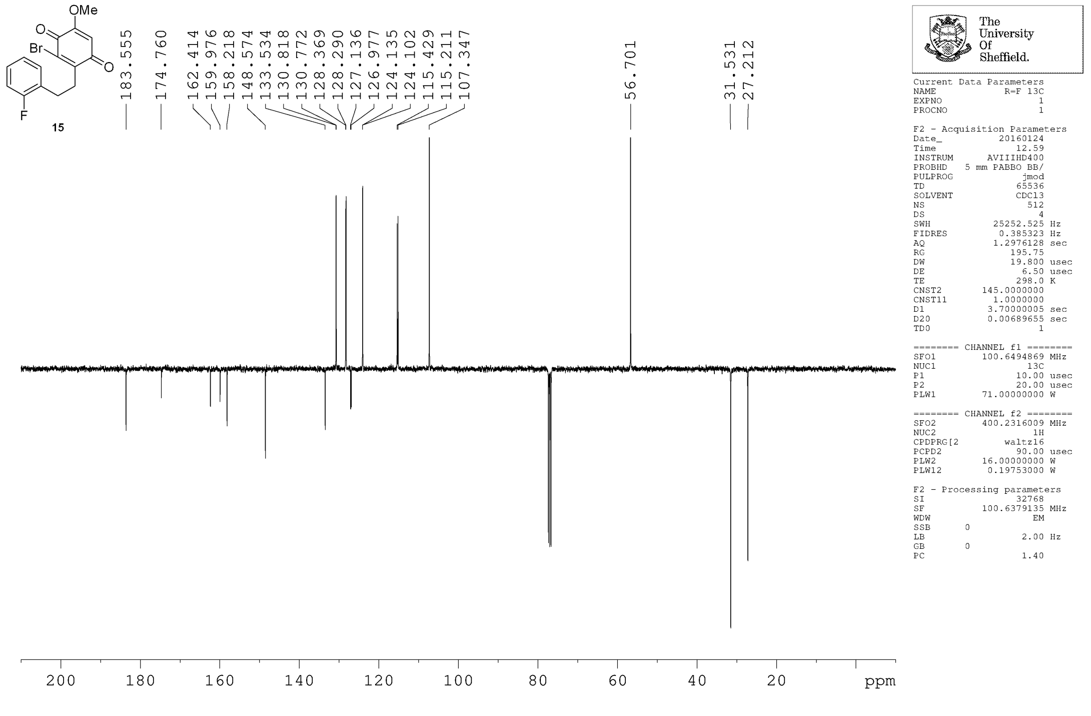

**MJF-1-79-P2**
**PRO CDCl3 {C:\02Feb2016} ch3sj 28**

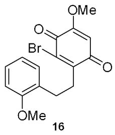


16

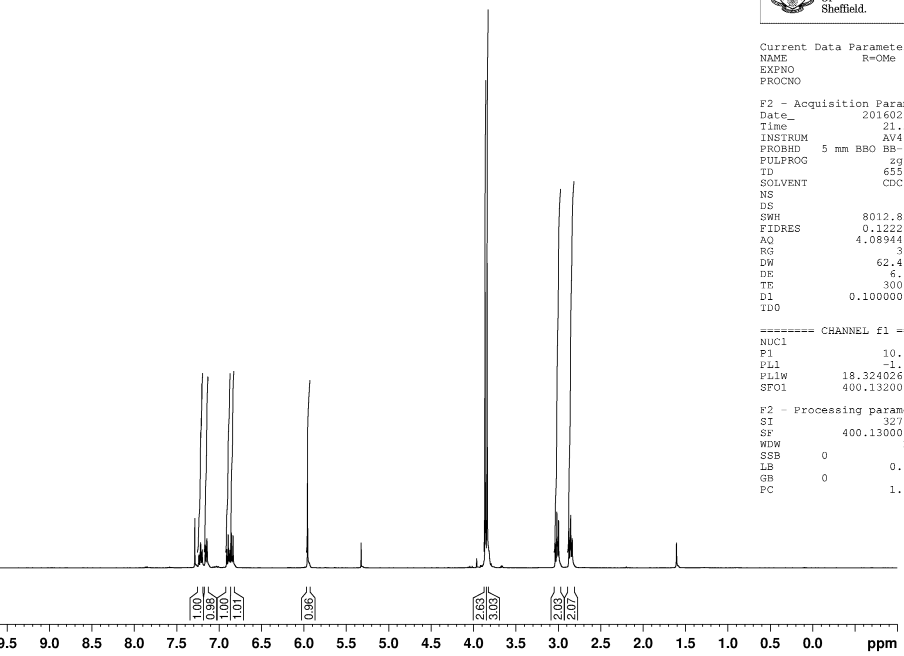


The University Of Sheffield.

Current Data Parameters
NAME R=OMe 1H
EXPNO 3
PROCNO 1

F2 - Acquisition Parameters
Date_ 20160212
Time 21.20
INSTRUM AV400
PROBHD 5 mm BBO BB-1H
PULPROG zg30
TD 65536
SOLVENT CDCl3
NS 64
DS 4
SWH 8012.820 Hz
FIDRES 0.122266 Hz
AQ 4.0894465 sec
RG 362
DW 62.400 usec
DE 6.50 usec
TE 300.0 K
D1 0.10000000 sec
TD0 1

======== CHANNEL f1 ========
NUC1 1H
P1 10.00 usec
PL1 -1.90 dB
PL1W 18.32402611 W
SFO1 400.1320007 MHz

F2 - Processing parameters
SI 32768
SF 400.1300000 MHz
WDW EM
SSB 0
LB 0.30 Hz
GB 0
PC 1.00

**MJF-1-79-P2**
**JMOD250PPM CDCl3 {C:\02Feb2016} ch3sj 8**

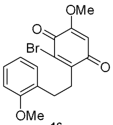


16

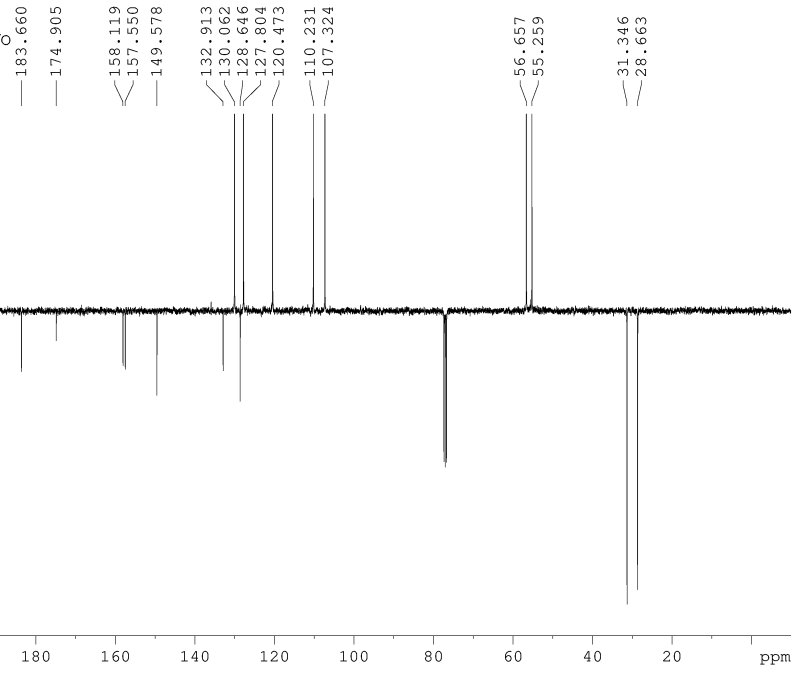


The University Of Sheffield.

Current Data Parameters
NAME R=OMe 13C
EXPNO 3
PROCNO 1

F2 - Acquisition Parameters
Date_ 20160214
Time 19.06
INSTRUM AV400
PROBHD 5 mm BBO BB-1H
PULPROG jmodsp
TD 65536
SOLVENT CDCl3
NS 512
DS 4
SWH 25125.629 Hz
FIDRES 0.383387 Hz
AQ 1.3041664 sec
RG 16384
DW 19.900 usec
DE 6.50 usec
TE 300.0 K
CNST2 145.0000000
CNST11 1.0000000
D1 4.00000000 sec
D20 0.00689655 sec
P2 16.00 usec
TD0 1

======== CHANNEL f1 ========
NUC1 13C
P1 8.00 usec
P12 2000.00 usec
PL1 -3.10 dB
PL1W 58.97905731 W
SFO1 100.6243395 MHz
SP2 7.00 dB
SPNAM[2] Crp60comp.4
SPOAL2 0.500
SPOFFS2 0 Hz

======== CHANNEL f2 ========
CPDPRG[2 waltz16
NUC2 1H
PCPD2 88.00 usec
PL2 -1.90 dB
PL12 16.99 dB
PL2W 18.32402611 W
PL12W 0.23660338 W
SFO2 400.1316005 MHz

F2 - Processing parameters
SI 32768
SF 100.6127690 MHz
WDW EM
SSB 0
LB 2.00 Hz
GB 0
PC 1.40

**MJF-2-30-P1**
**PRO CDCl3 {C:\08Aug2016} ch3sj 27**

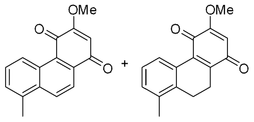


**17** **18**

*Partially purified material taken on directly to next step*

The University Of Sheffield.

```
Current Data Parameters
NAME              R=Me 1H
EXPNO                   3
PROCNO                  1

F2 - Acquisition Parameters
Date_            20160802
Time                19.08
INSTRUM             AV400
PROBHD   5 mm BBO BB-1H
PULPROG              zg30
TD                  65536
SOLVENT             CDCl3
NS                     64
DS                      4
SWH              8012.820 Hz
FIDRES           0.122266 Hz
AQ              4.0894465 sec
RG                  456.1
DW                 62.400 usec
DE                   6.50 usec
TE                  300.0 K
D1             0.10000000 sec
TD0                     1

======== CHANNEL f1 ========
NUC1                   1H
P1                  10.00 usec
PL1                 -1.90 dB
PL1W          18.32402611 W
SFO1          400.1320007 MHz

F2 - Processing parameters
SI                  32768
SF            400.1300000 MHz
WDW                    EM
SSB      0
LB                   0.30 Hz
GB       0
PC                   1.00
```

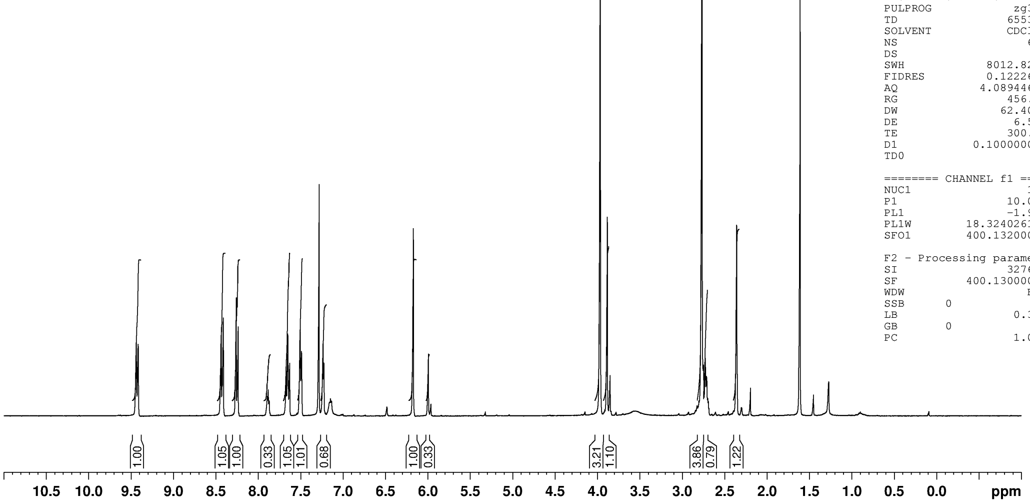


**MJF-2-23-P1**
**PRO CDCl3 {C:\07Jul2016} ch3sj 30**

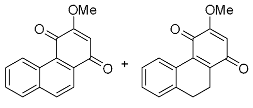


**19** **20**

*Partially purified material taken on directly to next step*

The University Of Sheffield.

```
Current Data Parameters
NAME               R=H 1H
EXPNO                   3
PROCNO                  1

F2 - Acquisition Parameters
Date_            20160726
Time                16.34
INSTRUM             AV400
PROBHD   5 mm BBO BB-1H
PULPROG              zg30
TD                  65536
SOLVENT             CDCl3
NS                     64
DS                      4
SWH              8012.820 Hz
FIDRES           0.122266 Hz
AQ              4.0894465 sec
RG                    362
DW                 62.400 usec
DE                   6.50 usec
TE                  300.0 K
D1             0.10000000 sec
TD0                     1

======== CHANNEL f1 ========
NUC1                   1H
P1                  10.00 usec
PL1                 -1.90 dB
PL1W          18.32402611 W
SFO1          400.1320007 MHz

F2 - Processing parameters
SI                  32768
SF            400.1300000 MHz
WDW                    EM
SSB      0
LB                   0.30 Hz
GB       0
PC                   1.00
```

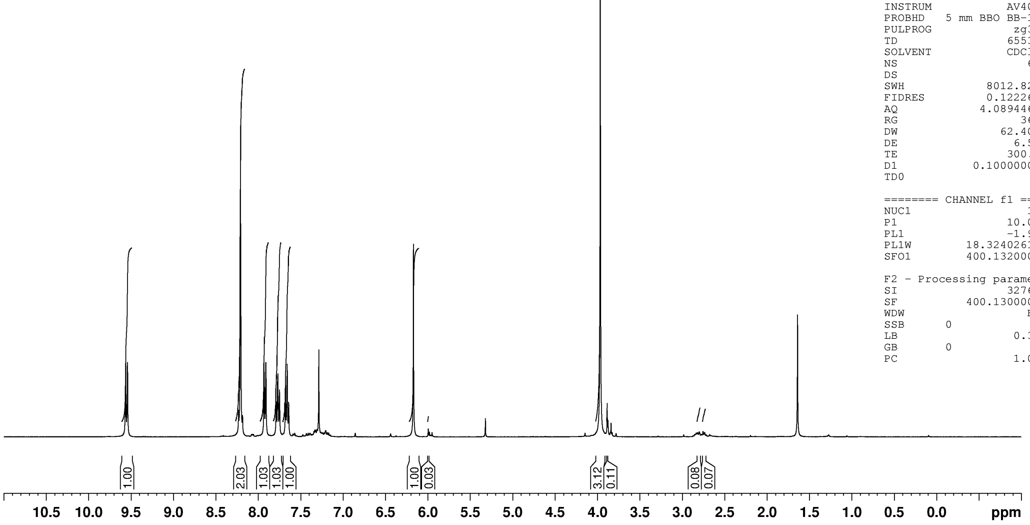

**MJF-2-37-P1**
**PRO CDCl3 {C:\08Aug2016} ch3sj 14**

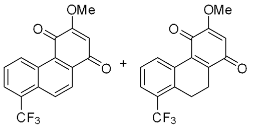

**21** + **22**

*Partially purified material taken on directly to next step*

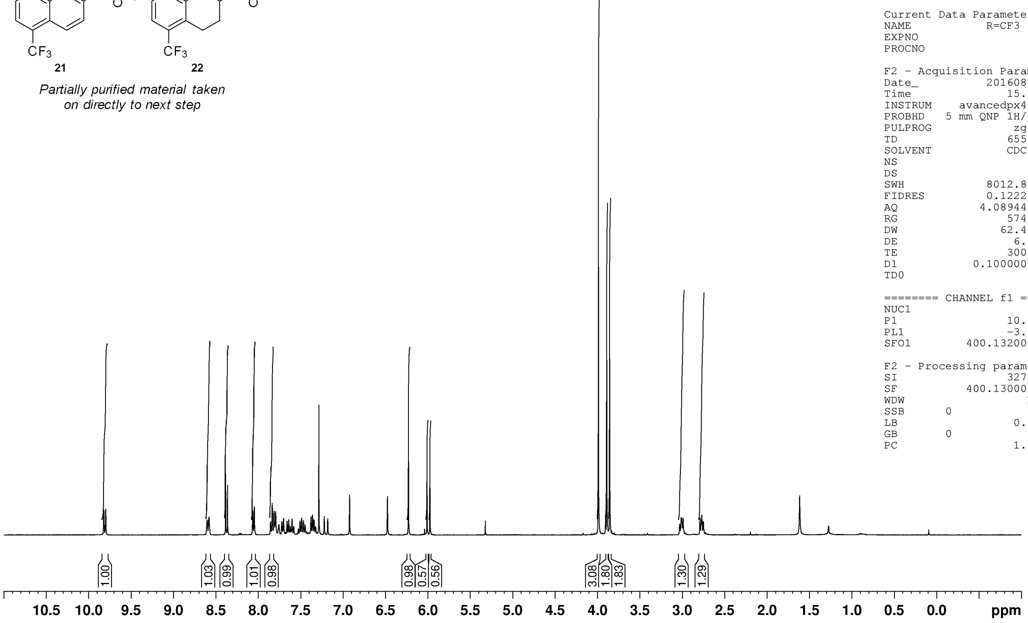


The University Of Sheffield.

```
Current Data Parameters
NAME            R=CF3 1H
EXPNO                  1
PROCNO                 1

F2 - Acquisition Parameters
Date_           20160816
Time               15.36
INSTRUM     avancedpx400
PROBHD   5 mm QNP 1H/13
PULPROG             zg30
TD                 65536
SOLVENT            CDCl3
NS                    64
DS                     2
SWH             8012.820 Hz
FIDRES          0.122266 Hz
AQ             4.0894465 sec
RG                 574.7
DW                62.400 usec
DE                  6.50 usec
TE                 300.0 K
D1            0.10000000 sec
TD0                    1

======== CHANNEL f1 ========
NUC1                  1H
P1                 10.50 usec
PL1                -3.00 dB
SFO1         400.1320007 MHz

F2 - Processing parameters
SI                 32768
SF           400.1300000 MHz
WDW                   EM
SSB      0
LB                  0.30 Hz
GB       0
PC                  1.00
```

**MJF-2-33-P1**
**PRO CDCl3 {C:\08Aug2016} ch3sj 36**

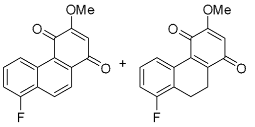

**23** + **24**

*Partially purified material taken on directly to next step*

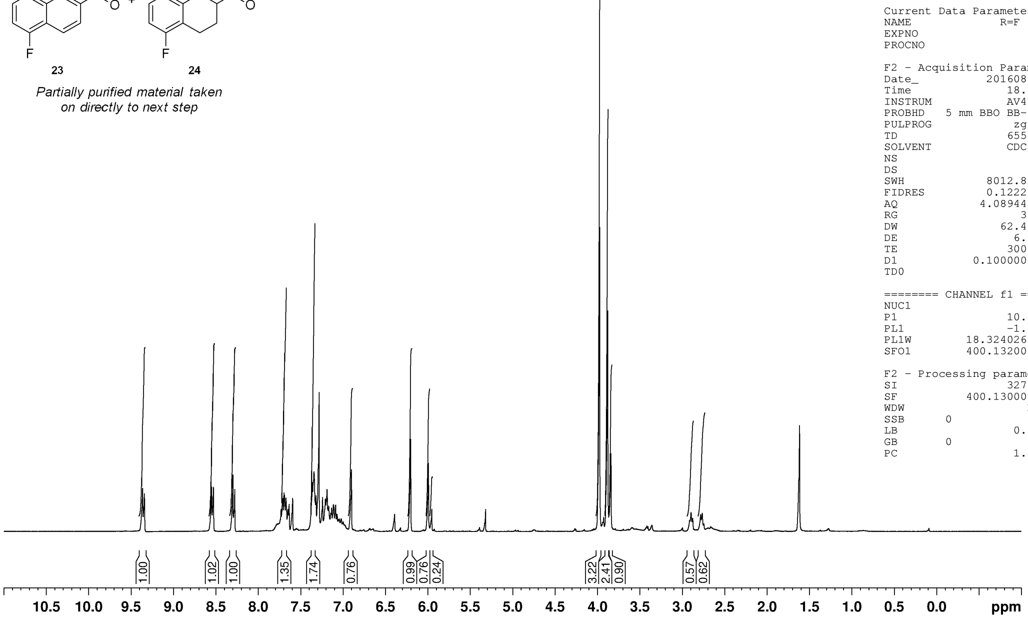


The University Of Sheffield.

```
Current Data Parameters
NAME              R=F 1H
EXPNO                  5
PROCNO                 1

F2 - Acquisition Parameters
Date_           20160809
Time               18.13
INSTRUM            AV400
PROBHD   5 mm BBO BB-1H
PULPROG             zg30
TD                 65536
SOLVENT            CDCl3
NS                    64
DS                     4
SWH             8012.820 Hz
FIDRES          0.122266 Hz
AQ             4.0894465 sec
RG                   362
DW                62.400 usec
DE                  6.50 usec
TE                 300.0 K
D1            0.10000000 sec
TD0                    1

======== CHANNEL f1 ========
NUC1                  1H
P1                 10.00 usec
PL1                -1.90 dB
PL1W         18.32402611 W
SFO1         400.1320007 MHz

F2 - Processing parameters
SI                 32768
SF           400.1300000 MHz
WDW                   EM
SSB      0
LB                  0.30 Hz
GB       0
PC                  1.00
```

**MJF–2–40–P1**
**PRO CDCl3 {C:\09Sep2016} ch3sj 26**

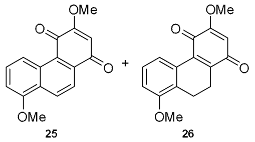


**25** **26**

*Partially purified material taken on directly to next step*

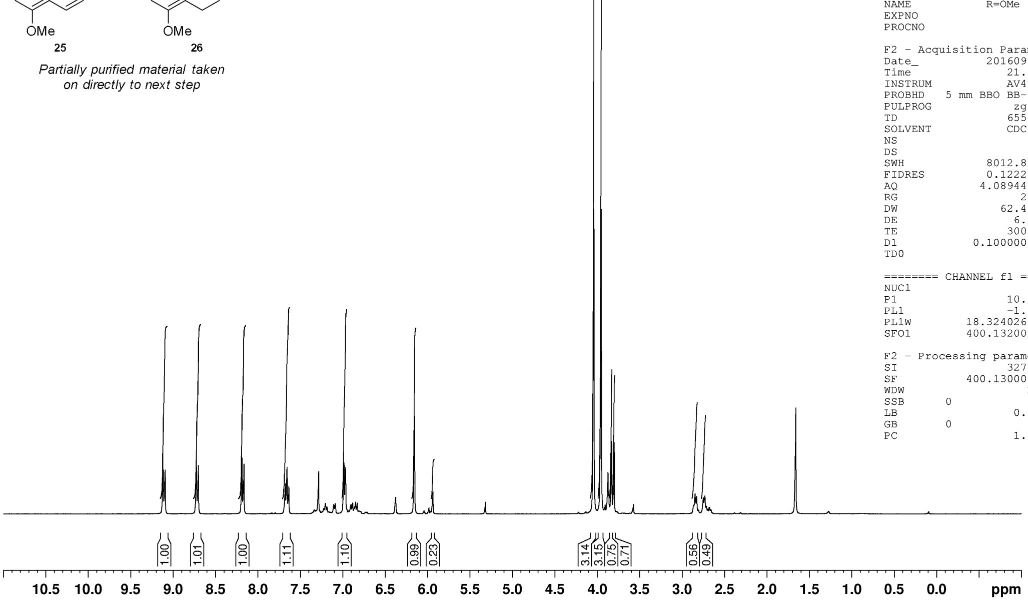


The University Of Sheffield.

```
Current Data Parameters
NAME              R=OMe 1H
EXPNO                    3
PROCNO                   1

F2 - Acquisition Parameters
Date_             20160905
Time                 21.18
INSTRUM              AV400
PROBHD   5 mm BBO BB-1H
PULPROG               zg30
TD                   65536
SOLVENT              CDCl3
NS                      64
DS                       4
SWH            8012.820 Hz
FIDRES         0.122266 Hz
AQ            4.0894465 sec
RG                     256
DW              62.400 usec
DE               6.50 usec
TE                300.0 K
D1          0.10000000 sec
TD0                      1

======== CHANNEL f1 ========
NUC1                    1H
P1              10.00 usec
PL1              -1.90 dB
PL1W       18.32402611 W
SFO1      400.1320007 MHz

F2 - Processing parameters
SI                   32768
SF        400.1300000 MHz
WDW                     EM
SSB      0
LB                0.30 Hz
GB       0
PC                    1.00
```

**MJF-1-062-C2**
**PRO DMSO {C:\10Oct2015} ch3sj 45**

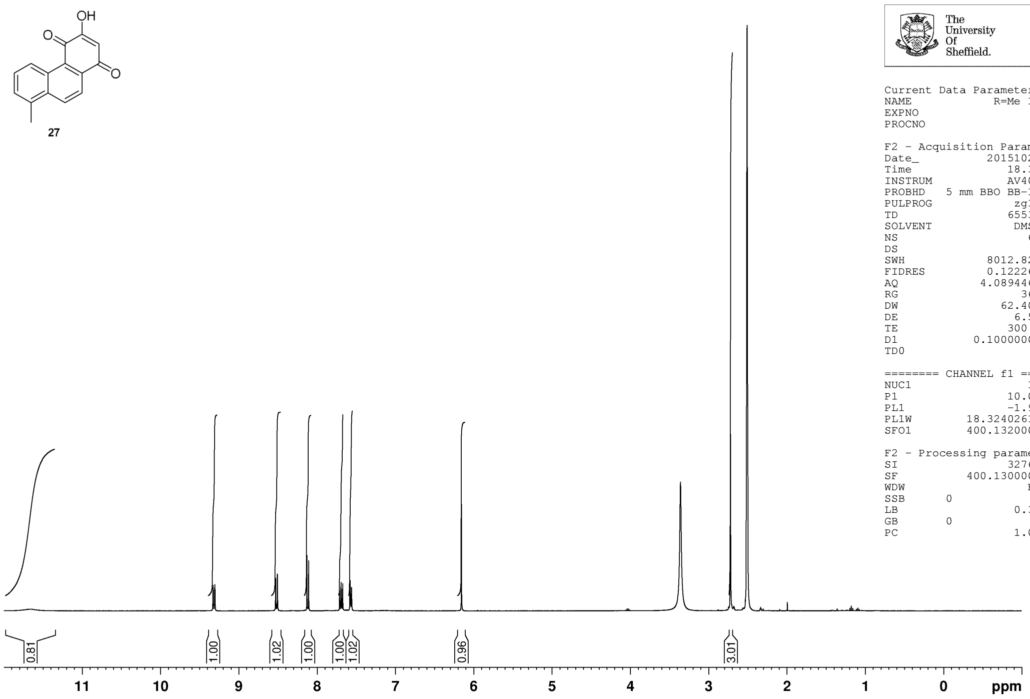


The University Of Sheffield.

```
Current Data Parameters
NAME              R=Me 1H
EXPNO                   3
PROCNO                  1

F2 - Acquisition Parameters
Date_            20151022
Time                18.30
INSTRUM             AV400
PROBHD   5 mm BBO BB-1H
PULPROG              zg30
TD                  65536
SOLVENT              DMSO
NS                     64
DS                      4
SWH              8012.820 Hz
FIDRES           0.122266 Hz
AQ              4.0894465 sec
RG                    362
DW                 62.400 usec
DE                   6.50 usec
TE                  300.0 K
D1             0.10000000 sec
TD0                     1

======== CHANNEL f1 ========
NUC1                   1H
P1                  10.00 usec
PL1                 -1.90 dB
PL1W          18.32402611 W
SFO1          400.1320007 MHz

F2 - Processing parameters
SI                  32768
SF            400.1300000 MHz
WDW                    EM
SSB      0
LB                   0.30 Hz
GB       0
PC                   1.00
```

**MJF-1-062-C2**
**JMOD250PPM DMSO {C:\10Oct2015} ch3sj 8**

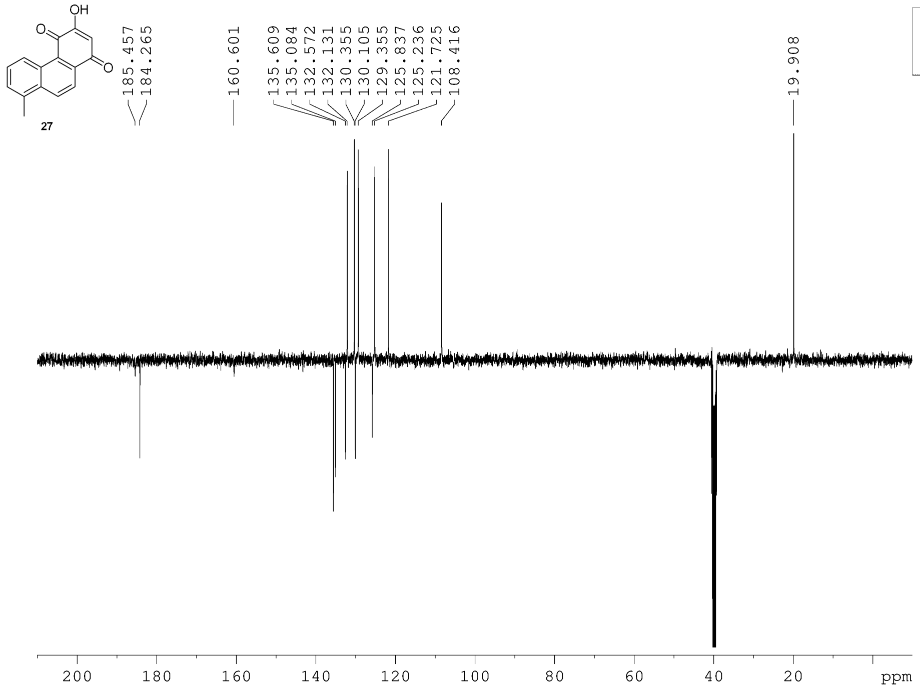


The University Of Sheffield.

```
Current Data Parameters
NAME             R=Me 13C
EXPNO                   1
PROCNO                  1

F2 - Acquisition Parameters
Date_            20151024
Time                14.56
INSTRUM             AV400
PROBHD   5 mm BBO BB-1H
PULPROG            jmodsp
TD                  65536
SOLVENT              DMSO
NS                    512
DS                      4
SWH             25125.629 Hz
FIDRES           0.383387 Hz
AQ              1.3041664 sec
RG                  16384
DW                 19.900 usec
DE                   6.50 usec
TE                  300.0 K
CNST2         145.0000000
CNST11          1.0000000
D1             4.00000000 sec
D20            0.00689655 sec
P2                  16.00 usec
TD0                     1

======== CHANNEL f1 ========
NUC1                  13C
P1                   8.00 usec
P12               2000.00 usec
PL1                 -3.10 dB
PL1W          58.97905731 W
SFO1          100.6243395 MHz
SP2                  7.00 dB
SPNAM[2]    Crp60comp.4
SPOAL2              0.500
SPOFFS2  0 Hz

======== CHANNEL f2 ========
CPDPRG[2          waltz16
NUC2                   1H
PCPD2               88.00 usec
PL2                 -1.90 dB
PL12                16.99 dB
PL2W          18.32402611 W
PL12W          0.23660338 W
SFO2          400.1316005 MHz

F2 - Processing parameters
SI                  32768
SF            100.6127690 MHz
WDW                    EM
SSB      0
LB                   2.00 Hz
GB       0
PC                   1.40
```

**MJF-2-35-P2**
**PRO Acetone {C:\09Sep2016} ch3sj 17**

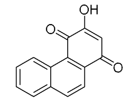

**28**

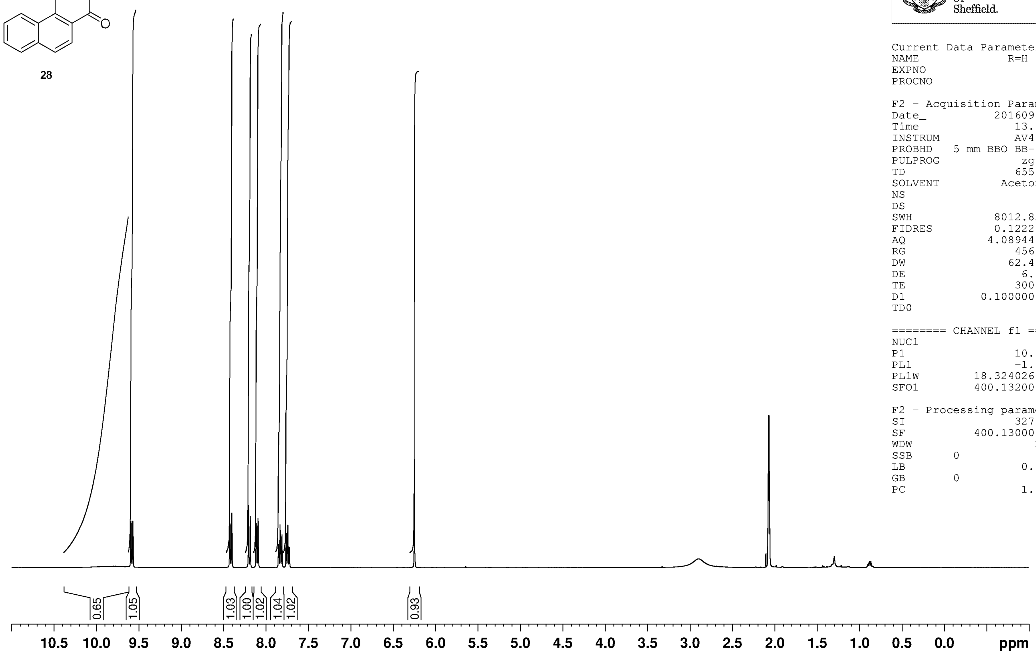


The University Of Sheffield.

```
Current Data Parameters
NAME              R=H 1H
EXPNO                  5
PROCNO                 1

F2 - Acquisition Parameters
Date_           20160916
Time               13.02
INSTRUM            AV400
PROBHD   5 mm BBO BB-1H
PULPROG             zg30
TD                 65536
SOLVENT          Acetone
NS                    64
DS                     4
SWH             8012.820 Hz
FIDRES          0.122266 Hz
AQ             4.0894465 sec
RG                 456.1
DW                62.400 usec
DE                  6.50 usec
TE                 300.0 K
D1            0.10000000 sec
TD0                    1

======== CHANNEL f1 ========
NUC1                  1H
P1                 10.00 usec
PL1                -1.90 dB
PL1W         18.32402611 W
SFO1         400.1320007 MHz

F2 - Processing parameters
SI                 32768
SF           400.1300000 MHz
WDW                   EM
SSB      0
LB                  0.30 Hz
GB       0
PC                  1.00
```

**MJF-1-076-P1**
**PRO Acetone {C:\02Feb2016} ch3sj 1**

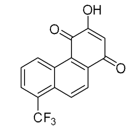

**29**
*Crude material taken on directly to next step*

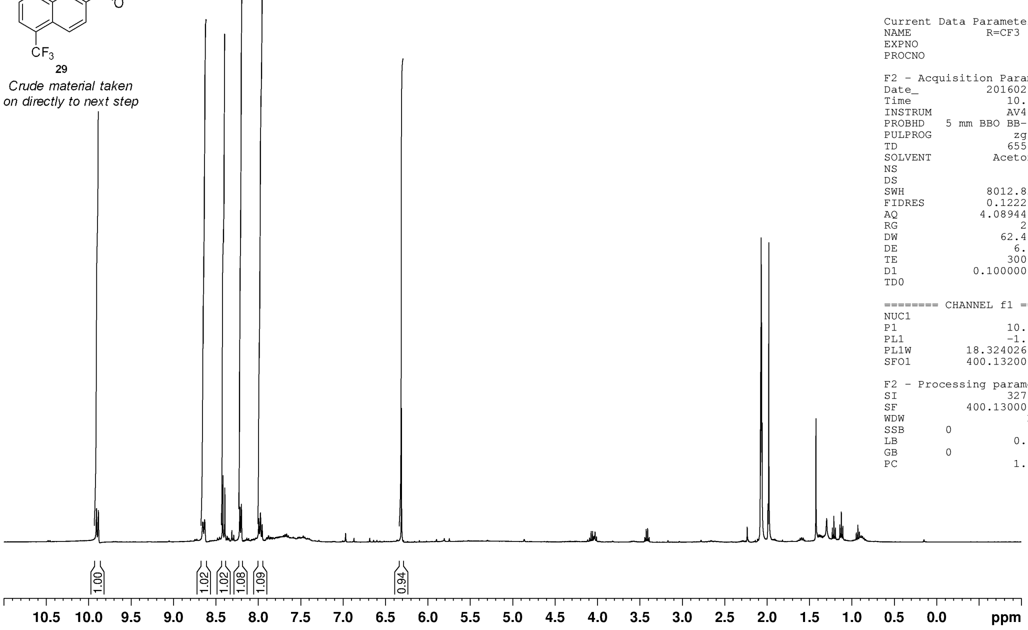


The University Of Sheffield.

```
Current Data Parameters
NAME            R=CF3 1H
EXPNO                  1
PROCNO                 1

F2 - Acquisition Parameters
Date_           20160203
Time               10.50
INSTRUM            AV400
PROBHD   5 mm BBO BB-1H
PULPROG             zg30
TD                 65536
SOLVENT          Acetone
NS                    64
DS                     4
SWH             8012.820 Hz
FIDRES          0.122266 Hz
AQ             4.0894465 sec
RG                   256
DW                62.400 usec
DE                  6.50 usec
TE                 300.0 K
D1            0.10000000 sec
TD0                    1

======== CHANNEL f1 ========
NUC1                  1H
P1                 10.00 usec
PL1                -1.90 dB
PL1W         18.32402611 W
SFO1         400.1320007 MHz

F2 - Processing parameters
SI                 32768
SF           400.1300000 MHz
WDW                   EM
SSB      0
LB                  0.30 Hz
GB       0
PC                  1.00
```

**MJF-1-84-P1**
**PRO Acetone {C:\02Feb2016} ch3sj 51**

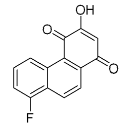


**30**
*Crude material taken on directly to next step*

The University Of Sheffield.

```
Current Data Parameters
NAME              R=F 1H
EXPNO                  9
PROCNO                 1

F2 - Acquisition Parameters
Date_           20160223
Time                0.01
INSTRUM            AV400
PROBHD   5 mm BBO BB-1H
PULPROG             zg30
TD                 65536
SOLVENT          Acetone
NS                    64
DS                     4
SWH             8012.820 Hz
FIDRES          0.122266 Hz
AQ             4.0894465 sec
RG                   362
DW                62.400 usec
DE                  6.50 usec
TE                 300.0 K
D1            0.10000000 sec
TD0                    1

======== CHANNEL f1 ========
NUC1                  1H
P1                 10.00 usec
PL1                -1.90 dB
PL1W         18.32402611 W
SFO1         400.1320007 MHz

F2 - Processing parameters
SI                 32768
SF           400.1300000 MHz
WDW                   EM
SSB      0
LB                  0.30 Hz
GB       0
PC                  1.00
```

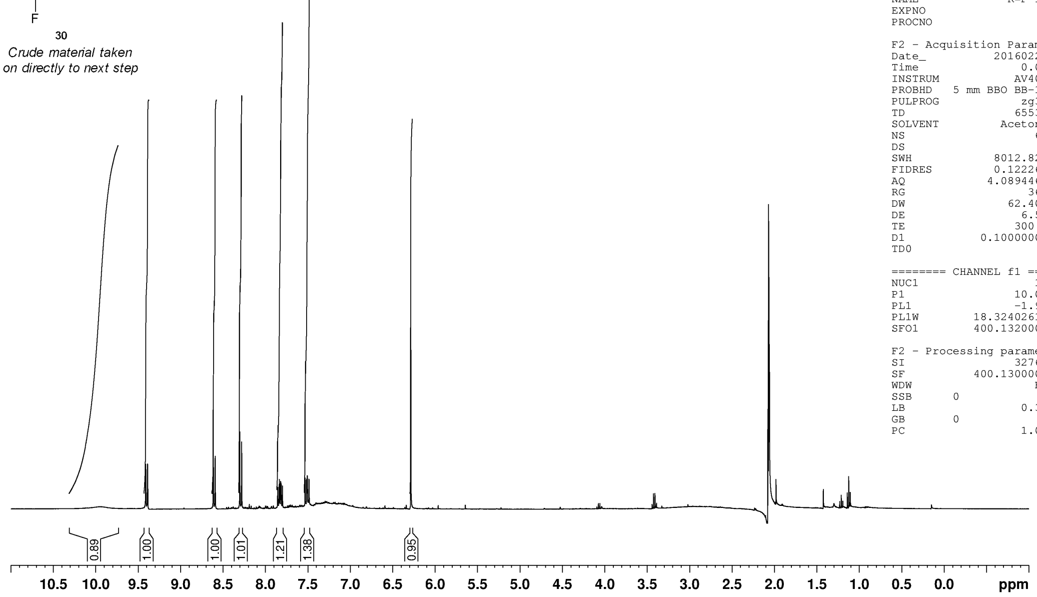


**MJF-2-41-P2**
**PRO DMSO {C:\09Sep2016} ch3sj 34**

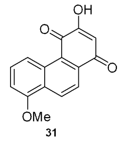


**31**

The University Of Sheffield.

```
Current Data Parameters
NAME            R=OMe 1H
EXPNO                  2
PROCNO                 1

F2 - Acquisition Parameters
Date_           20160914
Time               17.42
INSTRUM            AV400
PROBHD   5 mm BBO BB-1H
PULPROG             zg30
TD                 65536
SOLVENT             DMSO
NS                    64
DS                     4
SWH             8012.820 Hz
FIDRES          0.122266 Hz
AQ             4.0894465 sec
RG                   256
DW                62.400 usec
DE                  6.50 usec
TE                 300.0 K
D1            0.10000000 sec
TD0                    1

======== CHANNEL f1 ========
NUC1                  1H
P1                 10.00 usec
PL1                -1.90 dB
PL1W         18.32402611 W
SFO1         400.1320007 MHz

F2 - Processing parameters
SI                 32768
SF           400.1300000 MHz
WDW                   EM
SSB      0
LB                  0.30 Hz
GB       0
PC                  1.00
```

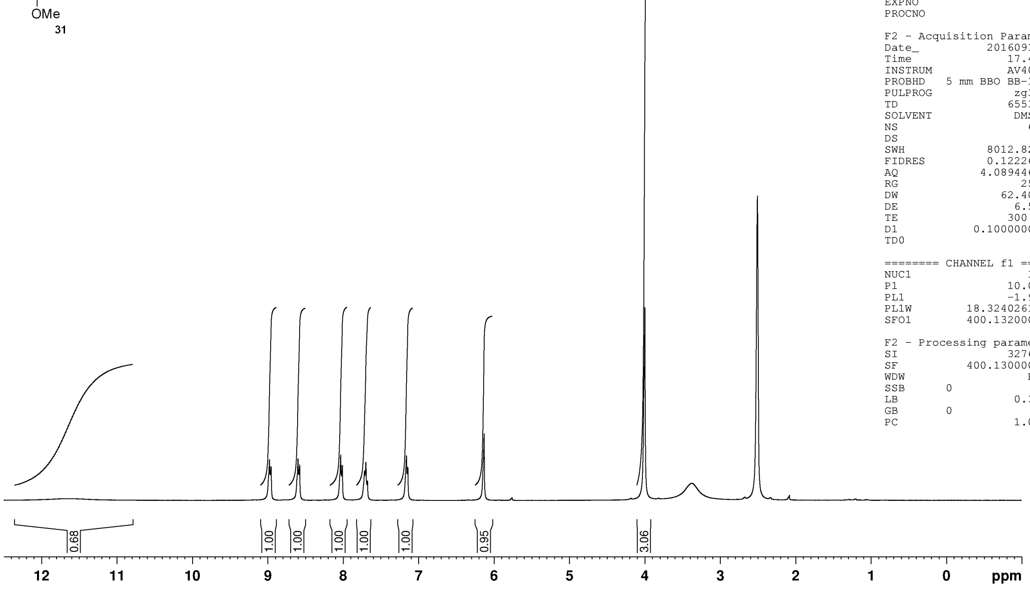

**MJF-2-43-P2A**
**PRO CDCl3 {C:\NMRData\Jones_current_year} mdp14mjf 17**

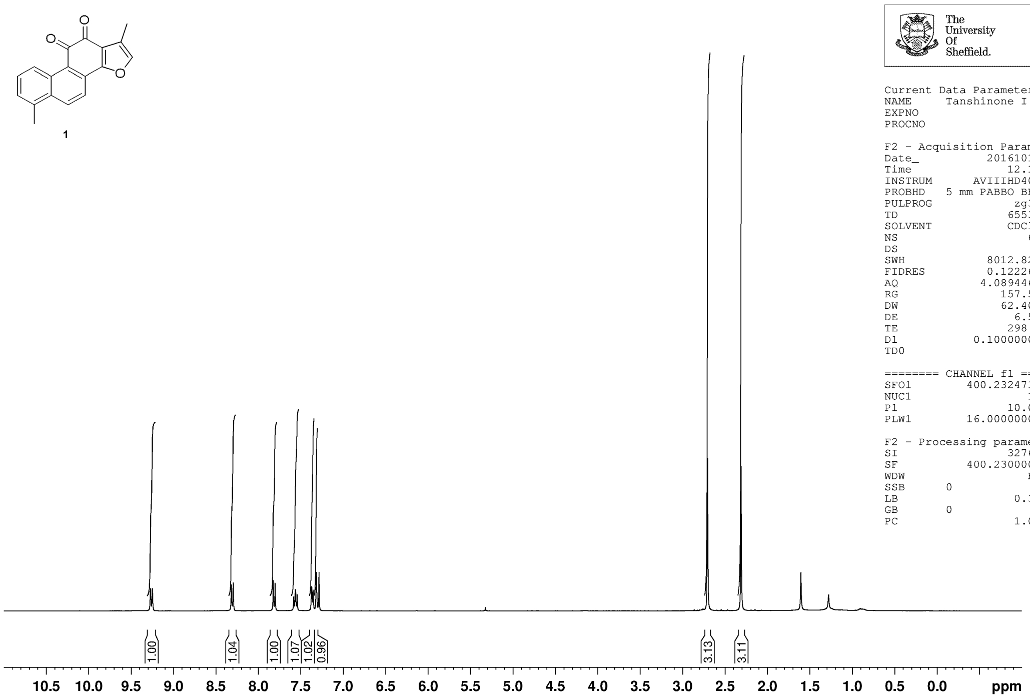


The University Of Sheffield.

```
Current Data Parameters
NAME      Tanshinone I 1H
EXPNO                   1
PROCNO                  1

F2 - Acquisition Parameters
Date_            20161018
Time                12.17
INSTRUM         AVIIIHD400
PROBHD    5 mm PABBO BB/
PULPROG              zg30
TD                  65536
SOLVENT             CDCl3
NS                     64
DS                      2
SWH              8012.820 Hz
FIDRES           0.122266 Hz
AQ              4.0894465 sec
RG                 157.59
DW                 62.400 usec
DE                   6.50 usec
TE                  298.0 K
D1             0.10000000 sec
TD0                     1

======== CHANNEL f1 ========
SFO1          400.2324716 MHz
NUC1                   1H
P1                  10.00 usec
PLW1          16.00000000 W

F2 - Processing parameters
SI                  32768
SF            400.2300000 MHz
WDW                    EM
SSB      0
LB                   0.30 Hz
GB       0
PC                   1.00
```

**MJF-1-064- F11-22**
**JMOD250ppm CDCl3 {C:\NMRData\11Nov2015} ch3sj 20**

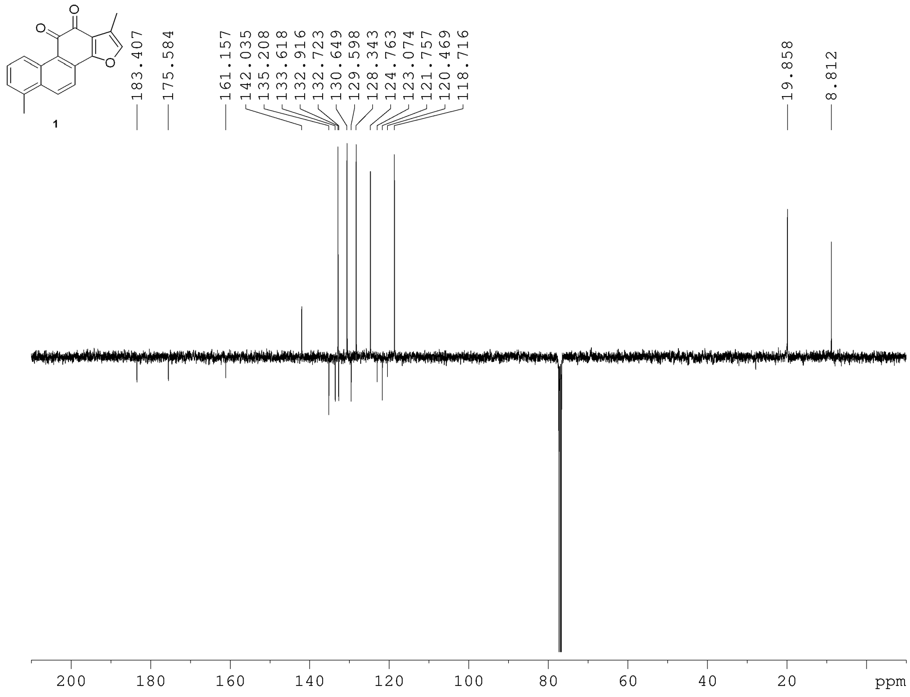


The University Of Sheffield.

```
Current Data Parameters
NAME      Tanshinone I 13C
EXPNO                   2
PROCNO                  1

F2 - Acquisition Parameters
Date_            20151108
Time                22.01
INSTRUM         AVIIIHD400
PROBHD    5 mm PABBO BB/
PULPROG              jmod
TD                  65536
SOLVENT             CDCl3
NS                    512
DS                      4
SWH             25252.525 Hz
FIDRES           0.385323 Hz
AQ              1.2976128 sec
RG                 195.75
DW                 19.800 usec
DE                   6.50 usec
TE                  298.0 K
CNST2         145.0000000
CNST11          1.0000000
D1             3.70000005 sec
D20            0.00689655 sec
TD0                     1

======== CHANNEL f1 ========
SFO1          100.6494869 MHz
NUC1                  13C
P1                  10.00 usec
P2                  20.00 usec
PLW1          71.00000000 W

======== CHANNEL f2 ========
SFO2          400.2316009 MHz
NUC2                   1H
CPDPRG[2          waltz16
PCPD2               90.00 usec
PLW2          16.00000000 W
PLW12          0.19753000 W

F2 - Processing parameters
SI                  32768
SF            100.6379135 MHz
WDW                    EM
SSB      0
LB                   2.00 Hz
GB       0
PC                   1.40
```

**MJF-2-43-P1D**
**PRO CDCl3 {C:\NMRData\Jones_current_year} mdp14mjf 55**

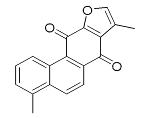

**32**

The University Of Sheffield.

```
Current Data Parameters
NAME      Isotanshinone I 1H
EXPNO                  2
PROCNO                 1

F2 - Acquisition Parameters
Date_           20161018
Time               18.00
INSTRUM      AVIIIHD400
PROBHD   5 mm PABBO BB/
PULPROG             zg30
TD                 65536
SOLVENT            CDCl3
NS                    64
DS                     2
SWH             8012.820 Hz
FIDRES          0.122266 Hz
AQ             4.0894465 sec
RG                128.39
DW                62.400 usec
DE                  6.50 usec
TE                 298.0 K
D1            0.10000000 sec
TD0                    1

======== CHANNEL f1 ========
SFO1         400.2324716 MHz
NUC1                  1H
P1                 10.00 usec
PLW1         16.00000000 W

F2 - Processing parameters
SI                 32768
SF           400.2300000 MHz
WDW                   EM
SSB      0
LB                  0.30 Hz
GB       0
PC                  1.00
```

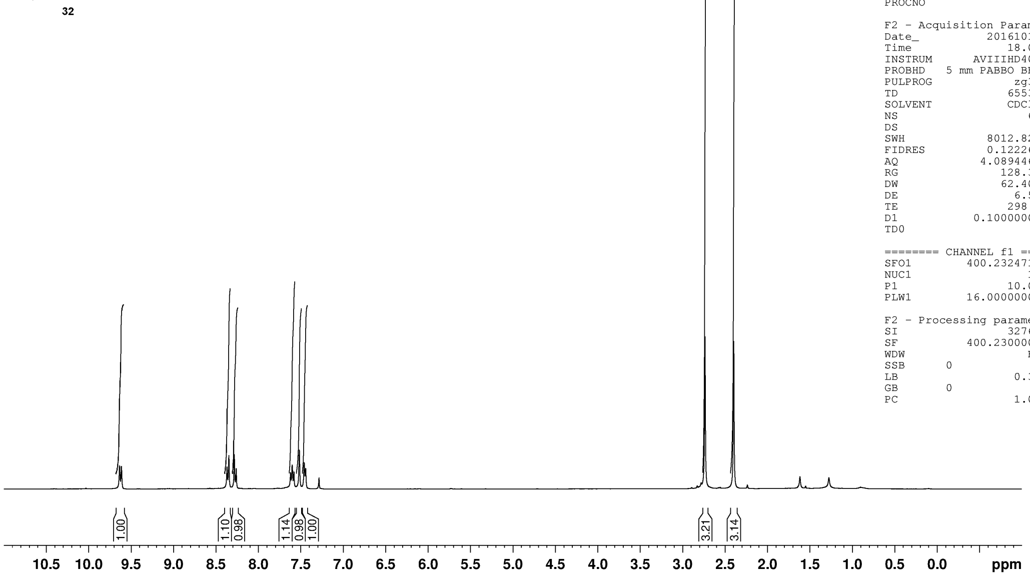


**MJF-2-43-P1D**
**JMOD250ppm CDCl3 {C:\NMRData\Jones_current_year} mdp14mjf 29**

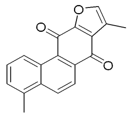

**32**

The University Of Sheffield.

```
Current Data Parameters
NAME             R=Me 13C
EXPNO                   3
PROCNO                  1

F2 - Acquisition Parameters
Date_            20161019
Time                 0.53
INSTRUM        AVIIIHD400
PROBHD     5 mm PABBO BB/
PULPROG              jmod
TD                  65536
SOLVENT             CDCl3
NS                    512
DS                      4
SWH             25252.525 Hz
FIDRES           0.385323 Hz
AQ              1.2976128 sec
RG                 195.75
DW                 19.800 usec
DE                   6.50 usec
TE                  298.0 K
CNST2         145.0000000
CNST11          1.0000000
D1             3.70000005 sec
D20            0.00689655 sec
TD0                     1

======== CHANNEL f1 ========
SFO1          100.6494869 MHz
NUC1                  13C
P1                  10.00 usec
P2                  20.00 usec
PLW1          71.00000000 W

======== CHANNEL f2 ========
SFO2          400.2316009 MHz
NUC2                   1H
CPDPRG[2          waltz16
PCPD2               90.00 usec
PLW2          16.00000000 W
PLW12          0.19753000 W

F2 - Processing parameters
SI                  32768
SF            100.6379135 MHz
WDW                    EM
SSB       0
LB                   2.00 Hz
GB        0
PC                   1.40
```

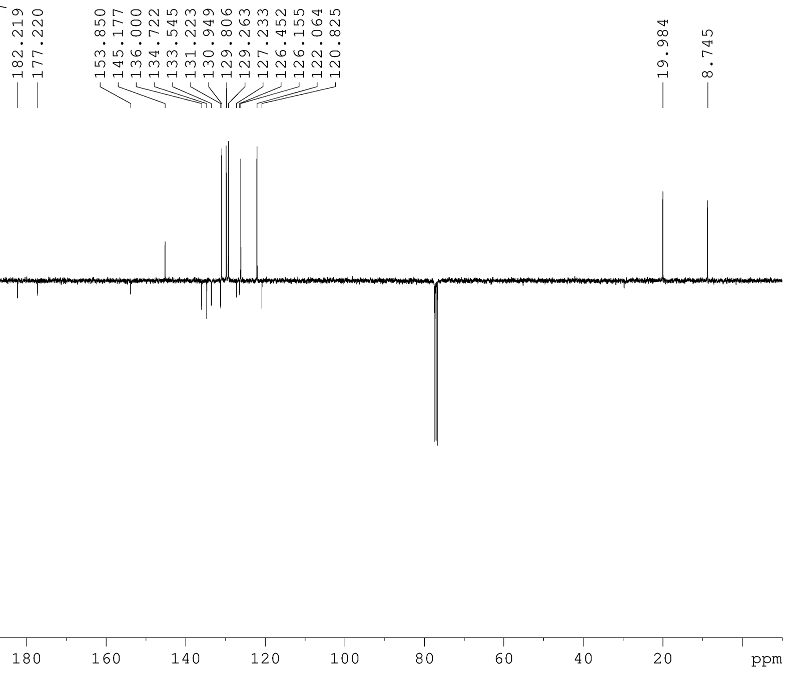

**MJF−2−45−P2C**
**PRO CDCl3 {C:\NMRData\Jones_current_year} mdp14mjf 2**

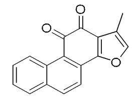

**33**

The University Of Sheffield.

```
Current Data Parameters
NAME                R=H 1H
EXPNO                    4
PROCNO                   1

F2 - Acquisition Parameters
Date_             20161128
Time                 21.19
INSTRUM        AVIIIHD400
PROBHD   5 mm PABBO BB/
PULPROG               zg30
TD                   65536
SOLVENT              CDCl3
NS                      64
DS                       2
SWH             8012.820 Hz
FIDRES          0.122266 Hz
AQ             4.0894465 sec
RG                   175.8
DW                62.400 usec
DE                  6.50 usec
TE                 293.9 K
D1            0.10000000 sec
TD0                      1

======== CHANNEL f1 ========
SFO1         400.2324716 MHz
NUC1                    1H
P1                   10.00 usec
PLW1         16.00000000 W

F2 - Processing parameters
SI                   32768
SF           400.2300000 MHz
WDW                     EM
SSB      0
LB                    0.30 Hz
GB       0
PC                    1.00
```

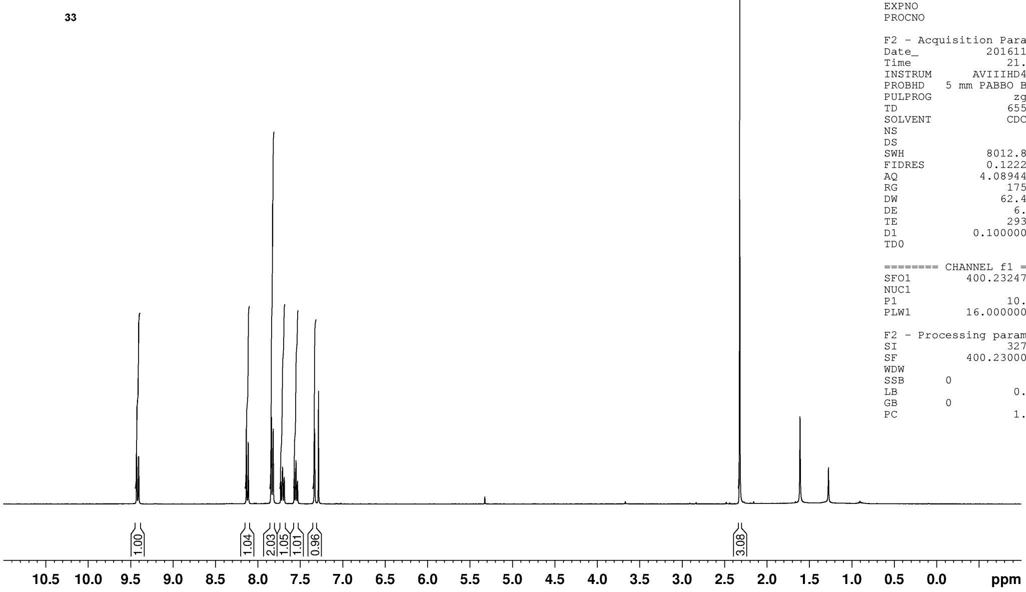

**MJF-2-69-P1A**
**PROTON.s CDCl3 {C:\NMRData\Jones_current_year} mdp14mjf 34**

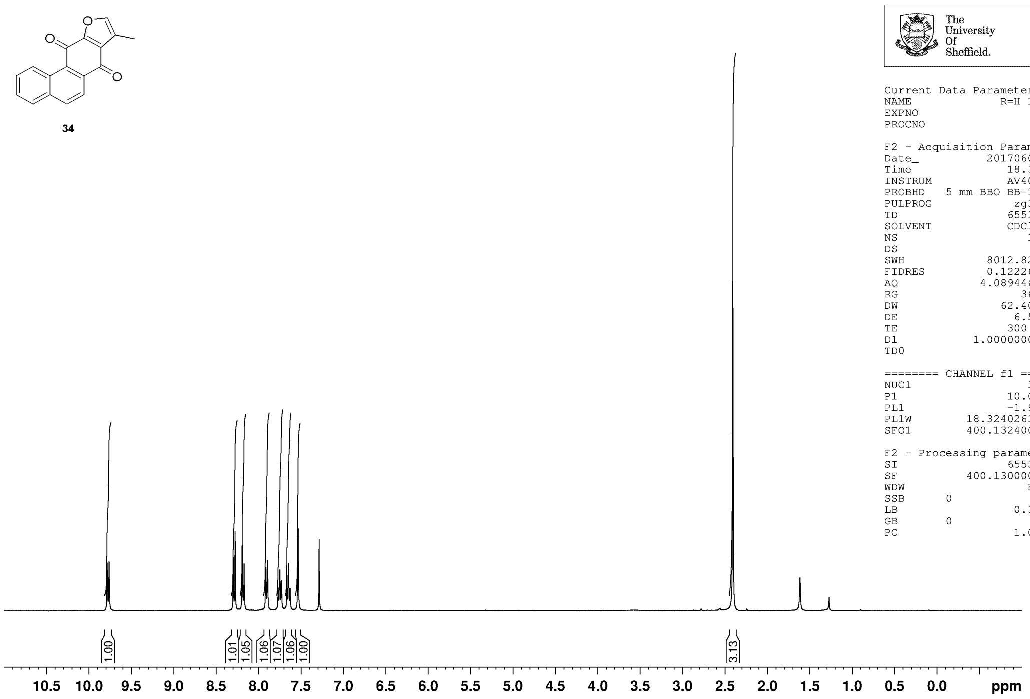


The University Of Sheffield.

```
Current Data Parameters
NAME              R=H 1H
EXPNO                  2
PROCNO                 1

F2 - Acquisition Parameters
Date_           20170606
Time               18.36
INSTRUM            AV400
PROBHD   5 mm BBO BB-1H
PULPROG             zg30
TD                 65536
SOLVENT            CDCl3
NS                    16
DS                     0
SWH             8012.820 Hz
FIDRES          0.122266 Hz
AQ             4.0894465 sec
RG                   362
DW                62.400 usec
DE                  6.50 usec
TE                 300.0 K
D1            1.00000000 sec
TD0                    1

======== CHANNEL f1 ========
NUC1                  1H
P1                 10.00 usec
PL1                -1.90 dB
PL1W         18.32402611 W
SFO1         400.1324008 MHz

F2 - Processing parameters
SI                 65536
SF           400.1300000 MHz
WDW                   EM
SSB      0
LB                  0.30 Hz
GB       0
PC                  1.00
```

**MJF-2-45-P1A**
**JMOD250ppm CDCl3 {C:\NMRData\Jones_current_year} mdp14mjf 57**

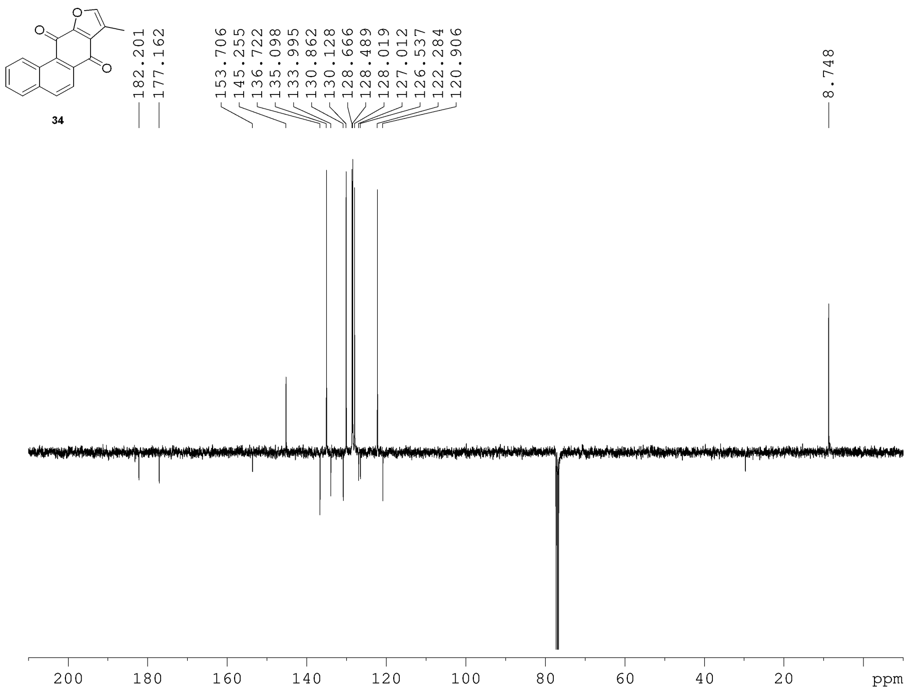


The University Of Sheffield.

```
Current Data Parameters
NAME              R=H 13C
EXPNO                  2
PROCNO                 1

F2 - Acquisition Parameters
Date_           20170123
Time               21.33
INSTRUM        AVIIIHD400
PROBHD   5 mm PABBO BB/
PULPROG             jmod
TD                 65536
SOLVENT            CDCl3
NS                   512
DS                     4
SWH            25252.525 Hz
FIDRES          0.385323 Hz
AQ             1.2976128 sec
RG                195.75
DW                19.800 usec
DE                  6.50 usec
TE                 298.0 K
CNST2        145.0000000
CNST11         1.0000000
D1            3.70000005 sec
D20           0.00689655 sec
TD0                    1

======== CHANNEL f1 ========
SFO1         100.6494869 MHz
NUC1                 13C
P1                 10.00 usec
P2                 20.00 usec
PLW1         71.00000000 W

======== CHANNEL f2 ========
SFO2         400.2316009 MHz
NUC2                  1H
CPDPRG[2         waltz16
PCPD2              90.00 usec
PLW2         16.00000000 W
PLW12         0.19753000 W

F2 - Processing parameters
SI                 32768
SF           100.6379135 MHz
WDW                   EM
SSB      0
LB                  2.00 Hz
GB       0
PC                  1.40
```

**MJF-2-49-F1**
**PROTON.s CDCl3 {C:\NMRData\Jones_current_year} mdp14mjf 39**

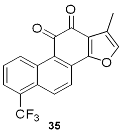

35

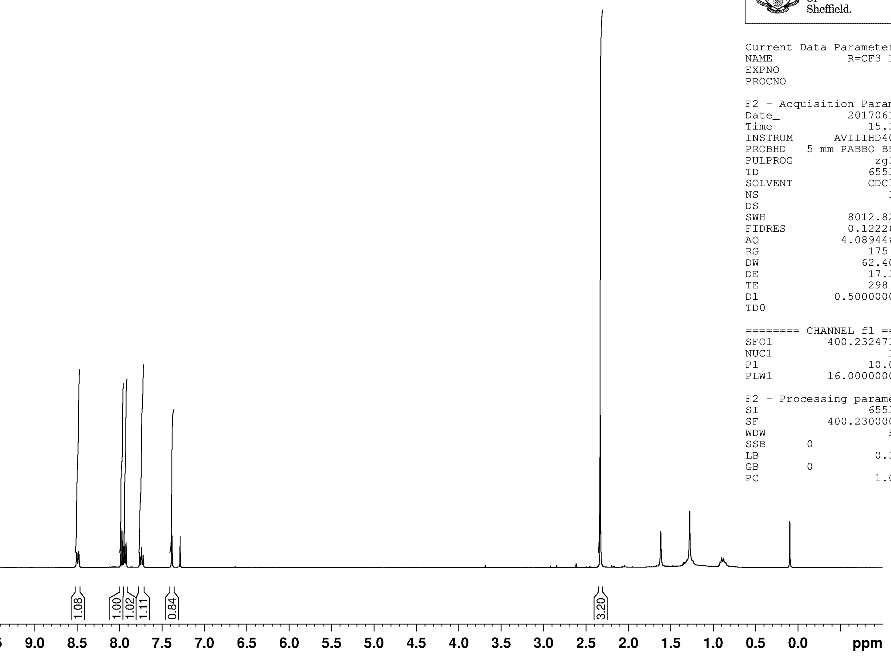


The University Of Sheffield.

```
Current Data Parameters
NAME             R=CF3 1H
EXPNO                   1
PROCNO                  1

F2 - Acquisition Parameters
Date_            20170613
Time                15.33
INSTRUM        AVIIIHD400
PROBHD   5 mm PABBO BB/
PULPROG              zg30
TD                  65536
SOLVENT             CDCl3
NS                     16
DS                      2
SWH              8012.820 Hz
FIDRES           0.122266 Hz
AQ              4.0894465 sec
RG                  175.8
DW                 62.400 usec
DE                  17.37 usec
TE                  298.0 K
D1             0.50000000 sec
TD0                     1

======== CHANNEL f1 ========
SFO1          400.2324716 MHz
NUC1                   1H
P1                  10.00 usec
PLW1          16.00000000 W

F2 - Processing parameters
SI                  65536
SF            400.2300000 MHz
WDW                    EM
SSB      0
LB                   0.30 Hz
GB       0
PC                   1.00
```

**Matt Foulks, mdp14mjf, Jones**
**MJF-2-49-F1A**
**C13udeft.lub CDCl3 {C:\NMRData\Jones_current_year\data\mdp14mjf\nmr} nmr 27**

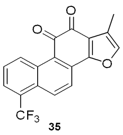

35

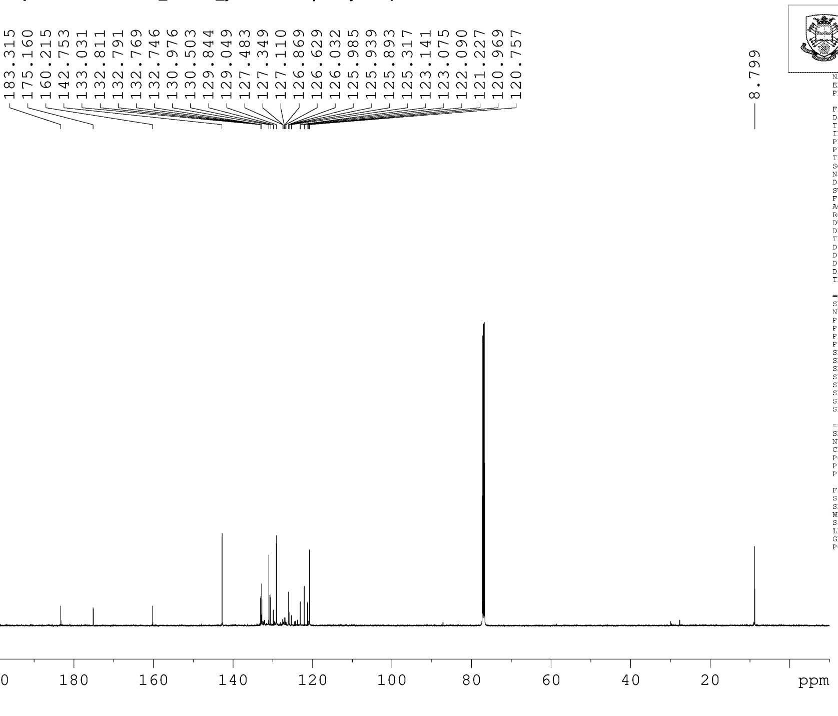


The University Of Sheffield.

```
NAME             R=CF3 13C
EXPNO                   1
PROCNO                  1

F2 - Acquisition Parameters
Date_            20170616
Time                20.35
INSTRUM             spect
PROBHD   5 mm PABBO BB/
PULPROG             udeft
TD                  21424
SOLVENT             CDCl3
NS                  20480
DS                      0
SWH             29761.904 Hz
FIDRES           1.389185 Hz
AQ              0.3599232 sec
RG                  189.1
DW                 16.800 use
DE                   8.90 use
TE                  298.0 K
D1             4.00000000 sec
D11            0.03000000 sec
D12            0.00002000 sec
D20          200.00000000 sec
TD0                    20

======== CHANNEL f1 =======
SFO1          125.7703656 MHz
NUC1                  13C
P1                   9.00 use
P13               2000.00 use
P26                500.00 use
PLW1          95.00000000 W
SPNAM[5]     Crp60comp.4
SPOAL5              0.500
SPOFFS5  0 Hz
SPW5          11.75699997 W
SPNAM[8] Crp60,0.5,20.1
SPOAL8              0.500
SPOFFS8  0 Hz
SPW8          11.75699997 W

======== CHANNEL f2 =======
SFO2          500.1320007 MHz
NUC2                   1H
CPDPRG[2         waltz16
PCPD2               80.00 use
PLW2          22.00000000 W
PLW12          0.34375000 W

F2 - Processing parameters
SI                 131072
SF            125.7577885 MHz
WDW                    EM
SSB      0
LB                   2.00 Hz
GB       0
PC                   1.40
```

**MJF−2−49−F1A**
**19F_DEC.s CDCl3 {C:\NMRData\Jones_current_year} mdp14mjf 23**

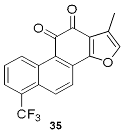


**35**

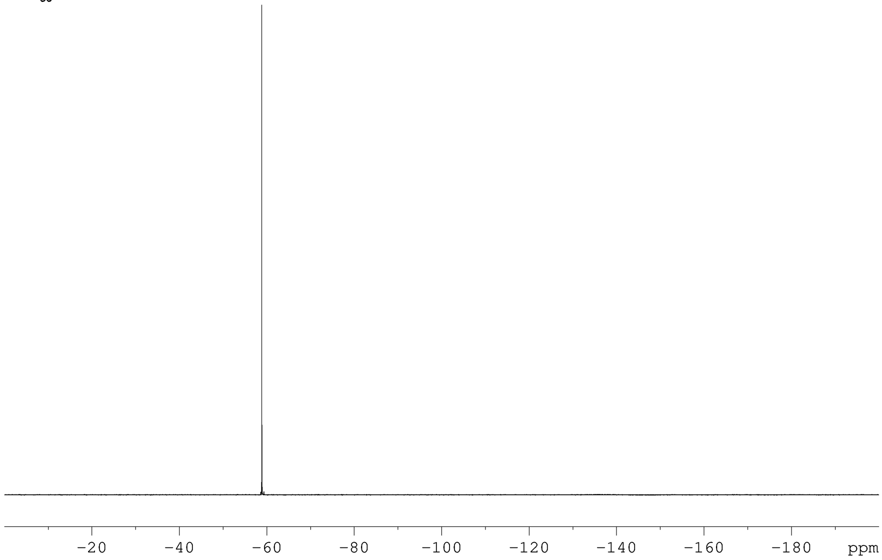


The University Of Sheffield.

```
Current Data Parameters
NAME             R=CF3 19F
EXPNO                    2
PROCNO                   1

F2 - Acquisition Parameters
Date_             20170620
Time                 13.24
INSTRUM          AVIIIHD400
PROBHD   5 mm PABBO BB/
PULPROG               zgig
TD                  167936
SOLVENT              CDCl3
NS                      16
DS                       4
SWH             89285.711 Hz
FIDRES           0.531665 Hz
AQ              0.9404416 sec
RG                  195.75
DW                   5.600 usec
DE                  150.63 usec
TE                   298.0 K
D1              1.00000000 sec
D11             0.03000000 sec
TD0                      1

======== CHANNEL f1 ========
SFO1          376.5548008 MHz
NUC1                   19F
P1                   13.50 usec
PLW1           25.00000000 W

======== CHANNEL f2 ========
SFO2          400.2324716 MHz
NUC2                    1H
CPDPRG[2           waltz16
PCPD2                90.00 usec
PLW2           16.00000000 W
PLW12           0.19753000 W

F2 - Processing parameters
SI                  131072
SF            376.5924600 MHz
WDW                     EM
SSB      0
LB                    1.00 Hz
GB       0
PC                    1.00
```

**MJF−2−49−P1A**
**PRO CDCl3 {C:\NMRData\Jones_current_year} mdp14mjf 34**

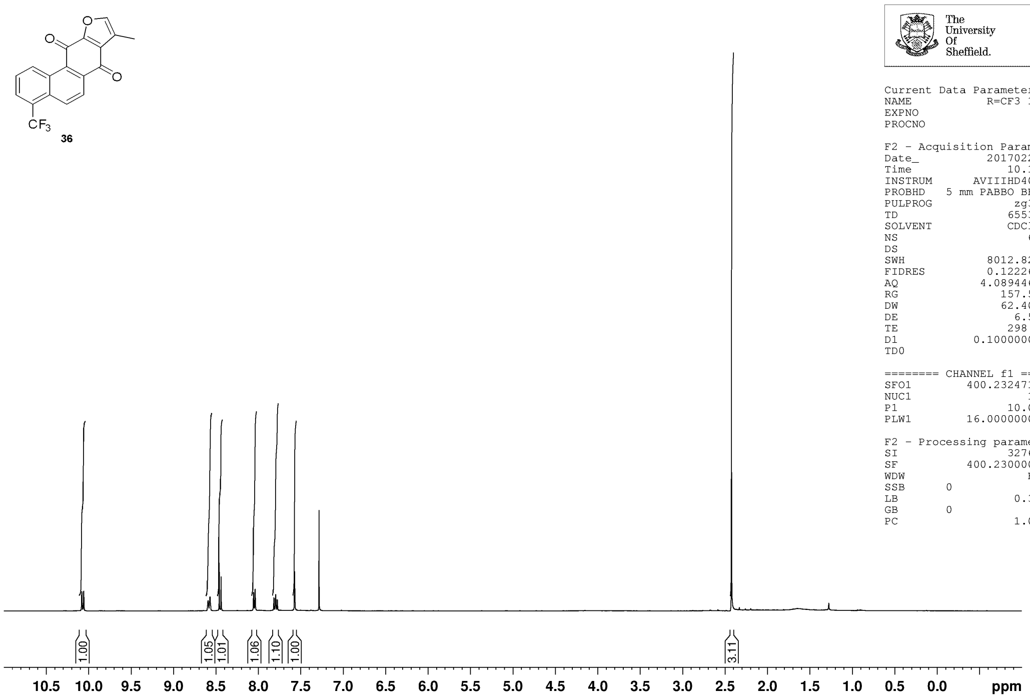


The University Of Sheffield.

```
Current Data Parameters
NAME              R=CF3 1H
EXPNO                    4
PROCNO                   1

F2 - Acquisition Parameters
Date_             20170225
Time                 10.19
INSTRUM         AVIIIHD400
PROBHD   5 mm PABBO BB/
PULPROG               zg30
TD                   65536
SOLVENT              CDCl3
NS                      64
DS                       2
SWH               8012.820 Hz
FIDRES            0.122266 Hz
AQ               4.0894465 sec
RG                  157.59
DW                  62.400 usec
DE                    6.50 usec
TE                   298.0 K
D1              0.10000000 sec
TD0                      1

======== CHANNEL f1 ========
SFO1           400.2324716 MHz
NUC1                    1H
P1                   10.00 usec
PLW1           16.00000000 W

F2 - Processing parameters
SI                   32768
SF             400.2300000 MHz
WDW                     EM
SSB      0
LB                    0.30 Hz
GB       0
PC                    1.00
```

**MJF−2−50−P1A**
**JMOD250ppm CDCl3 {C:\NMRData\Jones_current_year} mdp14mjf 7**

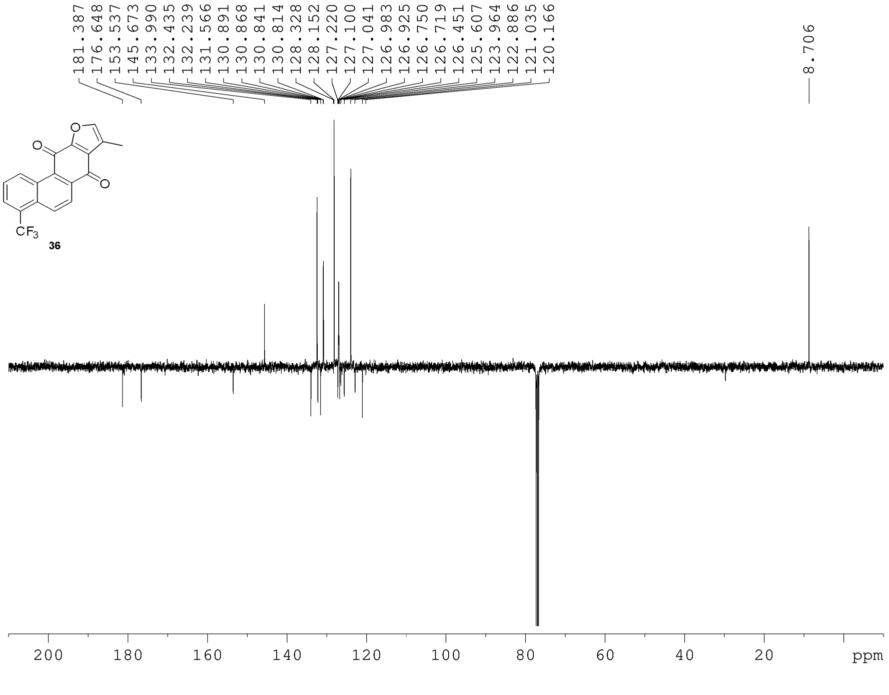


The University Of Sheffield.

```
NAME              R=CF3 13C
EXPNO                     3
PROCNO                    1

F2 - Acquisition Parameters
Date_              20170226
Time                  16.53
INSTRUM          AVIIIHD400
PROBHD   5 mm PABBO BB/
PULPROG                jmod
TD                    65536
SOLVENT               CDCl3
NS                      512
DS                        4
SWH               25252.525 Hz
FIDRES             0.385323 Hz
AQ                1.2976128 sec
RG                   195.75
DW                   19.800 usec
DE                     6.50 usec
TE                    298.0 K
CNST2           145.0000000
CNST11            1.0000000
D1               3.70000005 sec
D20              0.00689655 sec
TD0                       1

======== CHANNEL f1 ========
SFO1            100.6494869 MHz
NUC1                    13C
P1                    10.00 usec
P2                    20.00 usec
PLW1            71.00000000 W

======== CHANNEL f2 ========
SFO2            400.2316009 MHz
NUC2                     1H
CPDPRG[2            waltz16
PCPD2                 90.00 usec
PLW2            16.00000000 W
PLW12            0.19753000 W

F2 - Processing parameters
SI                    32768
SF              100.6379135 MHz
WDW                      EM
SSB      0
LB                     2.00 Hz
GB       0
PC                     1.40
```

**MJF−2−49−P1A**
**F19CPDA3 CDCl3 {C:\NMRData\Jones_current_year} mdp14mjf 3**

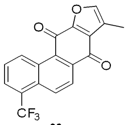

36

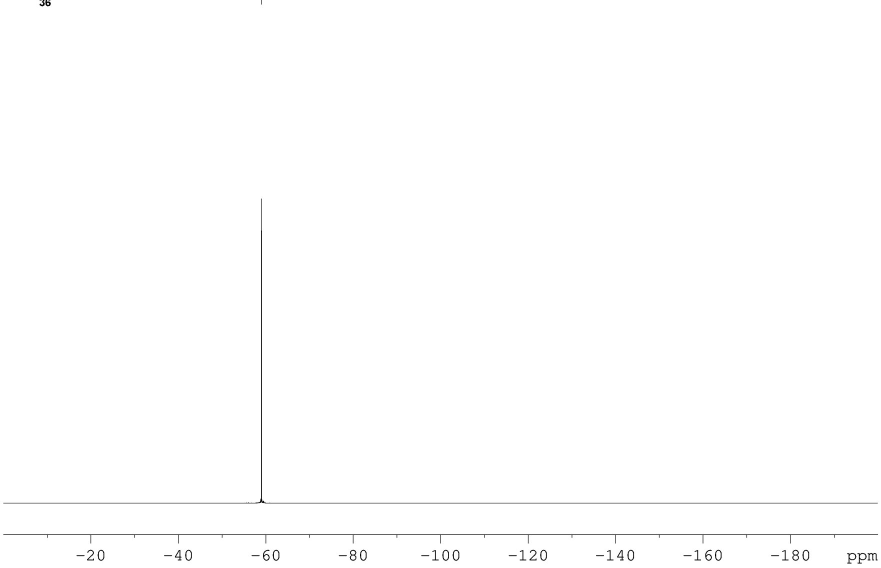


The University Of Sheffield.

```
Current Data Parameters
NAME             R=CF3 19F
EXPNO                    1
PROCNO                   1

F2 - Acquisition Parameters
Date_             20170226
Time                 13.46
INSTRUM          AVIIIHD400
PROBHD   5 mm PABBO BB/
PULPROG         zgfhigqn.2
TD                  130936
SOLVENT              CDCl3
NS                     128
DS                       4
SWH             113636.367 Hz
FIDRES            0.867877 Hz
AQ               0.5761184 sec
RG                  195.75
DW                   4.400 usec
DE                    6.50 usec
TE                   298.0 K
D1              1.50000000 sec
D11             0.03000000 sec
D12             0.00002000 sec
TD0                      1

======== CHANNEL f1 ========
SFO1           376.5397373 MHz
NUC1                   19F
P1                   13.50 usec
PLW1           25.00000000 W

======== CHANNEL f2 ========
SFO2           400.2316009 MHz
NUC2                    1H
CPDPRG[2           waltz16
PCPD2                90.00 usec
PLW2           16.00000000 W
PLW12           0.19753000 W

F2 - Processing parameters
SI                   65536
SF             376.5924602 MHz
WDW                     EM
SSB      0
LB                    2.00 Hz
GB       0
PC                    1.00
```

**MJF-2-50-P2A**
**PROTON.s CDCl3 {C:\NMRData\Jones_current_year} mdp14mjf 36**

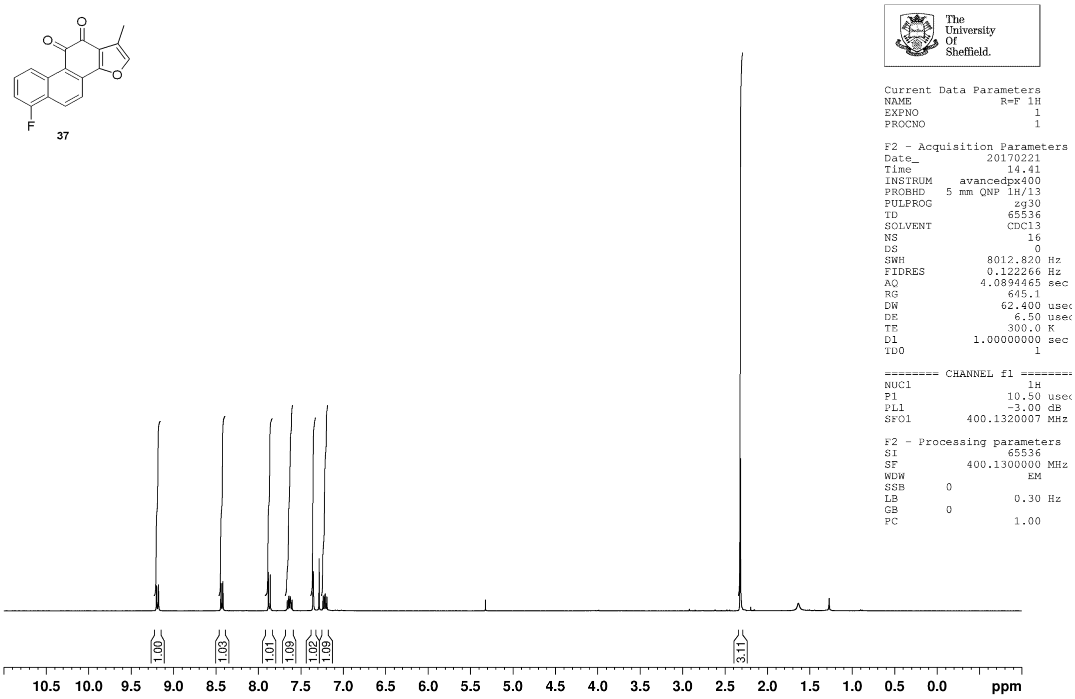


**MJF-2-50-P2A**
**JMOD250ppm CDCl3 {C:\NMRData\Jones_current_year} mdp14mjf 48**

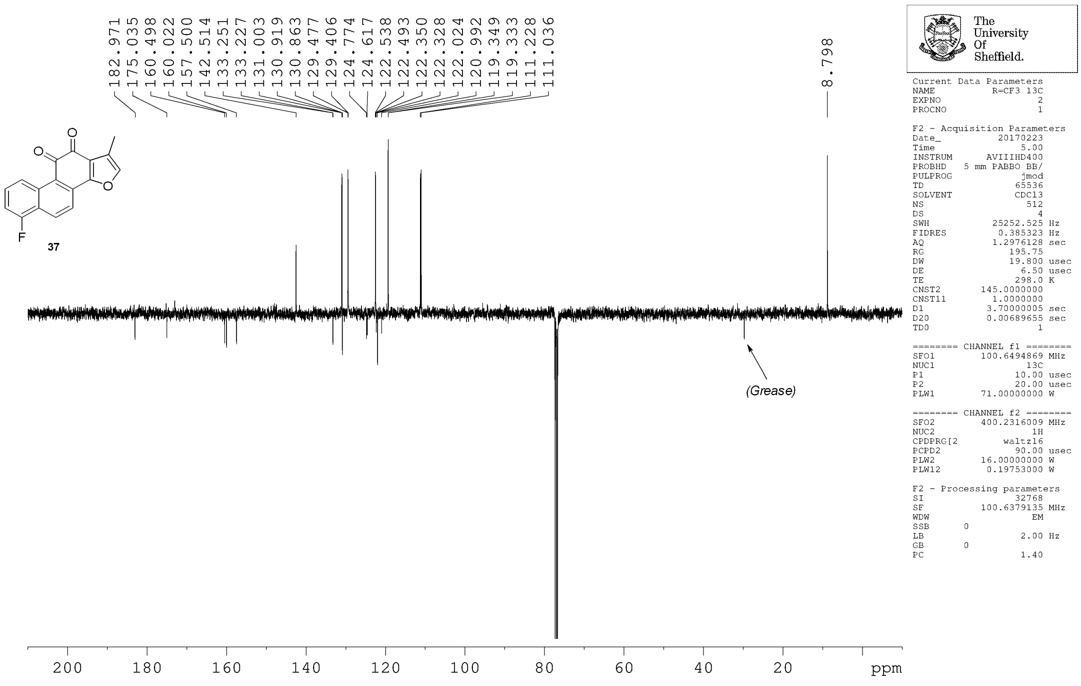

**MJF–2–50–P2A**
**F19CPD.s CDCl3 {C:\NMRData\Jones_current_year} mdp14mjf 8**

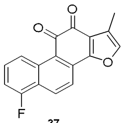


**37**

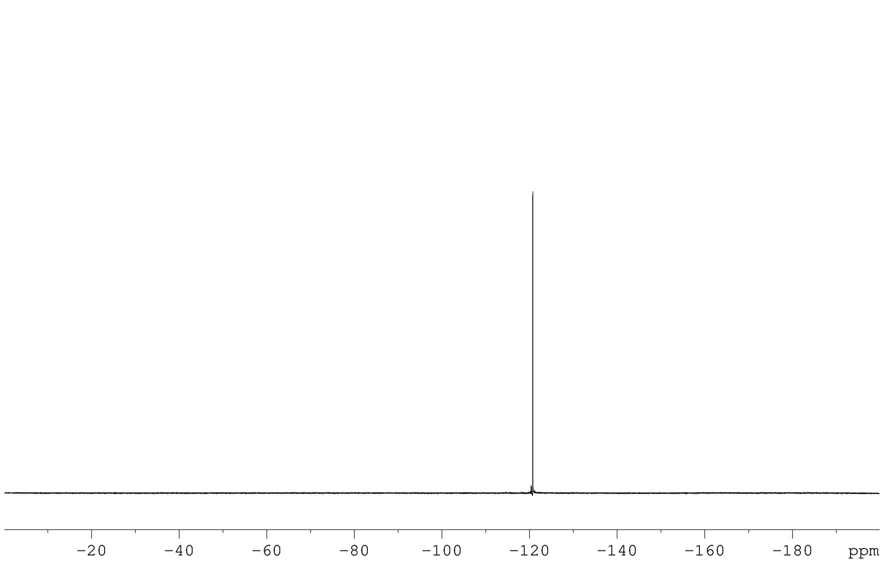


The University Of Sheffield.

Current Data Parameters
NAME R=F 19F
EXPNO 2
PROCNO 1

F2 - Acquisition Parameters
Date_ 20170224
Time 11.44
INSTRUM avancedpx400
PROBHD 5 mm QNP 1H/13
PULPROG zgfhigqn.2
TD 131072
SOLVENT CDCl3
NS 32
DS 4
SWH 75187.969 Hz
FIDRES 0.573639 Hz
AQ 0.8716288 sec
RG 3251
DW 6.650 usec
DE 6.50 usec
TE 300.0 K
D1 1.00000000 sec
D11 0.03000000 sec
D12 0.00002000 sec
TD0 1

======== CHANNEL f1 ========
NUC1 19F
P1 7.80 usec
PL1 3.00 dB
SFO1 376.4607162 MHz

======== CHANNEL f2 ========
CPDPRG[2 waltz16
NUC2 1H
PCPD2 100.00 usec
PL2 -3.00 dB
PL12 16.58 dB
SFO2 400.1316005 MHz

F2 - Processing parameters
SI 131072
SF 376.4983660 MHz
WDW EM
SSB 0
LB 2.00 Hz
GB 0
PC 1.00

**MJF-2-50-P1A**
**PRO CDCl3 {C:\NMRData\Jones_current_year} mdp14mjf 27**

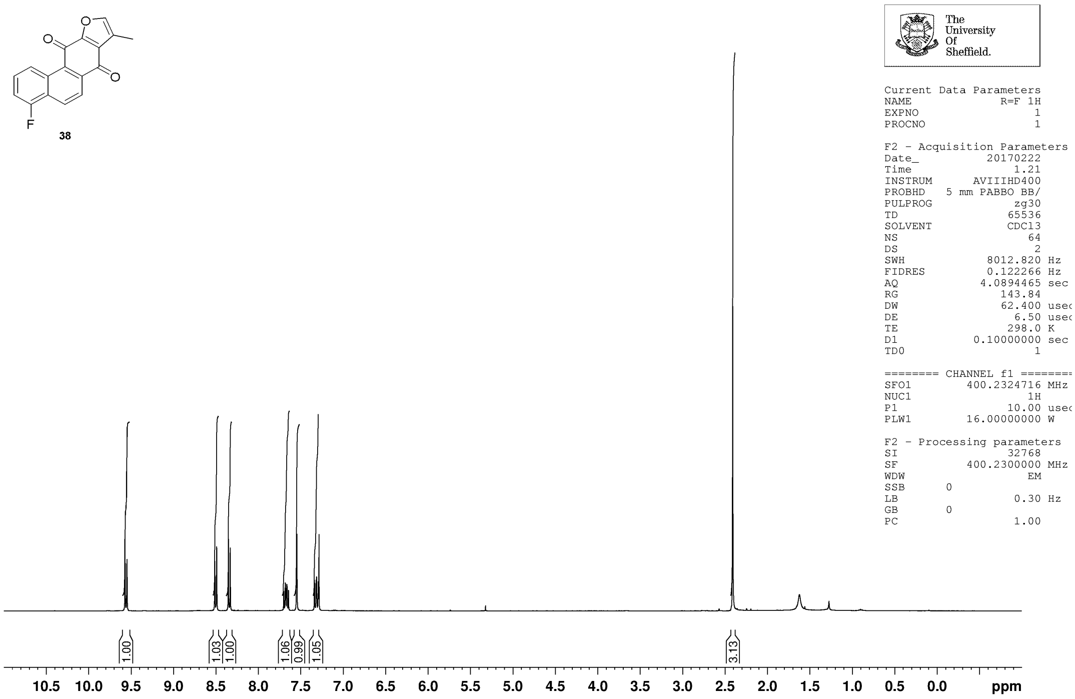


**MJF-2-50-P1A**
**JMOD250ppm CDCl3 {C:\NMRData\Jones_current_year} mdp14mjf 27**

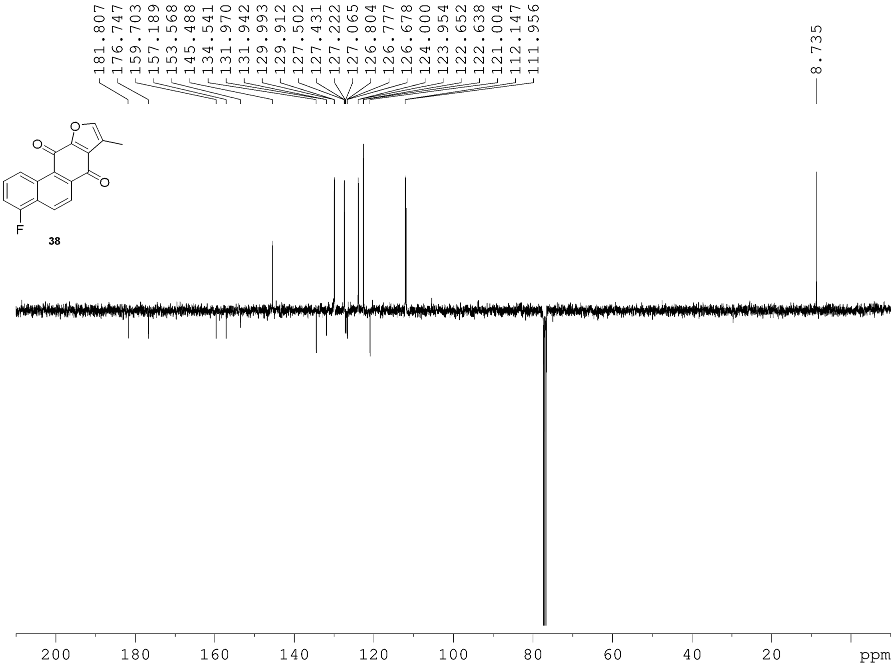


The University Of Sheffield.

```
Current Data Parameters
NAME            R=F 13C
EXPNO                 2
PROCNO                1

F2 - Acquisition Parameters
Date_          20170222
Time               2.06
INSTRUM      AVIIIHD400
PROBHD   5 mm PABBO BB/
PULPROG            jmod
TD                65536
SOLVENT           CDCl3
NS                  512
DS                    4
SWH           25252.525 Hz
FIDRES         0.385323 Hz
AQ            1.2976128 sec
RG               195.75
DW               19.800 usec
DE                 6.50 usec
TE                298.0 K
CNST2       145.0000000
CNST11        1.0000000
D1           3.70000005 sec
D20          0.00689655 sec
TD0                   1

======== CHANNEL f1 ========
SFO1        100.6494869 MHz
NUC1                13C
P1                10.00 usec
P2                20.00 usec
PLW1        71.00000000 W

======== CHANNEL f2 ========
SFO2        400.2316009 MHz
NUC2                 1H
CPDPRG[2        waltz16
PCPD2             90.00 usec
PLW2        16.00000000 W
PLW12        0.19753000 W

F2 - Processing parameters
SI                32768
SF          100.6379135 MHz
WDW                  EM
SSB      0
LB                 2.00 Hz
GB       0
PC                 1.40
```

**MJF–2–50–P1A**
**F19CPDA3 CDCl3 {C:\NMRData\Jones_current_year} mdp14mjf 41**

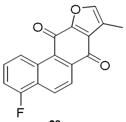


38

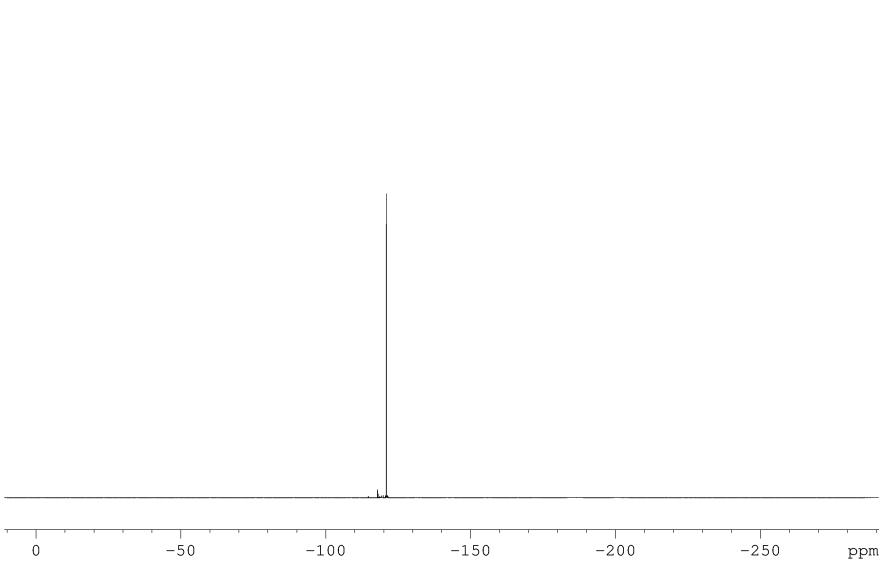


The University Of Sheffield.

```
Current Data Parameters
NAME              R=F 19F
EXPNO                   1
PROCNO                  1

F2 - Acquisition Parameters
Date_            20170222
Time                15.11
INSTRUM         AVIIIHD400
PROBHD   5 mm PABBO BB/
PULPROG       zgfhigqn.2
TD                 130936
SOLVENT             CDCl3
NS                    128
DS                      4
SWH            113636.367 Hz
FIDRES           0.867877 Hz
AQ              0.5761184 sec
RG                 195.75
DW                  4.400 usec
DE                   6.50 usec
TE                  298.0 K
D1             1.50000000 sec
D11            0.03000000 sec
D12            0.00002000 sec
TD0                     1

======== CHANNEL f1 ========
SFO1          376.5397373 MHz
NUC1                  19F
P1                  13.50 usec
PLW1          25.00000000 W

======== CHANNEL f2 ========
SFO2          400.2316009 MHz
NUC2                   1H
CPDPRG[2          waltz16
PCPD2               90.00 usec
PLW2          16.00000000 W
PLW12          0.19753000 W

F2 - Processing parameters
SI                  65536
SF            376.5924602 MHz
WDW                    EM
SSB      0
LB                   2.00 Hz
GB       0
PC                   1.00
```

**MJF-2-44-P2D**
**PRO CDCl3 {C:\NMRData\Jones_current_year} mdp14mjf 38**

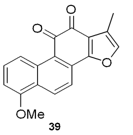

39

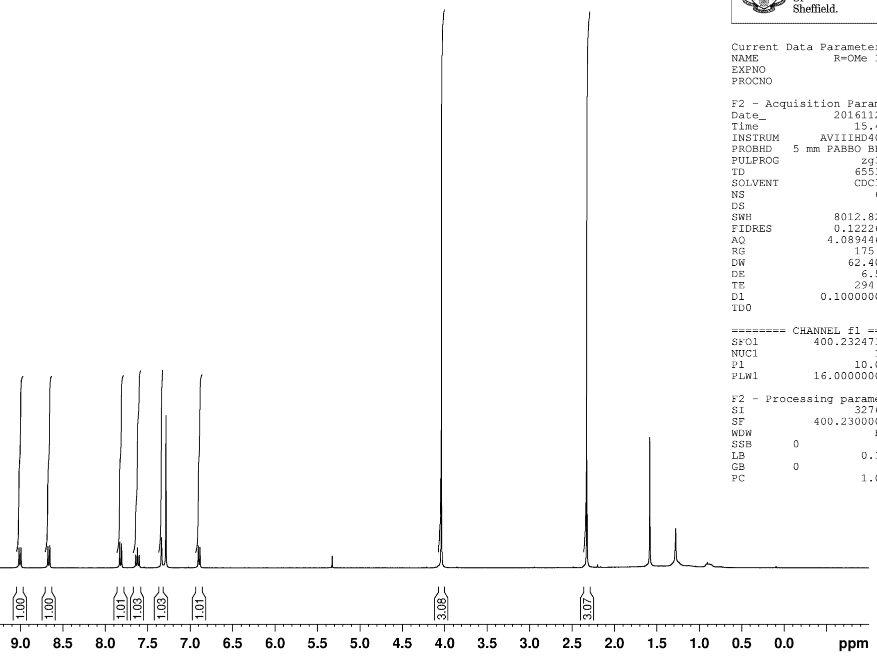


The University Of Sheffield.

```
Current Data Parameters
NAME             R=OMe 1H
EXPNO                   7
PROCNO                  1

F2 - Acquisition Parameters
Date_            20161125
Time                15.48
INSTRUM         AVIIIHD400
PROBHD   5 mm PABBO BB/
PULPROG              zg30
TD                  65536
SOLVENT             CDCl3
NS                     64
DS                      2
SWH              8012.820 Hz
FIDRES           0.122266 Hz
AQ              4.0894465 sec
RG                  175.8
DW                 62.400 usec
DE                   6.50 usec
TE                  294.5 K
D1             0.10000000 sec
TD0                     1

======== CHANNEL f1 ========
SFO1          400.2324716 MHz
NUC1                   1H
P1                  10.00 usec
PLW1          16.00000000 W

F2 - Processing parameters
SI                  32768
SF            400.2300000 MHz
WDW                    EM
SSB      0
LB                   0.30 Hz
GB       0
PC                   1.00
```

**Matt Foulks, mdp14mjf, Jones**
**MJF-2-68-P2D**
**C13udeft.lub CDCl3 {C:\NMRData\Jones_current_year\data\mdp14mjf\nmr} nmr 28**

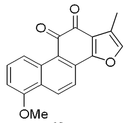

39

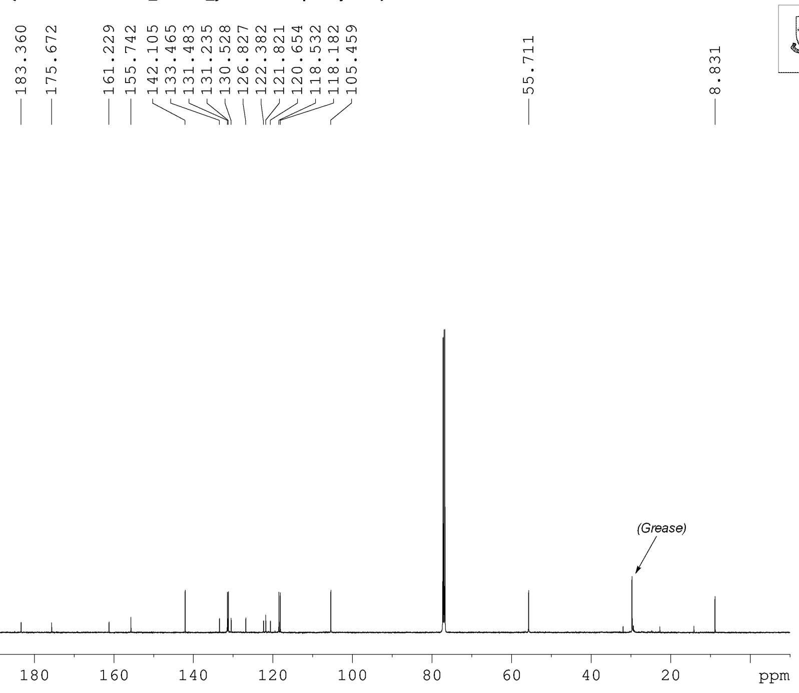


The University Of Sheffield.

```
Current Data Parameters
NAME             R=OMe 13C
EXPNO                   2
PROCNO                  1

F2 - Acquisition Parameters
Date_            20170618
Time                 0.01
INSTRUM             spect
PROBHD   5 mm PABBO BB/
PULPROG             udeft
TD                  21424
SOLVENT             CDCl3
NS                  20480
DS                      0
SWH             29761.904 Hz
FIDRES           1.389185 Hz
AQ              0.3599232 sec
RG                  189.1
DW                 16.800 usec
DE                   8.90 usec
TE                  298.0 K
D1             4.00000000 sec
D11            0.03000000 sec
D12            0.00002000 sec
D20          200.00000000 sec
TD0                    20

======== CHANNEL f1 ========
SFO1          125.7703656 MHz
NUC1                  13C
P1                   9.00 usec
P13               2000.00 usec
P26                500.00 usec
PLW1          95.00000000 W
SPNAM[5]     Crp60comp.4
SPOAL5              0.500
SPOFFS5  0 Hz
SPW5          11.75699997 W
SPNAM[8] Crp60,0.5,20.1
SPOAL8              0.500
SPOFFS8  0 Hz
SPW8          11.75699997 W

======== CHANNEL f2 ========
SFO2          500.1320007 MHz
NUC2                   1H
CPDPRG[2          waltz16
PCPD2               80.00 usec
PLW2          22.00000000 W
PLW12          0.34375000 W

F2 - Processing parameters
SI                 131072
SF            125.7577885 MHz
WDW                    EM
SSB      0
LB                   2.00 Hz
GB       0
PC                   1.40
```

**MJF-2-44-P1B**
**PRO CDCl3 {C:\NMRData\Jones_current_year} mdp14mjf 57**

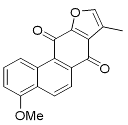


**40**

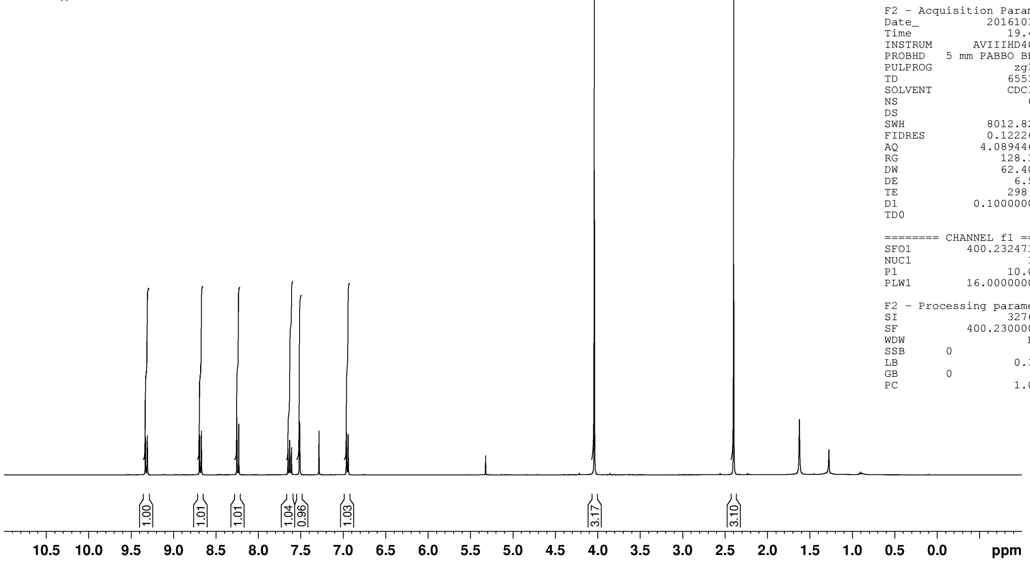


The University Of Sheffield.

```
Current Data Parameters
NAME              R=OMe 1H
EXPNO                    2
PROCNO                   1

F2 - Acquisition Parameters
Date_             20161010
Time                 19.42
INSTRUM         AVIIIHD400
PROBHD   5 mm PABBO BB/
PULPROG               zg30
TD                   65536
SOLVENT              CDCl3
NS                      64
DS                       2
SWH              8012.820 Hz
FIDRES           0.122266 Hz
AQ              4.0894465 sec
RG                  128.39
DW                  62.400 usec
DE                    6.50 usec
TE                   298.0 K
D1              0.10000000 sec
TD0                      1

======== CHANNEL f1 ========
SFO1          400.2324716 MHz
NUC1                    1H
P1                   10.00 usec
PLW1           16.00000000 W

F2 - Processing parameters
SI                   32768
SF            400.2300000 MHz
WDW                     EM
SSB      0
LB                    0.30 Hz
GB       0
PC                    1.00
```

**Matt Foulkes, mdp14mjf, Jones Gp**
**Sample Ref: MJF-2-44-P1B**

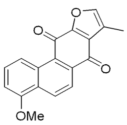


**40**

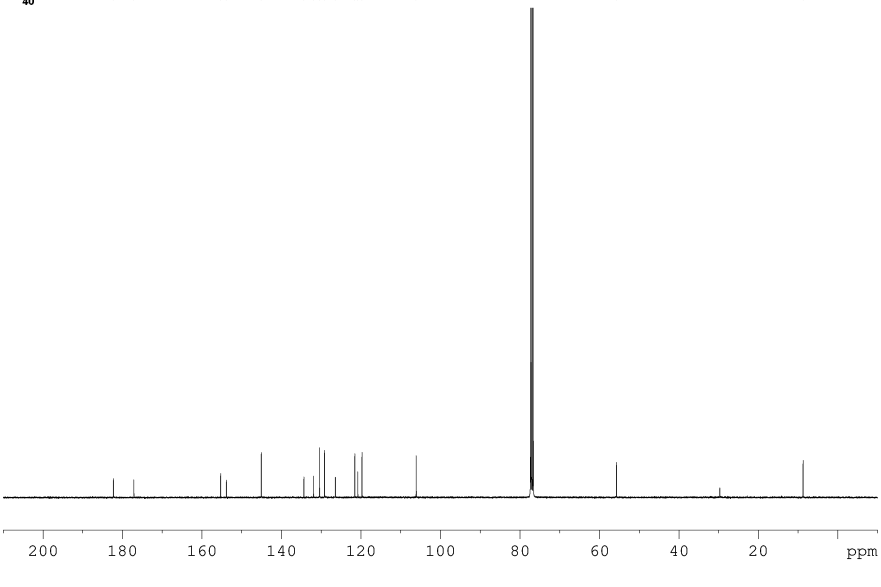


The University Of Sheffield.

```
Current Data Parameters
NAME             R=OMe 13C
EXPNO                    1
PROCNO                   1

F2 - Acquisition Parameters
Date_             20180309
Time                 12.01
INSTRUM              spect
PROBHD   5 mm PABBO BB-
PULPROG             zgpg30
TD                   65536
SOLVENT              CDCl3
NS                    3946
DS                       4
SWH              25252.525 Hz
FIDRES            0.385323 Hz
AQ               1.2976128 sec
RG                     362
DW                  19.800 usec
DE                    6.67 usec
TE                   298.0 K
D1              5.00000000 sec
D11             0.03000000 sec
TD0                      5

======== CHANNEL f1 ========
NUC1                   13C
P1                    8.10 usec
PL1                  -1.80 dB
PL1W           52.46613693 W
SFO1           100.6419425 MHz

======== CHANNEL f2 ========
CPDPRG[2           waltz16
NUC2                    1H
PCPD2                90.00 usec
PL2                  -2.70 dB
PL12                 16.83 dB
PL13                 16.90 dB
PL2W           18.38199043 W
PL12W           0.20482953 W
PL13W           0.20155454 W
SFO2           400.2016008 MHz

F2 - Processing parameters
SI                   32768
SF             100.6303712 MHz
WDW                     EM
SSB      0
LB                    1.00 Hz
GB       0
PC                    1.40
```

**MJF–2–56– FR1–3**
**PRO CDCl3 {C:\NMRData\Jones_current_year} mdp14mjf 21**

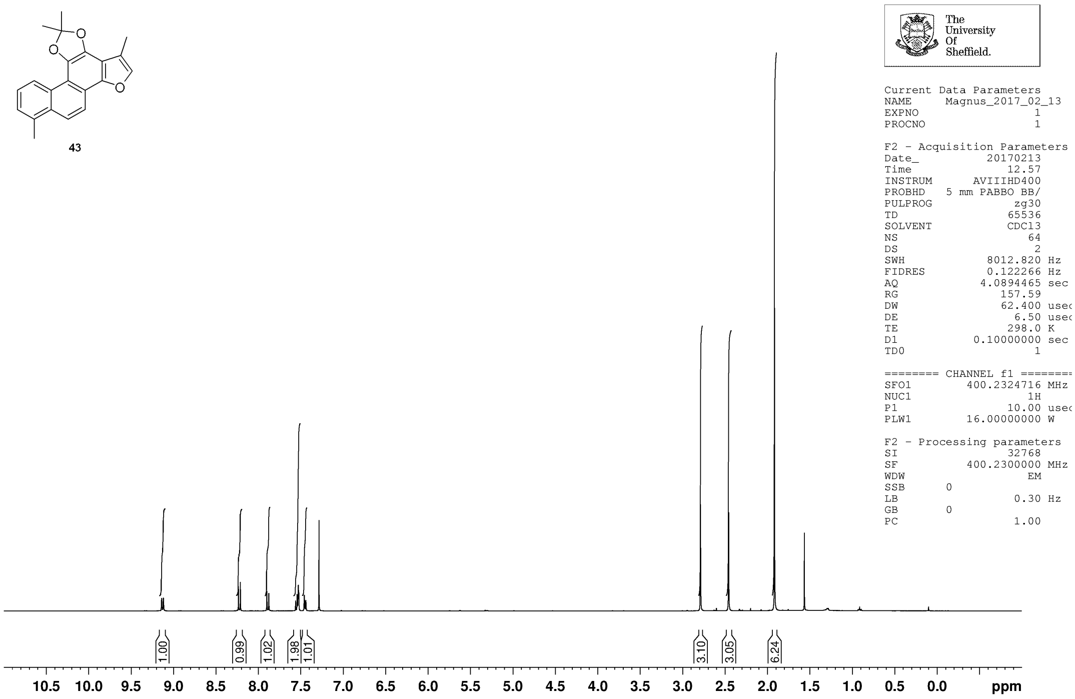


**Matt Foulkes, mdp14mjf, Jones Group**
**Sample Ref: MJF–2–56–Fr.1–3**

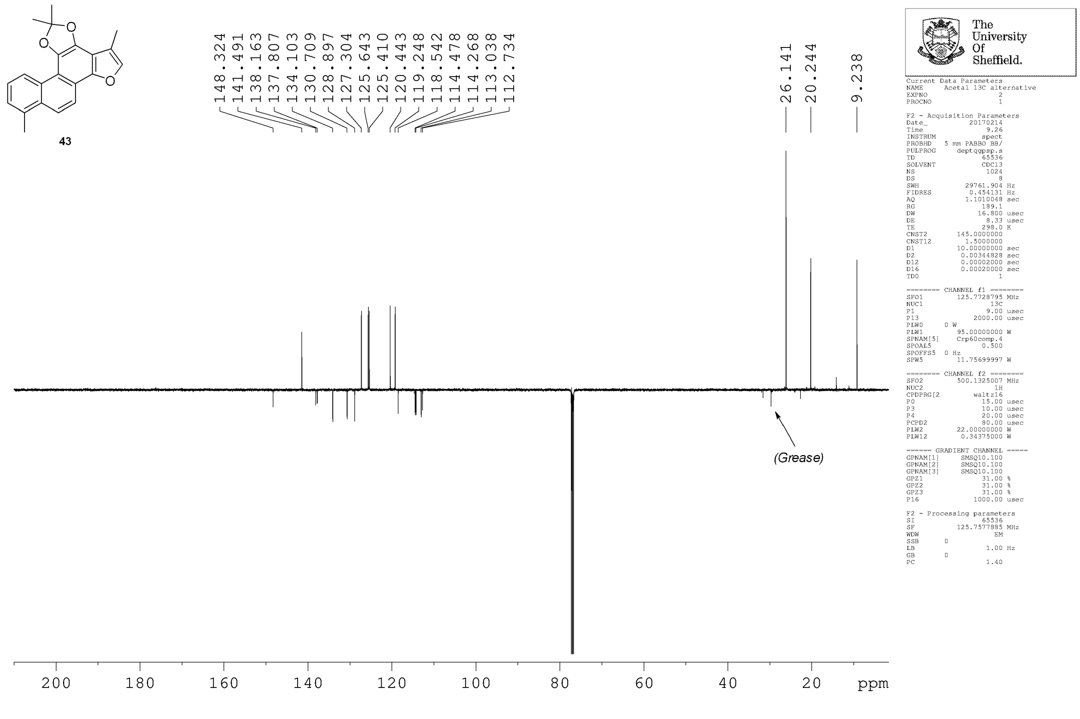

**MJF-2-75-P2**
**PROTON.s CDCl3 {C:\NMRData\Jones_current_year} mdp14mjf 9**

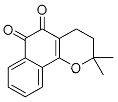

5

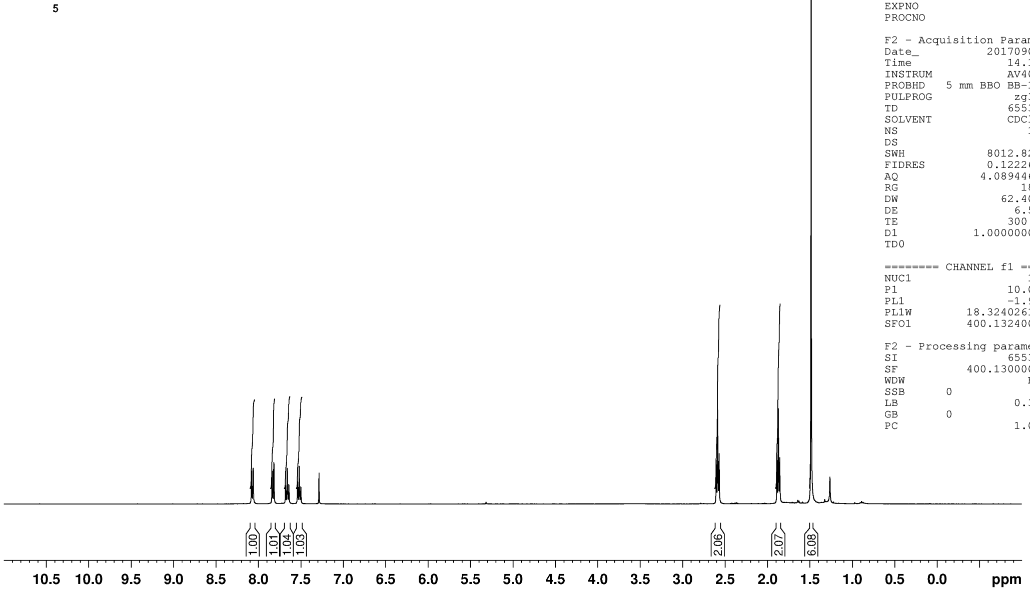


The University Of Sheffield.

```
Current Data Parameters
NAME      B-lapachone 1H
EXPNO                  1
PROCNO                 1

F2 - Acquisition Parameters
Date_           20170901
Time               14.10
INSTRUM            AV400
PROBHD   5 mm BBO BB-1H
PULPROG             zg30
TD                 65536
SOLVENT            CDCl3
NS                    16
DS                     0
SWH           8012.820 Hz
FIDRES        0.122266 Hz
AQ           4.0894465 sec
RG                   181
DW              62.400 usec
DE                6.50 usec
TE               300.0 K
D1          1.00000000 sec
TD0                    1

======== CHANNEL f1 ========
NUC1                  1H
P1               10.00 usec
PL1              -1.90 dB
PL1W       18.32402611 W
SFO1       400.1324008 MHz

F2 - Processing parameters
SI                 65536
SF         400.1300000 MHz
WDW                   EM
SSB      0
LB                0.30 Hz
GB       0
PC                1.00
```

**MJF-2-75-P2**
**DEPTQ.l CDCl3 {C:\NMRData\Jones_current_year} mdp14mjf 9**

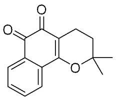

5

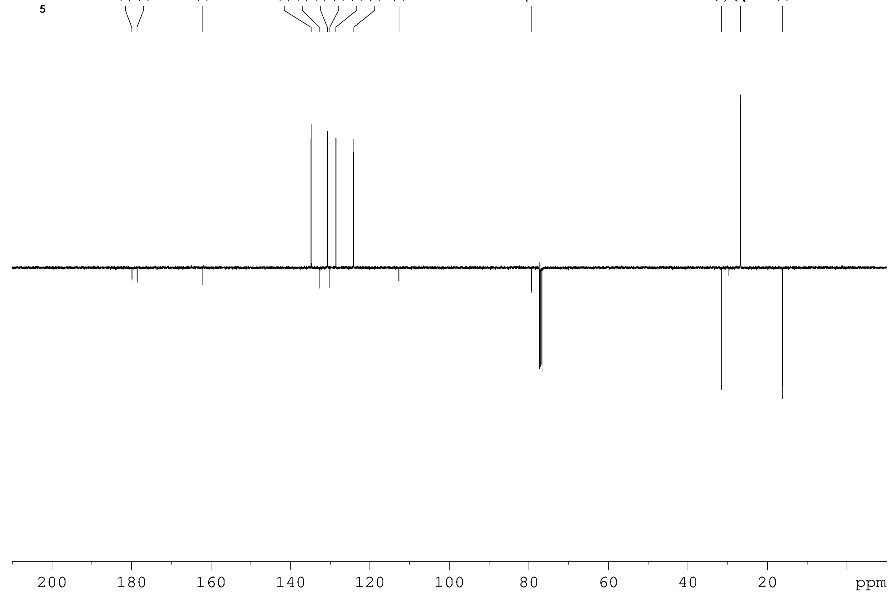


The University Of Sheffield.

```
Current Data Parameters
NAME      B-lapachone 13C
EXPNO                  2
PROCNO                 1

F2 - Acquisition Parameters
Date_           20170901
Time               20.44
INSTRUM            AV400
PROBHD   5 mm BBO BB-1H
PULPROG          deptqsp
TD                 65536
SOLVENT            CDCl3
NS                  1024
DS                     4
SWH          35211.270 Hz
FIDRES        0.537281 Hz
AQ           0.9306112 sec
RG                 16384
DW              14.200 usec
DE                6.50 usec
TE               300.0 K
CNST2      145.0000000
CNST12       1.5000000
D1          4.00000000 sec
D2          0.00344828 sec
D12         0.00002000 sec
TD0                    1
ZGOPTNS

======== CHANNEL f1 ========
NUC1                 13C
P1                8.00 usec
P12            2000.00 usec
PL0             120.00 dB
PL1              -3.10 dB
PL0W     0 W
PL1W       58.97905731 W
SFO1       100.6293701 MHz
SP2               7.00 dB
SPNAM[2]    Crp60comp.4
SPOAL2             0.500
SPOFFS2  0 Hz

======== CHANNEL f2 ========
CPDPRG[2         waltz16
NUC2                  1H
P0               15.00 usec
P3               10.00 usec
P4               20.00 usec
PCPD2            88.00 usec
PL2              -1.90 dB
PL12             16.99 dB
PL2W       18.32402611 W
PL12W       0.23660338 W
SFO2       400.1316005 MHz

F2 - Processing parameters
SI                 65536
SF         100.6127690 MHz
WDW                   EM
SSB      0
LB                1.00 Hz
GB       0
PC                1.40
```

**MJF-2-76-P2A**
**PROTON.s CDCl3 {C:\NMRData\Jones_current_year} mdp14mjf 7**

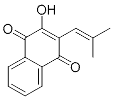


**45**

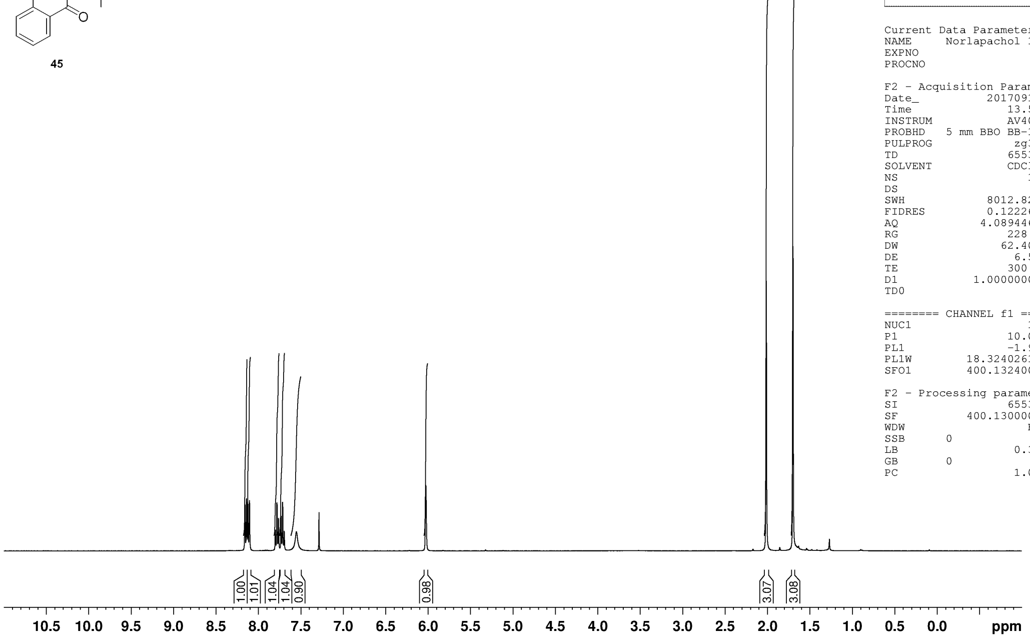


The University Of Sheffield.

```
Current Data Parameters
NAME        Norlapachol 1H
EXPNO                    2
PROCNO                   1

F2 - Acquisition Parameters
Date_             20170912
Time                 13.56
INSTRUM              AV400
PROBHD    5 mm BBO BB-1H
PULPROG               zg30
TD                   65536
SOLVENT              CDCl3
NS                      16
DS                       0
SWH               8012.820 Hz
FIDRES            0.122266 Hz
AQ               4.0894465 sec
RG                   228.1
DW                  62.400 usec
DE                    6.50 usec
TE                   300.0 K
D1              1.00000000 sec
TD0                      1

======== CHANNEL f1 ========
NUC1                    1H
P1                   10.00 usec
PL1                  -1.90 dB
PL1W           18.32402611 W
SFO1           400.1324008 MHz

F2 - Processing parameters
SI                   65536
SF             400.1300000 MHz
WDW                     EM
SSB      0
LB                    0.30 Hz
GB       0
PC                    1.00
```

**MJF-2-76-P2A**
**DEPTQ.s CDCl3 {C:\NMRData\Jones_current_year} mdp14mjf 13**

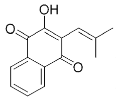


**45**

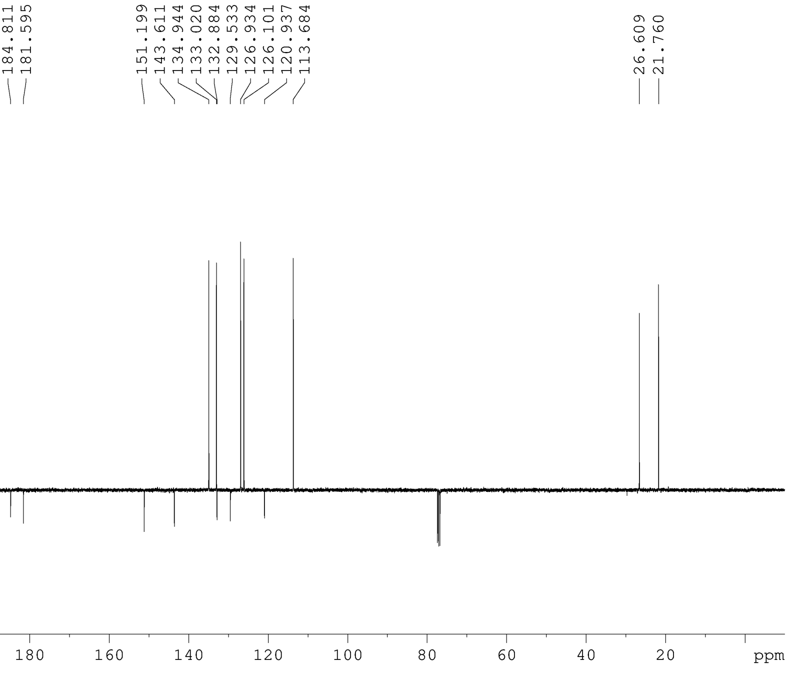


The University Of Sheffield.

```
Current Data Parameters
NAME       Norlapachol 13C
EXPNO                    3
PROCNO                   1

F2 - Acquisition Parameters
Date_             20170912
Time                 15.11
INSTRUM              AV400
PROBHD    5 mm BBO BB-1H
PULPROG            deptqsp
TD                   65536
SOLVENT              CDCl3
NS                     192
DS                       4
SWH              35211.270 Hz
FIDRES            0.537281 Hz
AQ               0.9306112 sec
RG                   16384
DW                  14.200 usec
DE                    6.50 usec
TE                   300.0 K
CNST2          145.0000000
CNST12           1.5000000
D1              4.00000000 sec
D2              0.00344828 sec
D12             0.00002000 sec
TD0                      1
ZGOPTNS

======== CHANNEL f1 ========
NUC1                   13C
P1                    8.00 usec
P12                2000.00 usec
PL0                 120.00 dB
PL1                  -3.10 dB
PL0W      0 W
PL1W           58.97905731 W
SFO1           100.6293701 MHz
SP2                   7.00 dB
SPNAM[2]       Crp60comp.4
SPOAL2               0.500
SPOFFS2   0 Hz

======== CHANNEL f2 ========
CPDPRG[2          waltz16
NUC2                    1H
P0                   15.00 usec
P3                   10.00 usec
P4                   20.00 usec
PCPD2                88.00 usec
PL2                  -1.90 dB
PL12                 16.99 dB
PL2W           18.32402611 W
PL12W           0.23660338 W
SFO2           400.1316005 MHz

F2 - Processing parameters
SI                   65536
SF             100.6127690 MHz
WDW                     EM
SSB      0
LB                    1.00 Hz
GB       0
PC                    1.40
```

**MJF-2-77-P1A**
**PROTON.s CDCl3 {C:\NMRData\Jones_current_year} mdp14mjf 18**

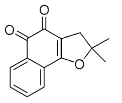

6

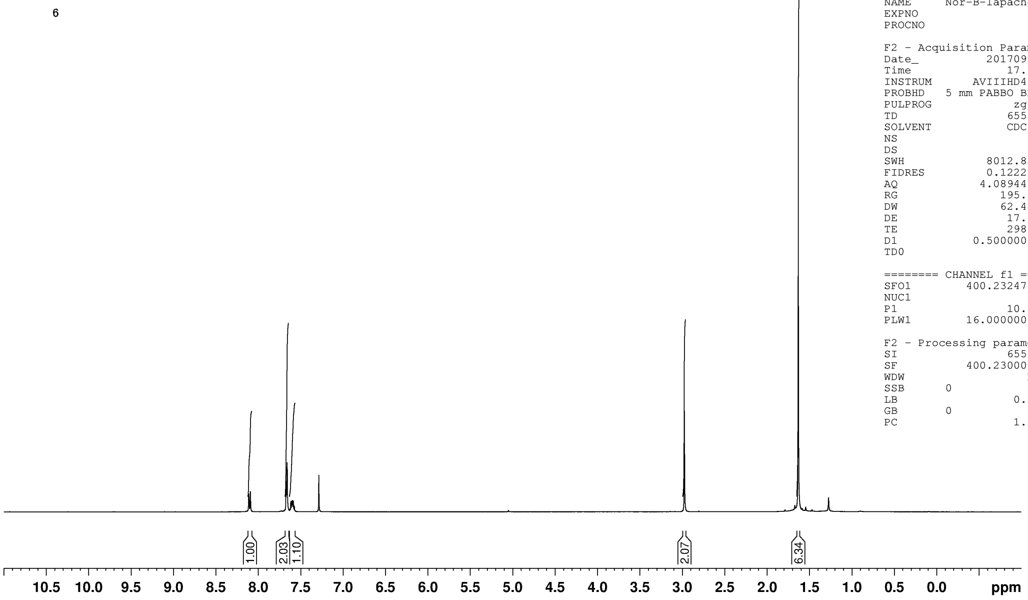


The University Of Sheffield.

```
Current Data Parameters
NAME       Nor-B-lapachone 1H
EXPNO                    1
PROCNO                   1

F2 - Acquisition Parameters
Date_             20170915
Time                 17.32
INSTRUM         AVIIIHD400
PROBHD     5 mm PABBO BB/
PULPROG               zg30
TD                   65536
SOLVENT              CDCl3
NS                      16
DS                       2
SWH              8012.820 Hz
FIDRES           0.122266 Hz
AQ              4.0894465 sec
RG                  195.75
DW                  62.400 usec
DE                   17.37 usec
TE                   298.0 K
D1             0.50000000 sec
TD0                      1

======== CHANNEL f1 ========
SFO1          400.2324716 MHz
NUC1                    1H
P1                   10.00 usec
PLW1          16.00000000 W

F2 - Processing parameters
SI                   65536
SF            400.2300000 MHz
WDW                     EM
SSB      0
LB                    0.30 Hz
GB       0
PC                    1.00
```

**MJF-2-77-P1**
**DEPTQ.I CDCl3 {C:\NMRData\Jones_current_year} mdp14mjf 30**

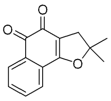

6

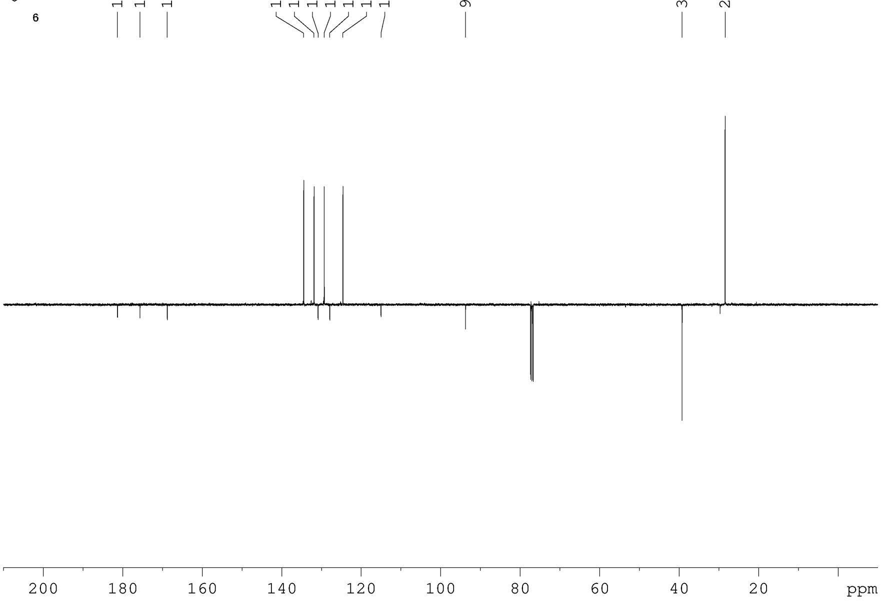


The University Of Sheffield.

```
Current Data Parameters
NAME       Nor-B-lapachone 13C
EXPNO                    2
PROCNO                   1

F2 - Acquisition Parameters
Date_             20170914
Time                 22.10
INSTRUM              AV400
PROBHD     5 mm BBO BB-1H
PULPROG            deptqsp
TD                   65536
SOLVENT              CDCl3
NS                    1024
DS                       4
SWH             35211.270 Hz
FIDRES           0.537281 Hz
AQ              0.9306112 sec
RG                   16384
DW                  14.200 usec
DE                    6.50 usec
TE                   300.0 K
CNST2         145.0000000
CNST12          1.5000000
D1             4.00000000 sec
D2             0.00344828 sec
D12            0.00002000 sec
TD0                      1
ZGOPTNS

======== CHANNEL f1 ========
NUC1                   13C
P1                    8.00 usec
P12                2000.00 usec
PL0                 120.00 dB
PL1                  -3.10 dB
PL0W     0 W
PL1W          58.97905731 W
SFO1          100.6293701 MHz
SP2                   7.00 dB
SPNAM[2]      Crp60comp.4
SPOAL2               0.500
SPOFFS2  0 Hz

======== CHANNEL f2 ========
CPDPRG[2          waltz16
NUC2                    1H
P0                   15.00 usec
P3                   10.00 usec
P4                   20.00 usec
PCPD2                88.00 usec
PL2                  -1.90 dB
PL12                 16.99 dB
PL2W          18.32402611 W
PL12W          0.23660338 W
SFO2          400.1316005 MHz

F2 - Processing parameters
SI                   65536
SF            100.6127690 MHz
WDW                     EM
SSB      0
LB                    1.00 Hz
GB       0
PC                    1.40
```
